# 3D genome topologies distinguish pluripotent epiblast and primitive endoderm cells in the mouse blastocyst

**DOI:** 10.1101/2022.10.19.512781

**Authors:** Gesa Loof, Dominik Szabó, Vidur Garg, Alexander Kukalev, Luna Zea-Redondo, Rieke Kempfer, Thomas M. Sparks, Yingnan Zhang, Christoph J Thieme, Sílvia Carvalho, Anja Weise, Milash Balachandran, Thomas Liehr, Lonnie R. Welch, Anna-Katerina Hadjantonakis, Ana Pombo

## Abstract

The development of embryonic cell lineages is tightly controlled by transcription factors that regulate gene expression and chromatin organisation. To investigate the specialisation of 3D genome structure in pluripotent or extra-embryonic endoderm lineages, we applied Genome Architecture Mapping (GAM) in embryonic stem (ES) cells, extra-embryonic endoderm (XEN) stem cells, and in their *in vivo* counterparts, the epiblast (Epi) and primitive endoderm (PrE) cells, respectively. We discover extensive differences in 3D genome topology including the formation domain boundaries that differ between Epi and PrE lineages, both *in vivo* and *in vitro*, at lineage commitment genes. In ES cells, *Sox2* contacts other active regions enriched for NANOG and SOX2 binding sites. PrE-specific genes, such as *Lama1* and *Gata6*, form repressive chromatin hubs in ES cells. *Lama1* activation in XEN or PrE cells coincides with its extensive decondensation. Putative binding sites for OCT4 and SNAIL, or GATA4/6, distinguish chromatin contacts unique to embryonic or extra-embryonic lineages, respectively. Overall, 3D genome folding is highly specialised in early development, especially at genes encoding factors driving lineage identity.

**Highlights:**

- ES and XEN cells have specialised 3D genome structures
- GAM applied in the blastocyst distinguishes Epi and PrE genome structures
- Lineage specific genes establish cell-type specific chromatin contacts
- Specific chromatin contacts feature putative bindings sites for GATA4/6 in XEN cells and SNAIL in ES cells

## Introduction

Mammalian pre-implantation development is a tightly regulated process, which culminates in the formation of a blastocyst stage embryo that implants into the maternal uterine wall. Within the first four and a half days of murine embryonic development, the embryo acquires cells of three distinct lineages, through two binary sequential lineage decisions, and for the first time establishes a population of pluripotent cells, as founders of the embryo proper. In the first lineage decision, which initiates at the 8-to-16-cell stage, the cells of the embryo become compacted and polarised and give rise to an outer cell population of trophectoderm (TE) and the inner cell mass (ICM) cells (Chazaud and Yamanaka, 2016). The ICM subsequently segregates into the primitive endoderm (PrE), which will predominantly contribute to extra-embryonic support structures (Gardner and Rossant, 1979), and the pluripotent epiblast (Epi) which gives rise to most of the cells of the developing organism. The ICM lineage decision is coordinated by signalling cascades chiefly driven by FGF/ERK, and orchestrated by transcription factors such as NANOG, SOX2, OCT4, GATA6, GATA4 and SOX17 (Kang et al., 2017; Kang et al., 2013; Saiz et al., 2016), where GATA6 and NANOG repress one another, while positively regulating their own expression (Bessonnard et al., 2014; Frankenberg et al., 2011; Schrode et al., 2014). Furthermore, the two ICM-derived lineages of the Epi and PrE can be captured ex vivo as self-renewing stem cell models in embryonic stem (ES) cells and extra-embryonic stem (XEN) cells, respectively (Garg et al., 2016; Watts et al., 2018).

Extensive differences in gene expression and local chromatin regulation have been reported in ES and XEN cells, and during blastocyst development (Choi et al., 2020; Santos et al., 2010; Wamaitha et al., 2015; Wu et al., 2016; Zheng and Xie, 2019). However, less is known about the extent of specialisation of the 3D genome structure during these early stages of embryo lineage commitment. For example, topologically associating domains (TADs) are first established during pre-implantation development within the ICM, concurrent with the onset of pluripotency (Du et al., 2017; Ke et al., 2017), but it remains unclear whether TAD organisation is cell-type specialised in the Epi and PrE lineages. Moreover, genomic regions separated by several megabases and enriched for binding of SOX2, OCT4 and NANOG, drivers of Epi development, have been found to contact each other in ES cells (Beagrie et al., 2017; de Wit et al., 2013; Quinodoz et al., 2018). In XEN cells, binding of GATA6 is known to occur at the promoters of extra-embryonic endoderm genes including *Gata4/6, Sox7/17* and *Pdgfra*, as well as pluripotency-associated genes, such as *Nanog, Esrrb* and *Pou5f1* (Wamaitha et al., 2015). However, the extent of cell-type specialisation of 3D genome structure in Epi and PrE cells, which develop from a common ICM precursor in the early embryo, remains to be studied.

To investigate whether lineage commitment in the early embryo coincides with a cell-type specification of 3D genome topologies, we applied Genome Architecture Mapping (GAM) to cells of the embryo and their *in vitro* lineage counterparts. We analysed the Epi and PrE cells of E4.5 blastocyst stage embryos, as well as their counterparts ES and XEN stem cells (**Figure 1**). GAM is a ligation-free technology that maps chromatin contacts by sequencing the DNA content from thin (~220 nm) nuclear cryosections collected in random orientations (Beagrie et al., 2017). Chromatin contacts are inferred from the probability of co-segregation of genomic loci across the collection of nuclear slices. GAM’s recent extension, immunoGAM, is ideally suited for studying cells directly in the embryo, as it enables mapping of 3D chromatin topology in specific cells within tissues, without dissociation, by immunolabelling the thin GAM cryosections with cell-type specific markers prior to microdissection (Winick-Ng et al., 2021). To investigate how lineage-specific 3D chromatin conformation relates with gene regulation, we also collected bulk RNA-seq and ATAC-seq datasets from ES and XEN cells, and mined single-cell RNA-seq data from early embryos that we had previously collected (Nowotschin et al., 2019).

**Figure 1.**
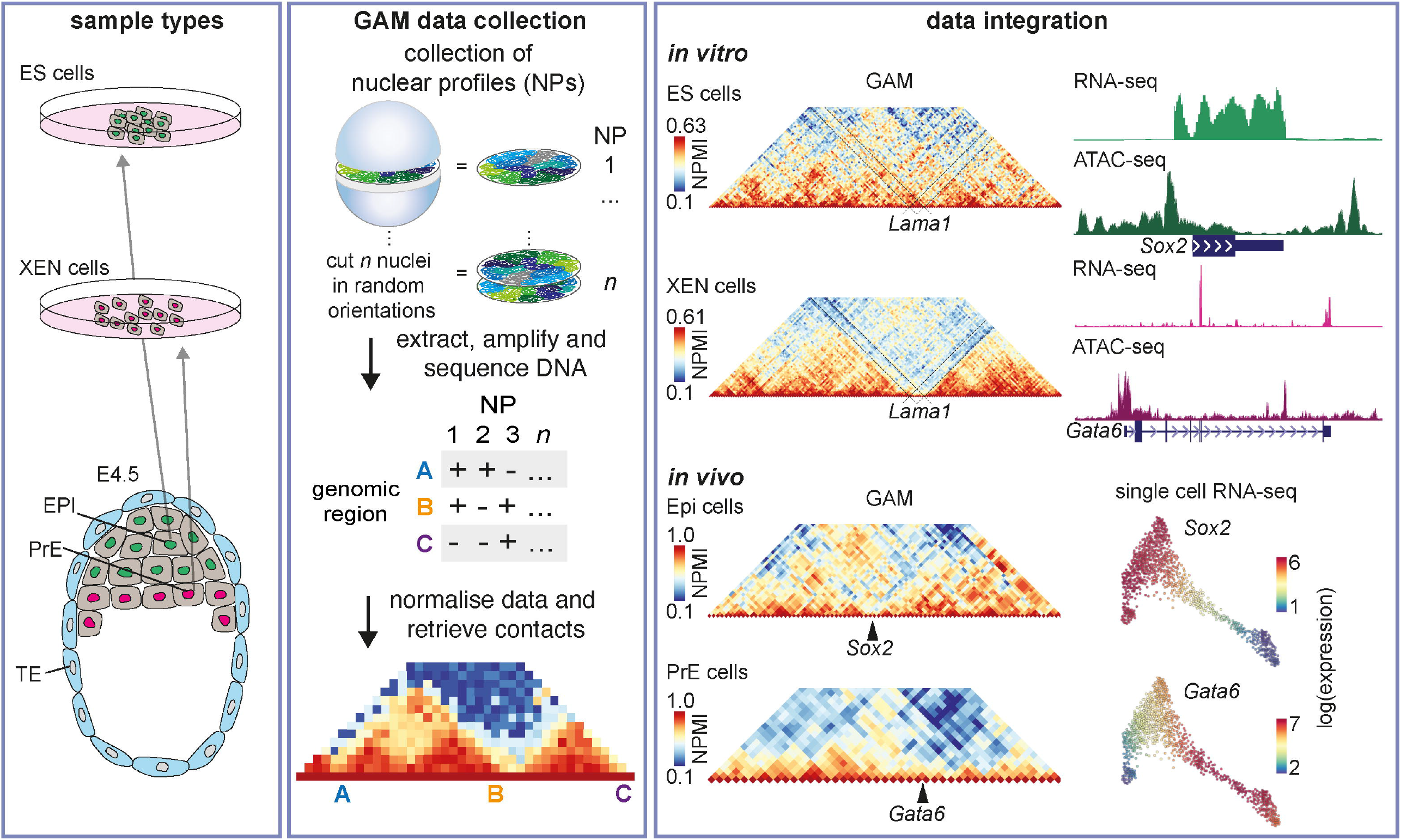
Overview of data collection. GAM data was collected from ES and XEN cells and the respective lineages in the E4.5 embryo, the Epiblast (Epi) and primitive endoderm (PrE). GAM data was integrated with RNA-seq and ATAC-seq data to study genome topology and its role in cell-type identity.

Our data reveal extensive differences in 3D chromatin conformation between the Epi and PrE lineages, including chromatin contacts that contain binding sites specific for GATA4 and GATA6 in XEN cells, or OCT4 and SNAIL in ES cells. We also uncover the extensive decondensation of *Lama1* in XEN cells, accompanied by compartment transitions and formation of a new TAD boundary. *Lama1* encodes for Laminin alpha 1, a protein which is expressed by extra-embryonic endoderm cells and essential for proper embryo development (Ueda et al., 2020), and is amongst the top 1% most upregulated genes in XEN cells. To expand from our experiments in stem cell models, we applied GAM to the PrE and Epi cells of E4.5 mouse embryos, and found that the cell-type specialisation of the 3D genome organisation observed *in vitro* also occurs *in vivo*, with extensive structural differences present at key lineage-associated loci.

## Results

### Mapping chromatin contacts genome-wide in ES and XEN cells

ES and XEN cells are *in vitro* stem cell models representing the epiblast and primitive endoderm lineages of the early mammalian embryo, and are characterised by the expression of specific developmental regulator transcription factors (TF), such as *Nanog* and *Gata6*, respectively (**Figure 2A**). To investigate any differences in genome architecture between ES and XEN cells, we took our previously published GAM dataset from ES cells (Beagrie et al., 2021), and generated additional GAM data for XEN cells (**Figure 2B**). XEN cell GAM datasets were collected in multiplex-GAM mode, a GAM approach that pools together three independent nuclear slices in each library, and produces contact matrices which are comparable to single-cell GAM collection mode, as shown previously for ES cells (Beagrie et al., 2021). Final GAM datasets consisted of a total of 765 ES cells and 1896 XEN cells, and showed good detectability of all possible pairs of intra-chromosomal genomic windows at 20-kb resolution (> 93%; **Figure S1A**).

**Figure 2.**
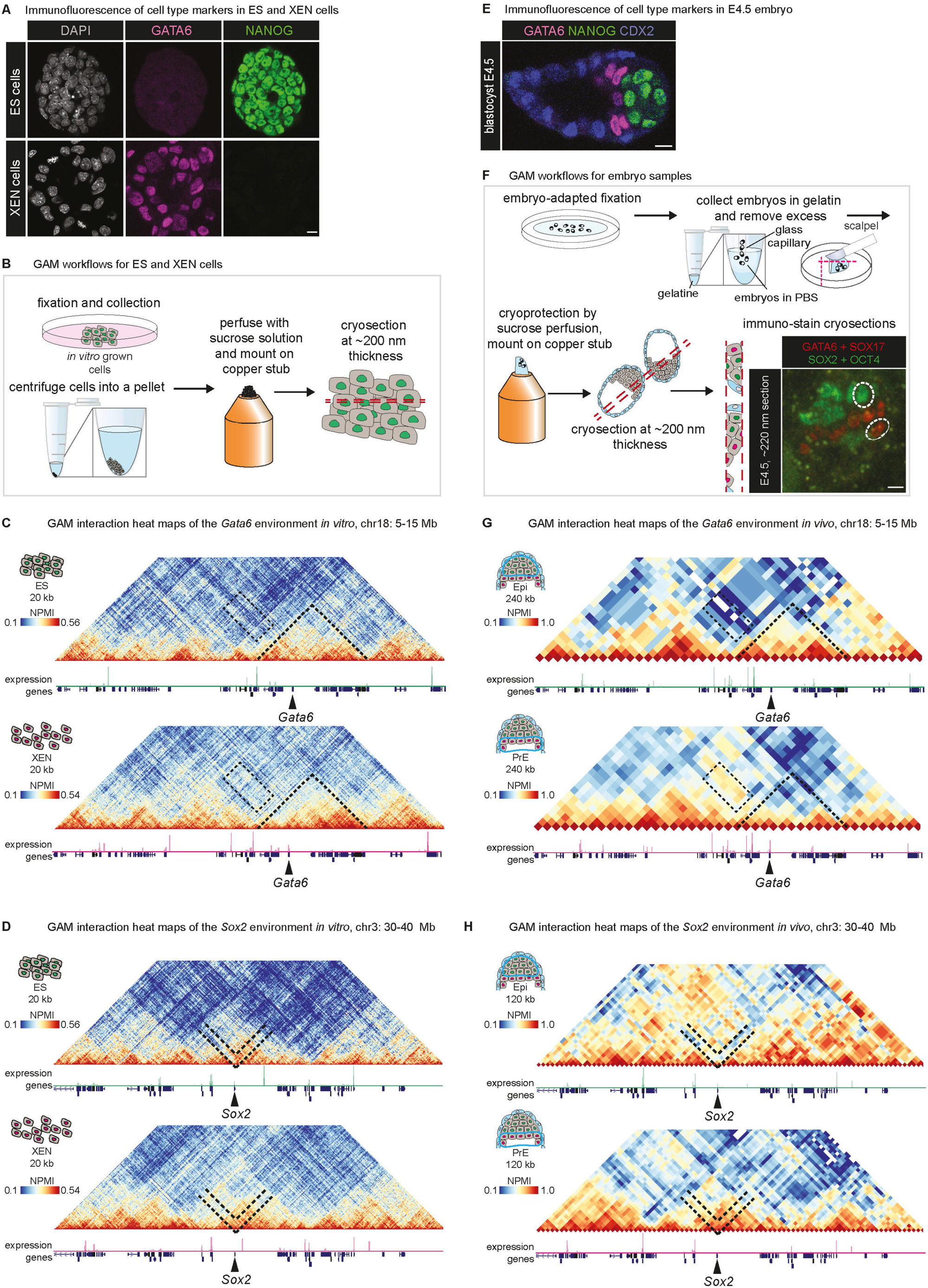
Genome Architecture mapping in *in vitro* and *in vivo* pre-implantation development shows topological changes of key lineage driving gene loci. A. Immunofluorescence of GATA6 and NANOG in ES and XEN cells. Scale bar corresponds to 10 µm. B. GAM workflow applied to tissue culture samples. The sample processing involves fixation of cells while attached to the tissue culture dish, and subsequently forming a cell pellet from scraped-off cells by centrifugation, followed by mounting of the sucrose-cryoprotected cell pellet on a copper stub and freezing in liquid nitrogen for cryosectioning. C. Interaction heat maps show NPMI-normalised GAM data for the region chr18:5-15 Mb, centred around the *Gata6* gene, at 20 kb resolution in ES and XEN cells. Underneath the GAM matrices, RPKM-normalised RNA-seq tracks show gene expression. D. Same as (C), for region chr3:30-40 Mb, centred around the *Sox2* gene. E. Immunofluorescence of GATA6, NANOG and CDX2 in an E4.5 embryo. Scale bar corresponds to 10 µm. F. Workflow of GAM sample preparation for E4.5 embryos, including embryo-adapted fixation and gelatin embedding of up to 15 embryos per sample. Excess gelatin is removed and sucrose-cryoprotected samples are frozen on copper stubs for cryosectioning. Cryosections are immunostained before the collection of nuclear profiles. Scale bar corresponds to 10 µm. G. Interaction heat maps show NPMI-normalised GAM data for the region chr18:5-15 Mb, centred around the *Gata6* gene, at 120 kb resolution in Epi and PrE cells. Underneath the GAM matrices, pseudo-bulk scRNA-seq tracks show gene expression. H. Same as (E), for region chr3:30-40 Mb, centred around the *Sox2* gene.

GAM matrices from ES and XEN cells showed visible differences in chromatin contacts. For example, the activation of *Gata6* in XEN cells coincided with the decondensation of its surrounding genomic region (**Figure 2C**, dotted lines), while the activation of *Sox2* in ES cells resulted in the loss of local interactions with its genomic neighbourhood compared with XEN cells (**Figure 2D**, dotted lines). These changes in genome architecture revealed a connection of DNA folding to cell identity in early embryonic development.

### Mapping chromatin contacts in Epi and PrE cells of the embryo

*Nanog* and *Gata6* are also specifically expressed in Epi and PrE cells, respectively, in the E4.5 mouse embryo (**Figure 2E**). To map 3D genome conformation in Epi and PrE cells directly within the embryo, we developed a novel sample preparation protocol for performing GAM in early mouse embryos that preserves their morphology during fixation, freezing and ultrathin cryosectioning, and avoids aggressive dissociation of the embryo (**Figure 2F**). E4.5 embryos were collected, fixed with electron-microscopy grade formaldehyde, and embedded in gelatin for cryosectioning. Confocal imaging of SOX17, GATA6, SOX2 and OCT4 immunostained 220 nm-thick cryosections showed intact embryo and nuclear morphology and enabled high resolution imaging and lineage assignment in embryos (**Figure S1B**). We collected 70 and 126 single nuclear slices from Epi and PrE cells, respectively (**Figure S1C**). The GAM data collected was of high quality, with 59 Epi and 111 PrE GAM samples passing quality control checks (84% and 88%, respectively; see Methods), at rates similar to previous reports (Winick-Ng et al., 2021).

To evaluate the quality of the GAM datasets collected from Epi and PrE cells of blastocyst stage embryos, we plotted the pairwise contact matrices centred on *Gata6* and *Sox2*, using genomic resolutions of 120 or 240 kb, respectively. We found local 3D genome architectures of key lineage-associated loci to be similar to those in ES and XEN cells, such as the local decondensation of the expressed *Gata6* gene in PrE cells, with increased detection of contacts upstream connecting *Gata6* to other genes and an intergenic region (**Figure 2G**, dashed rectangle). Loss of local contacts of the active *Sox2* locus was found in Epi cells, to an even larger extent than observed in ES cells, in contrast with marked chromatin condensation of the locus in both XEN and PrE cells where *Sox2* is silent (**Figure 2H**). Further lineage differences between ES/Epi cells or XEN/PrE cells could be observed for other lineage markers, such as *Sox17* (extra-embryonic endoderm specific), *Pou5f1* and *Nanog* (pluripotency associated; **Figure S1D-F**). The robustness and working resolution of the smaller GAM datasets collected from Epi and PrE cells was assessed in detail (see Methods), which confirmed their good sampling quality at 120 kb resolution, and showed that they captured general features of local chromatin contacts. To exercise caution, these GAM datasets were used here exclusively for empirical comparisons with the *in vitro* GAM datasets from ES and XEN cells, without further quantification at the genome-wide level.

### ES and XEN cells have highly differential gene expression programs

To interpret how differences in genome topology relate to changes in gene expression between ES and XEN cells, we collected RNA-seq data in five and three biological replicates, respectively. Single gene tracks showed gene expression at lineage-specific genes, *Sox2* and *Gata6* (**Figure S2A**), and principal component analysis showed high variance between ES and XEN expression by clustering of datasets according to cell type (**Figure S2B**).

First, we compared the expression patterns of lineage-specific marker genes in ES and XEN cell bulk RNA-seq data and previously published single-cell RNA-seq data collected from E4.5 embryos (Nowotschin et al., 2019; **Figure 3A**). We found comparable expression patterns between *in vitro* and *in vivo* cells, such as the expression of *Pou5f1, Sox2, Fgf4* in both ES and Epi cells, the expression of *Gata6, Gata4, Sox17* and *Pdgfra* in both XEN and PrE cells, with little detectable expression of TE markers *Cdx2* and *Eomes* in either cell line. We also observed known differences, such as lower detection of *Nanog* transcript in Epi in comparison to ES cells, and intermediate expression of *Pou5f1* in PrE relative to XEN cells (Kunath et al., 2005; Morgani et al., 2017).

**Figure 3.**
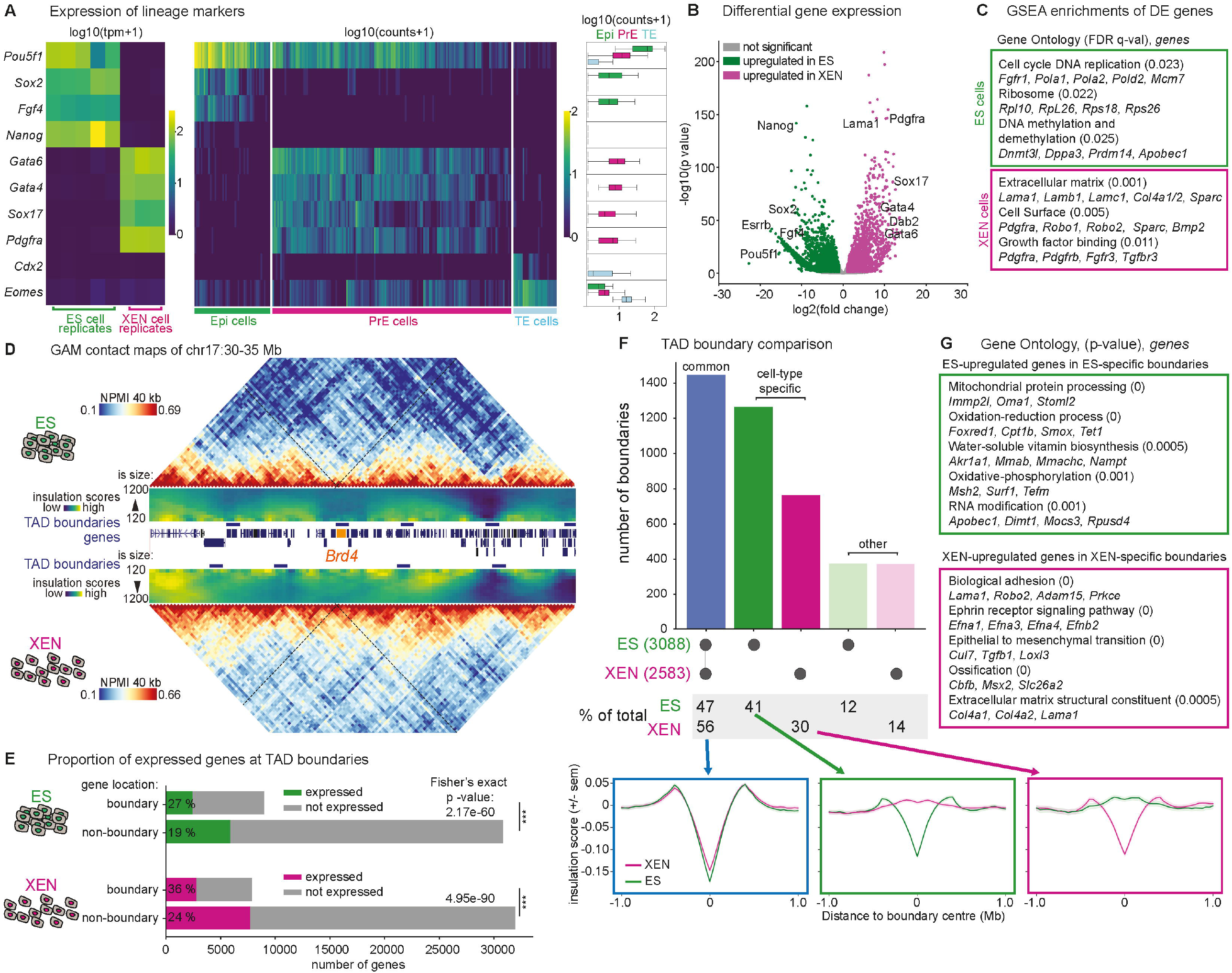
Changes in transcriptional profiles are connected to large-scale changes in genome folding. A. Heat maps show expression of lineage marker genes in total RNA-seq of ES and XEN cells and in scRNA-seq of cells classified as Epi, PrE and TE in Nowotschin et al. (2019). TPM and count values are log10-scaled with a pseudo-count of 1 added to avoid negative values. Boxplots show distribution of counts in scRNA-seq data across all Epi, PrE and TE cells. B. Differential gene expression analysis between ES and XEN cells shows differential expression of lineage driving genes. 7421 genes were called differentially expressed, using the cut-offs log2(fold changes) > 1 and adjusted p-value < 0.05 (marked in green and magenta). C. GSEA enrichments were calculated for genes ranked by their p-values of differential expression, and split by the sign of their fold changes. Full list of terms is provided in Table S1. D. NPMI-normalised GAM interaction heat maps of region chr17:30-35 Mb, at 40 kb, centred around the Brd4 gene, indicated by the black dashed line. Below the GAM heat maps, insulation score heat maps indicate contact density. Insulation scores were calculated at 10 insulation square sizes, between 120 and 1200 kb, in steps of 120 kb. TAD boundary locations are indicated underneath insulation score heat maps by dark blue bars. E. Bar plots show the total numbers of expressed and not expressed genes inside versus outside of TAD boundaries in ES and XEN cells. Percentages of genes that are expressed are indicated on the bars. F. Comparison of boundary locations between ES and XEN cells show high levels of cell type-specific boundaries. Boundaries we scored specific, when there was no boundary in the other cell type in at least 80 kb up-or downstream, common were called if boundaries overlap or are directly adjacent. Insulation score profiles are centred around common or specific boundaries and show the same distribution around the boundary centre for common boundaries and different profiles for cell type-specific boundaries. G. GO term enrichment analysis of upregulated genes in cell type-specific boundaries shows enrichment of genes with house-keeping functions in ES cell-specific boundaries and extra-embryonic endoderm functions in XEN cell-specific boundaries. Enrichments were calculated using GOELite using standard parameters. A full list of gene ontology enrichments can be found in Table S2.

To identify genes differentially expressed (DE) between ES and XEN cells, we applied DESeq2 (Love et al., 2014) and found 7421 DE genes, of which 4070 and 3351 are upregulated in ES and XEN cells, respectively (parameters p-adj < 0.05, log_2_(fold change) > 1 or < -1; **Figure 3B**). The number of DE genes was substantially higher than formerly reported (Choi et al., 2020) likely due to increased sensitivity and statistical robustness because of our higher number of biological replicates. Upregulated genes in ES cells had top 20 gene ontologies (GO) related to ‘cell cycle DNA replication’, ‘ribosome’ and ‘DNA methylation and demethylation’, containing *Fgfr1, Dnmt3l, Dppa3, Prdm14* and *Apobec1* (**Figure 3C**; **Table S1**). In XEN cells, the top 20 GO terms often contained endoderm specific genes such as *Lama1, Pdgfra, Col4a2, Sparc* and *Fgfr3*. These results show that ES and XEN cells have highly divergent transcriptional programs, even though they correspond to sister lineages that derive from the same ICM progenitor pool of the embryo.

### TAD boundaries specific to ES or XEN cells contain lineage-specific genes

To begin exploring how changes in gene regulation relate to the reorganisation of the 3D genome structure in ES and XEN cells, we mapped the genomic locations of topologically associating domains (TADs) using the insulation square method at 40 kb resolution (400 kb insulation squares; **Table S2**; Beagrie et al., 2021; Crane et al., 2015). We identified 3088 and 2583 TAD boundaries in ES and XEN cells, respectively, in the range previously described in ES cells by GAM (Beagrie et al., 2021; Winick-Ng et al., 2021) and elsewhere using various TAD calling methods in Hi-C data (Forcato et al., 2017). To help understand the extent of insulation between neighbouring TADs, or their compactness, we also calculated maps of insulation scores using squares from 120-1200 kb (**Figure 3D**).

TAD boundaries have been previously found enriched for housekeeping genes in ES cells (Dixon et al., 2012), and neuronal specific genes in brain cells (Winick-Ng et al., 2021). We first noted that most boundaries overlap genes (85% and 87% in ES and XEN cells, respectively). To investigate the association of TAD boundaries with gene activity, we asked how many TAD boundaries contain expressed genes, and found a highly significant enrichment with 36% and 27% in ES and XEN cells, respectively, compared with 24% and 19% elsewhere in the genome (two-sided Fisher’s exact test; **Figure 3E**).

Next, we measured the extent of TAD reorganisation in ES and XEN cells by comparing boundary positions. TAD boundaries were considered cell-type specific when separated by at least two genomic bins (80 kb), and classified as common when boundaries overlapped or were directly adjacent to each other (where lowest insulation coordinates within boundaries could still be separated by 120 kb; **Figure 3F**). Boundaries that were not directly adjacent but separated by less than 80 kb were not considered as common or specific (‘other’). We found that 41% and 30% of TAD boundaries are specific in ES or XEN cells, respectively. Cell-type specific boundaries are on average weaker than common boundaries, but their specificity to a given cell type is confirmed by a clear depletion of the average insulation compared with the other cell type (**Figure 3F**, lower row).

Finally, we asked whether cell-type specific boundaries contained genes differentially expressed in ES and XEN cells. We found that 9% and 7% of genes up-regulated in ES and XEN cells overlap ES- and XEN-specific boundaries, respectively (**Table S3**). GO analyses revealed that the ES-upregulated genes at ES-specific boundaries often have housekeeping functions, such as mitochondrial protein processing or oxidation-reduction process (**Figure 3G**). In contrast, the XEN-upregulated genes at XEN-specific boundaries have roles in extra-embryonic endoderm identity and function, such as *Lama1/2, Col4a1/2*, and *Robo2*, with terms associated with adhesion and extra-cellular matrix structural constituent. Together, our data show that ES and XEN cell expression programs coincide with lineage-specific rewiring of topological domains, with gene activation co-occurring with loss of chromatin contacts that can lead to the formation of TAD boundaries over activated genes. In contrast, gene repression coincides with increased local contacts and positioning within TADs, away from their borders.

### Key lineage genes are found in conserved compartment A domains in both ES and XEN cells

Next, we investigated compartment A/B classification and their differences between ES and XEN cells (**Figure 4A, Table S4**). Compartments A/B were computed as previously described (Beagrie et al., 2021; Lieberman-Aiden et al., 2009; Winick-Ng et al., 2021), and confirmed to contain genes significantly more highly expressed in compartments A than in compartment B in either cell type (**Figure S3A**). We found that between ES and XEN cells, 45% and 21% of the genome maintains its A or B compartment state, respectively (**Figure 4B**). Approximately one third of the genome changes from A-to-B or B-to-A between ES and XEN cells (15.4% and 18.5%, respectively). However, most differentially expressed genes are located in conserved compartment A in ES and XEN cells (71%; 5250 out of 7421; **Figure S3B**), including major lineage-specific regulators, such as *Sox2, Pou5f1, Nanog, Gata4*/*6, Pdgfra* and *Fgf4* (**Tables S3 and S4**). Genomic windows that acquired compartment A state in ES or XEN cells overlapped with only 5.3% and 7.9% of ES-or XEN-upregulated genes, respectively (**Figure S3B**). Exceptions of lineage-specific genes that underwent compartment B-to-A transitions between ES and XEN cells, include *Lama1* and *Sox17*, two extra-embryonic endoderm genes upregulated in XEN cells by 259 and 3920-fold, respectively. *Lama1* expression increased from 2-4 transcripts per million (TPM) in ES cell replicates to very high expression values of 1287-1520 TPM in XEN cell replicates, which also coincided with the formation of a XEN-specific TAD boundary (**Figures 3G**). Together, these results show that most differentially expressed genes, including many lineage-specific genes, maintain their association with compartment A in both cell types, with notable exceptions including *Lama1*, whose strong upregulation in XEN cells was accompanied by the acquisition of compartment A state.

**Figure 4.**
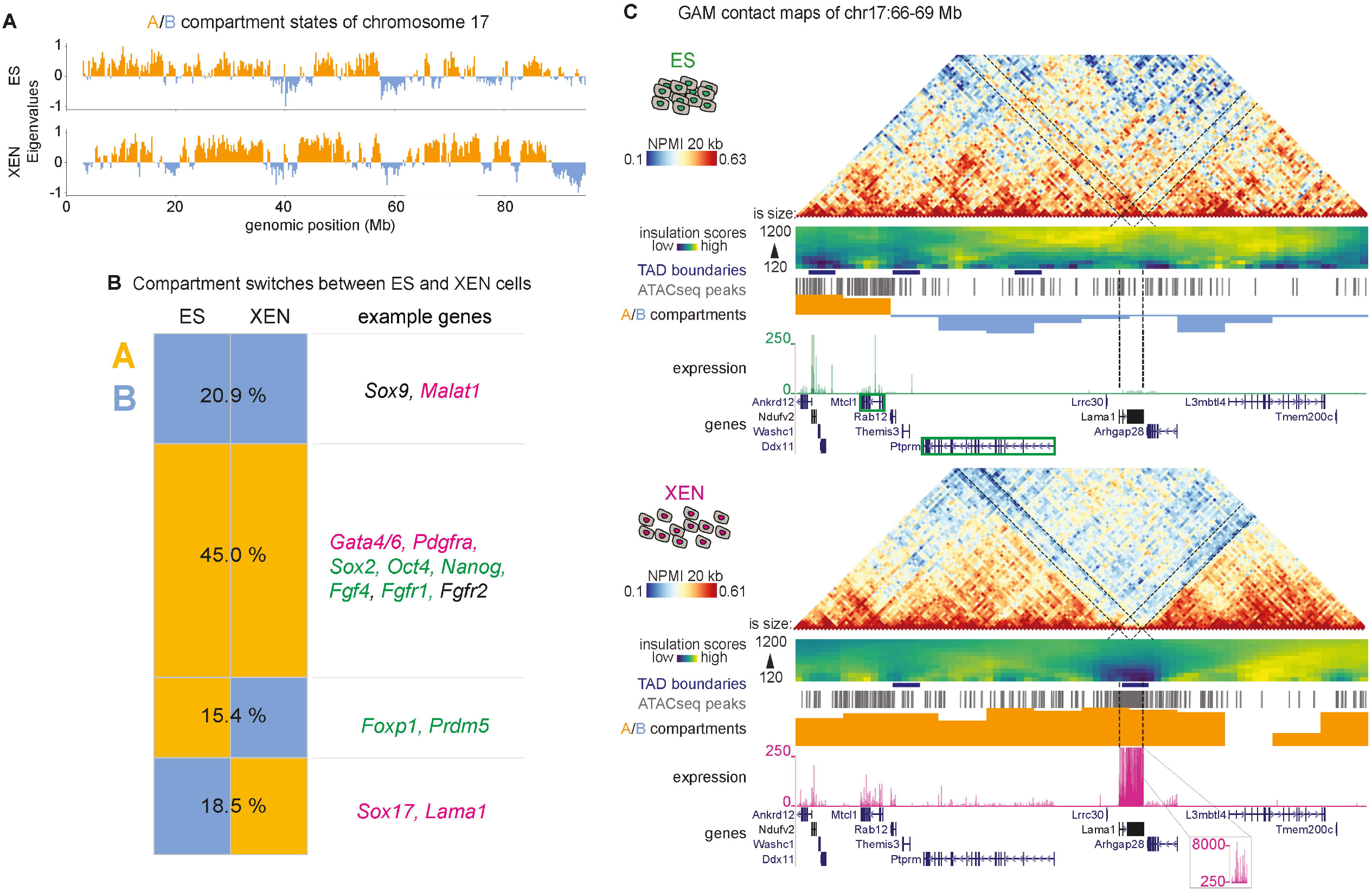
Conserved A compartments contain the majority of lineage driving genes. A. Eigenvalues of the first principal component of principal component analysis of GAM data indicate compartment A (positive sign) and B (negative sign) states. Eigenvalues are normalised between 1 and -1 and shown for chromosome 17. B. Compartment states were compared genome wide between ES and XEN cells in 250 kb windows. Example genes in conserved or switching compartment regions are shown for each group. Genes are coloured in green when they are significantly upregulated in ES cells and in magenta when they are significantly upregulated in XEN cells. C. The genome conformation of the *Lama1* locus is shown by NPMI-normalised GAM data, insulation score heat maps and TAD boundary location, location of ATAC-seq peaks, eigenvalues of the first principal component indicating compartment A/B state and RPKM-normalised RNA-seq data. Gene positions are shown underneath the RNA-seq tracks. The strong upregulation of *Lama1* is concurrent with a switch to compartment A, an increase in chromatin accessibility and a strong loss in insulation in XEN cells.

### Upregulation of Lama1 coincides with its extensive decondensation

The *Lama1* gene spans 125 kb and encodes laminin alpha 1, an extra-cellular matrix factor which is a major component of the Reichert’s membrane, which comprises PrE derived parietal endoderm (ParE) cells. Reichert’s membrane stabilises the embryo inside the uterus and is indispensable for embryo survival (Alpy et al., 2005; Paca et al., 2012; Ueda et al., 2020). To explore the changes in 3D contacts at the *Lama1* locus in greater detail, we plotted GAM contact matrices in ES and XEN cells, over a 3 Mb region of chromosome 17 centred at *Lama1* and which also contains several other XEN-upregulated genes (**Figure 4C**). Weak expression of *Lama1* in ES cells coincided with extensive contacts with neighbouring genes, which were also weakly expressed or silent. The extensive upregulation of *Lama1* in XEN cells was associated with a major loss of contacts between the whole gene body and all its surrounding regions, and the formation of a TAD boundary that covers the whole *Lama1* gene. This loss of contacts is reminiscent of the melting of chromatin contacts which we recently identified with GAM at highly expressed, long (>300 kb) genes in neurons and oligodendroglia (Winick-Ng et al., 2021), showing here that melting can also occur in actively dividing cells and is not an exclusive property of terminally differentiated neuronal cell types.

### ES-repressed Lama1 establishes strong contacts across tens of megabases

To further explore the differences in 3D conformation of the *Lama1* locus in ES and XEN cells, we subtracted their corresponding GAM matrices after z-score transformation (**Figure S4A**). We extracted the top 5% most differential contacts for each ES and XEN cells as cell-type specific contacts, and a similarly sized set of strong common contacts detected in both cell types (**Figure 5A**; Beagrie et al., 2021). In its repressed state in ES cells, *Lama1* established strong cell-type specific contacts with its local and long-range neighbouring genomic regions across a 30 Mb genomic region (green-marked contacts, **Figure 5A**). A closer look at the chromosome-wide interactions of all genomic bins across the *Lama1* gene and its neighbourhood (500 kb region) revealed that its transcription end site (TES) established the largest number of ES-cell specific contacts, compared to the rest of the neighbourhood or its transcription start site (TSS; 499 TES and 346 TSS contacts out of 2268 possible intra-chromosomal contacts), in contrast with few XEN-specific contacts (**Figure 5B**).

**Figure 5.**
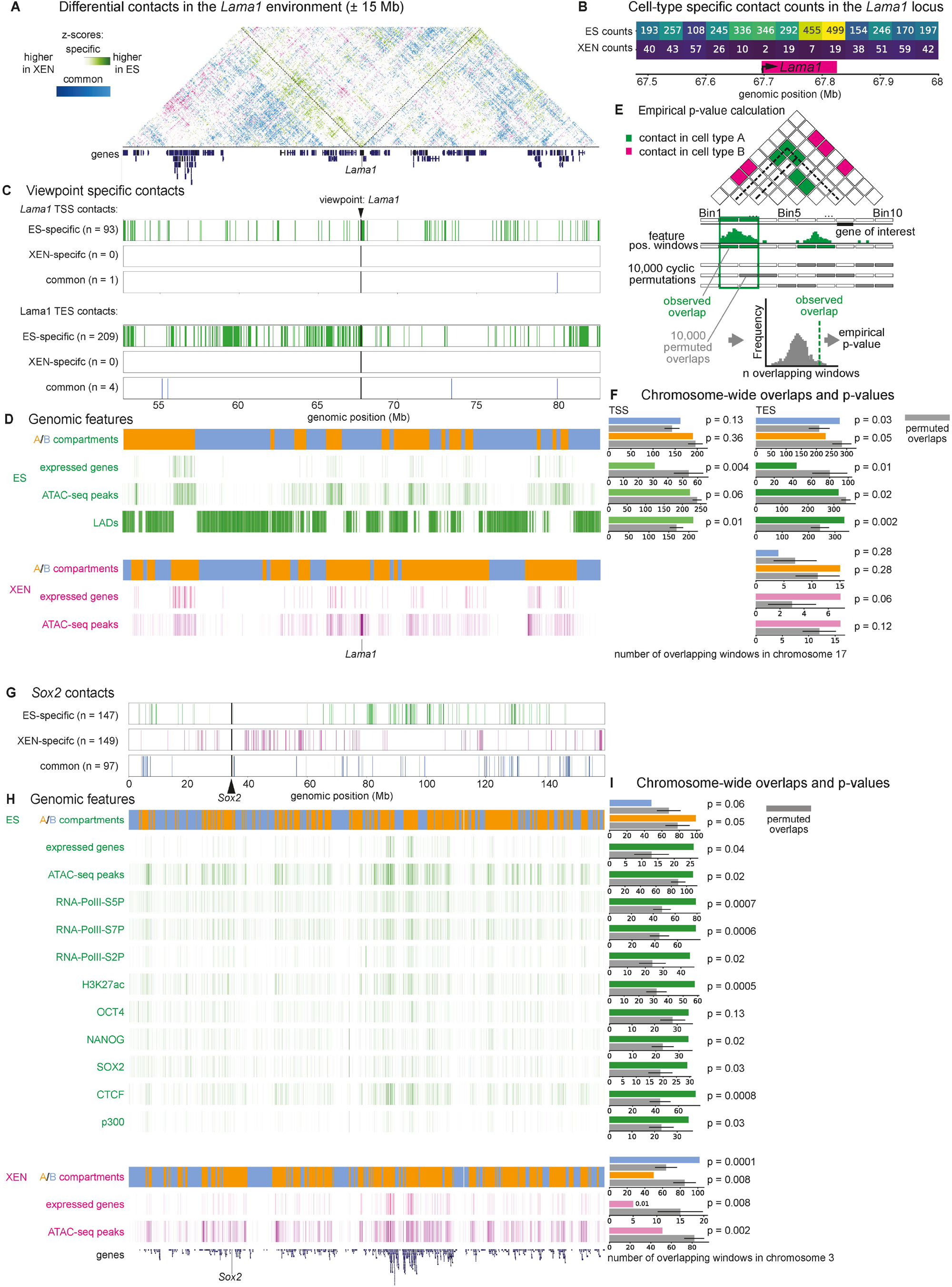
Differential contacts of the *Lama1* and *Sox2* loci show cell-type specificity of genome structures. A. Heat map of a 30 Mb region centred around the *Lama1* gene showing cell-type specific contacts (z-score normalised NPMI GAM data). Magenta shows ES-specific contacts, green XEN-specific contacts, and blue strong common contacts. *Lama1* position is indicated by a dashed line in the heat map. Gene tracks are shown underneath. B. Amounts of ES- and XEN-specific in a 500 kb region centred around the *Lama1* gene shows strong enrichment of ES-specific contacts in ES cells and depletion of contacts in XEN cells. C. Cell-type specific contacts with the *Lama1* TSS and TES in the same genomic region as the heat map in (A). Each bar represents a contact of that region with the *Lama1* viewpoint. D. Genomic features aligned with contact plots in (A) and (C). Shown are positions of A/B compartments, expressed genes, ATAC-seq peaks and LADs for ES cells and positions of A/B compartments, expressed genes and ATAC-seq peaks for XEN cells. E. Schematic explaining the calculation of empirical p-values for enrichments and depletions of genomic features in specifically contacting windows. F. Overlaps of features and empirical p-values for enrichments and depletions of overlaps in cell-type specific *Lama1* contacts. Bar plots refer to overlaps of features aligned in (D). G. Cell-type specific contacts with the *Sox2* gene on chromosome 3. Each bar represents a contact of that region with the *Sox2* viewpoint. H. Genomic features aligned with contact plots in (G). Shown are positions of A/B compartments, expressed genes, ATAC-seq peaks, RNAPolII-S5P/S7P/S2P, H3K27ac, OCT4, NANOG, SOX2, CTCF and p300 for ES cells and positions of A/B compartments, expressed genes and ATAC-seq peaks for XEN cells. I. Overlaps of features and empirical p-values for enrichments and depletions of overlaps in cell-type specific *Sox2* contacts. Bar plots refer to overlaps of features aligned in (H).

To explore the strong interactions of *Lama1* more broadly, we represented the ES-or XEN-specific and common contacts established between the TSS or TES of *Lama1* with other genomic regions, in linear tracks across 30 Mb (**Figure 5C**). We found that the *Lama1* TSS, but especially its TES, established many strong intra-chromosomal contacts in ES cells, but showed no detectable XEN-specific contacts and only a few common contacts, across the whole 30 Mb genomic region. These results show that *Lama1* undergoes extensive decondensation when highly transcribed in XEN cells.

### The silent Lama1 locus contacts B compartments and LADs in ES cells

To further investigate the molecular mechanisms underlying the differences in chromatin contacts between ES and XEN cells, we collected bulk ATAC-seq data in three biological replicates from each cell type (**Figure S4B**). Datasets showed specific detection of open chromatin at active genes and intergenic regulatory regions, as seen at *Gata6* and *Sox2*, with the expected enrichment at gene promoters. We further collected publicly available genomic classifications in ES cells, for lamina-associating domains (LADs; Peric-Hupkes et al., 2010), chromatin occupancy of RNAPII (Brookes et al., 2012; Ferrai et al., 2017), OCT4, NANOG and SOX2 (Marson et al., 2008), as well as H3K27ac and CTCF (Shen et al., 2012).

To shed light on the possible drivers of the strong contacts established by *Lama1* in its repressed state, we began by aligning the contacts established by its TSS and TES with the compartment A/B genomic classification of their interacting windows, the presence of expressed genes or accessible chromatin, and in the case of ES cells, with published classification of LADs (Peric-Hupkes et al., 2010; **Figure 5D**). Visual inspection suggested that *Lama1*, present within a large compartment B in ES cells, but not directly overlapping a LAD, established contacts with other distant compartment B and LAD regions which also contained fewer expressed genes and fewer open chromatin regions. In XEN cells, the transition of *Lama1* to compartment A and its loss of intra-chromosomal contacts coincided with a strong local enrichment of accessible chromatin.

To quantify the significance of the association between the strong *Lama1* contacts with other genomic regions and their linear genomic features, we computed empirical p-values for the chromosome-wide overlaps using a permutation test (10,000 cyclic permutations; **Figure 5E**). In ES cells, the genomic regions that strongly interact with the *Lama1* TES are enriched for B compartments (p-value 0.03), whereas both TSS or TES are significantly depleted for active gene promoters (p-values 0.004 and 0.01, respectively) and enriched for LADs (p-values 0.01 and 0.002; **Figure 5F**). In XEN cells, the *Lama1* TSS has too few intra-chromosomal contacts to calculate enrichments (n=2) and the contacts established by its TES did not show significant enrichments. Other XEN-specific genes also presented specific chromosome-wide contacts. For example, *Gata6* showed significant preference for interactions with other LADs in its repressed state in ES cells (0 TPM), and increased decondensation in XEN cells when upregulated (112 TPM; **Figure S4C**,**D**).

Taken together, these results reveal that *Lama1* and other lineage-specific loci have highly specific 3D conformations in ES and XEN cells, which strongly correlate with their transcriptional state. Our results suggest that the strong upregulation of *Lama1* (from rank 5955 to 16 of the most expressed genes between ES and XEN cells) promotes the extensive topological rearrangements of chromosome 17.

### The active Sox2 locus contacts other active regions enriched for binding of pluripotency TFs in ES cells

To study changes in the organisation of a gene active in ES cells and silent in XEN cells, we chose the *Sox2* locus on chromosome 3, which displays both ES-specific and XEN-specific contacts across its whole chromosome (**Figure 5G**). In ES cells, *Sox2* belongs to a compartment A and contacts other genomic regions containing expressed genes (p-value 0.04), accessible chromatin (p-value 0.02), and RNA polymerase II (RNAPolII) occupancy (p-values ranging between 0.0006 -0.02, for its different phosphorylated forms). The ES-specific contacts were especially highly significantly associated with presence of H3K27ac (p-value 0.0005), and with binding of SOX2 itself (p-value 0.03) and/or NANOG (p-value 0.02; **Figure 5H,I**). In its inactive state in XEN cells, the *Sox2* locus has preferred contacts with compartment B regions and becomes significantly depleted for interactions with active genes or accessible chromatin regions (p-values 0.0001, 0.008 and 0.002, respectively). Together, these findings support the formation of active hubs in the genome of ES cells (Quinodoz et al., 2018), and align with previous reports of long-range contacts established by the *Pou5f1* (Li et al., 2020), or *Nanog* genes with regions bound by pluripotency TFs, Mediator or cohesin (Apostolou et al., 2013; de Wit et al., 2013; Li et al., 2020). Our results show that hubs of active chromatin are cell-type specialised and restructured during the first cell lineage commitment events, early in embryonic development, concurrent with changes in gene activity.

### Accessible Gata4/6 binding sites discriminate ES- and XEN-specific contacts

To further dissect the molecular mediators of cell-type specific chromatin conformations, we searched for TF binding motifs, as a proxy for putative binding events, present at accessible chromatin regions within cell-type specific contacts (**Figure 6A**), as done previously to distinguish neuronal cell types (Winick-Ng et al., 2021).

**Figure 6.**
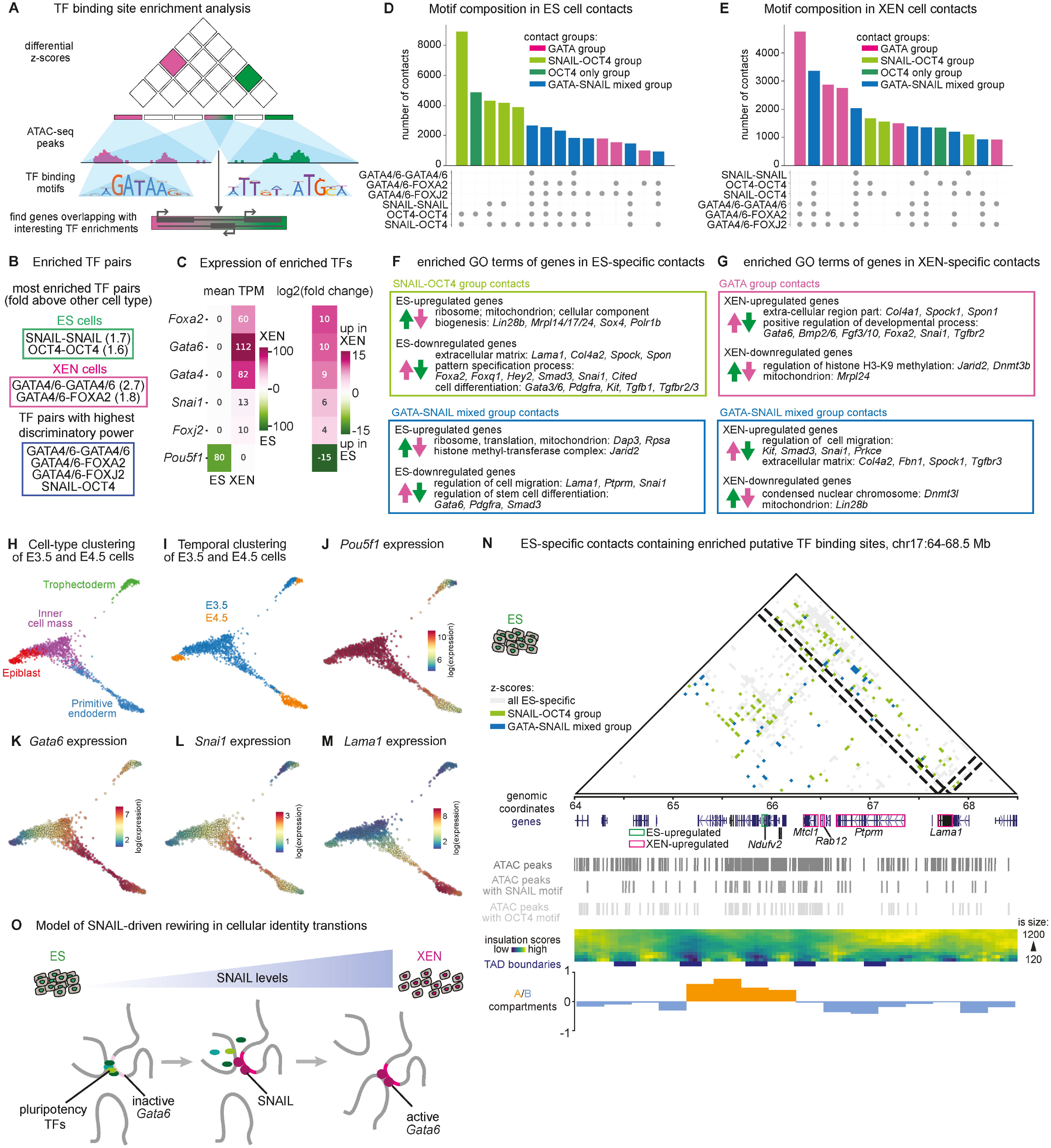
Enrichment of putative binding sites for SNAIL, OCT4 and GATA4/6 characterise differential interactions in ES and XEN cells. A. Schematic describing the workflow of calculating putative TF binding enrichments in cell type-specifically interacting window pairs. B. Enriched TF binding motifs pairs in accessible regions of cell-type specifically contacting window pairs. Enrichments are indicated in fold above other cell type. Discriminatory power is determined by info gain scores. C. Expression (TPM) and log_2_(fold change) of TFs found most enriched of with highest discriminatory power, as reported in (B). A positive log_2_(fold change) indicates upregulation in XEN cells, negative values indicate upregulation in ES cells. D. Most common combinations of TF pairs reported in (B) in ES cell-specific contacts. E. Most common combinations of TF pairs reported in (B) in XEN cell-specific contacts. F. GO terms enriched for up- and downregulated genes in specific groups of contacts in ES cells, merged from analysis in (D), as indicated in the figure legend. Go enrichments were calculated using GOElite with standard parameters, using all expressed genes as background sets. G. GO terms enriched for up- and downregulated genes in specific groups of contacts in XEN cells, merged from analysis in (E), as indicated in the figure legend. Go enrichments were calculated using GOElite with standard parameters, using all expressed genes as background sets. H. Cell-type classifications in scRNA-seq data from E3.5 and E4.5 embryos from Nowotschin et al. (2019). Plots are plotted using https://endoderm-explorer.com/. I. Temporal clustering of cells from E3.5 and E4.5 embryos in scRNA-seq data from Nowotschin et al. (2019). Plots are plotted using https://endoderm-explorer.com/. J. Single-cell expression trajectories of the *Pou5f1* (Oct4) gene in E3.5 and E4.5 embryos, shown in force-directed layouts. Plots are plotted using https://endoderm-explorer.com/. K. Same as in (J) for *Gata6*. L. Same as in (J) for *Snai1*. M. Same as in (J) for *Lama1*. N. ES-specific contacts around the *Lama1* gene (chr17:64.5-68.5 Mb), coloured by contact groups as reported in (D), containing putative SNAIL and OCT4 binding sites (green), or putative GATA4/6 and SNAIL binding sites (blue). Gene tracks under the interaction heat map indicate whether genes are upregulated in ES cells (green) or in XEN cells (magenta). Locations of ATAC-seq peaks, insulation score heat maps, TAD boundaries and compartments states (eigenvalues of the first principal component) are plotted underneath. O. Working model for the function of SNAIL in opening up repressive chromatin networks.

First, we selected TFs that are most likely to contribute to the formation of cell-type specific contacts. We identified the TFs that were significantly differentially expressed between ES and XEN cells (p-adj < 10^−10^; expression > 2 TPM in at least one cell type), and whose putative binding sites were most frequently detected within ATAC-seq peaks in ES-or XEN-specific contacts (at least 20% of genomic windows; **Figure S5A**,**B**). A set of 53 TFs, including SOX2, OCT4, NANOG, GATA4/6 and SOX17, was selected for subsequent analysis, in which GATA4 and GATA6 were combined, due to high degree of motif similarity. Each 40 kb window contained a median of only 2.6 or 2.3 open chromatin peaks, that cover a total median of 520 or 511 bp per window, in ES and XEN cells, respectively, ensuring selectivity in the search process of TF binding sites within cell-type specific contacts (**Figure S5C**). The search specificity was further increased by restricting the analyses to contacts of maximum 5 Mb genomic distance.

Second, we assessed the presence of pairs of TF binding motifs within the accessible regions found in contacting window pairs (**Figure S5D**). TF pairs found in homo- and heterotypic combinations were prioritised for further analyses if they were most enriched in ES-or in XEN-specific contacts, or if they showed highest discriminatory power of cell-type specific contacts measured by information gain score. We found that the most enriched TF pairs in ES-specific contacts were homotypic pairs of putative SNAIL and OCT4 binding sites, whereas XEN-specific contacts were enriched for homotypic pairs of combined GATA4/6, followed by the heterotypic GATA4/6 -FOXA2 pair (**Figure 6B, Table S5**). The pairs of TFs with highest discriminatory power between contacts of the two cell types are the homotypic GATA4/6 pairs, followed by the heterotypic pair of GATA4/6 with either FOXA2, or FOXJ2 and the heterotypic OCT4-SNAIL pair. Out of the six TFs found enriched and/or discriminatory of ES- and XEN-specific contacts, all but OCT4 were upregulated in XEN cells, even if they were found enriched in ES-specific contacts (**Figure 6C**), pointing to a function in resolving contacts rather than mediating them.

Lastly, as many enriched TF pairs could be found in the same set of contacts, we searched for redundancy of contacts and sets of TFs that coexisted therein. We found that the largest subgroup of ES-specific contacts was characterised by both OCT4-OCT4 and OCT4-SNAIL binding sites (8885 contacts), followed by contacts containing only homotypic OCT4 binding sites (4881 contacts), or in addition homotypic SNAIL (4318 contacts; **Figure 6D**). The finding of enriched binding motifs for SNAIL, a repressive TF exclusively expressed in XEN cells, in accessible regions of ES-specific contacts may suggest a role for SNAIL in disassembling or preventing ES-specific contacts in XEN cells. In XEN-specific contacts, the most abundant combination of enriched TF pairs is homotypic GATA4/6 in combination with pairing of GATA4/6 with FOXA2 or FOXJ2 (4745 contacts; **Figure 6E**). These findings suggest a role for the GATA-family of TFs in the organisation of the genome in the extra-embryonic endoderm lineage, supported by pioneering TFs of the Forkhead box (FOX) family, in addition to SNAIL-dependent repression of ES-specific contacts.

### ES-specific hubs of extra-embryonic endoderm genes and regulatory regions contain SNAIL binding sites

To further investigate whether the presence of TF binding sites for SNAIL, OCT4 and GATA4/6 in ES-or XEN-specific contacts related to lineage specific patterns of gene expression, we searched whether they contained lineage-specific genes. To simplify the analyses, we started by combining the most abundant contacts associated with similar sets of TFs. For ES cells, we grouped all contacts that contained the SNAIL-OCT4 pair (SNAIL-OCT4 group, light green), but not the OCT4-OCT4 pair alone (OCT4-only group, dark green), or contacts that contained both GATA4/6 and SNAIL (GATA-SNAIL mixed group of contacts, blue; **Figure 6D**). For XEN cells, we combined contacts containing GATA4/6, FOXA2 and/or FOXJ2 (GATA group, pink), and separately GATA4/6 with SNAIL or OCT4 in various combinations (GATA-SNAIL group, blue; **Figure 6E**). Next, we asked whether these groups of ES- and XEN-specific contacts contained genes that were upregulated in ES or XEN cells. We found that 61% and 54% of ES-upregulated genes were found in the ES-specific contacts containing GATA-OCT4 and GATA-SNAIL (2476 and 2199 genes out of 4070, respectively). Conversely, 51% and 52% of XEN-upregulated genes were found in the XEN-specific contacts containing GATA4/6 and GATA-SNAIL (1696 and 1757 out of 3351 genes, respectively). These observations suggest that the early lineage commitment represented by ES and XEN cells is accompanied by the formation of highly specific hubs of upregulated genes mediated by the activity of transcription factors.

More detailed investigation of ES-specific contacts containing the transcription repressor SNAIL showed the formation of hubs of genes that were significantly less expressed in ES than XEN cells, with roles in the extra-embryonic endoderm, such as *Lama1, Gata3*/*6, Ptprm, Foxa2, Foxq1, Pdgfra* and *Snai1* itself, associated with GO terms ‘extra-cellular matrix’, ‘cell differentiation’ and ‘pattern specification process’ (**Figure 6F**). Further, a large number of SNAIL-OCT4 ES-specific contacts contained XEN-upregulated genes (535 contacts; separated on average 2.7 Mb), which for example connected *Lama1* with *Ptprm* or *Arhgap28*, and *Gata3* with *Proser2* and other repressed genes. The observation that repressed genes, pivotal for ensuring the extra-embryonic endoderm fate, were involved in strong ES-specific chromatin contacts containing putative OCT4 binding sites suggests that OCT4 contributes to ES-specific chromatin topologies associated with gene repression. The presence of SNAIL binding sites at many of the same contacts, implicates SNAIL is interfering or preventing these ES-specific contacts in XEN cells, when *Snai1* is expressed. In contrast, the ES-specific contacts of pluripotency genes, such as *Sox2* or *Pou5f1*, do not contain OCT4-SNAIL or OCT4-OCT4 motif pairs; in general, the GO terms of upregulated genes at these contacts were associated with cellular components, such as mitochondria and ribosomes (full lists in **Table S6**).

Equivalent hubs of active or repressed genes were also found in XEN cells. XEN-specific contacts in the Gata group contained genes up-regulated in XEN cells that have functions related to extra-embryonic endoderm identity and specification, such as *Gata6, Col4a1, Foxa2* and *Snai1*, and are associated with enriched GO terms such as ‘regulation of epithelial to mesenchymal transition’, ‘anatomical structure development’ and ‘mesenchymal cell differentiation’ (**Figure 6G**). Downregulated genes in the GATA group of XEN-contacts included some of the ES-upregulated genes found in ES-specific contacts, such as *Jarid2* and *Mrpl24*, suggesting that these genes transition from strong contacts with specific genomic regions in ES cells to strong contacts with a different set of regions in XEN cells, as their expression status changes from active to repressed. The same phenomenon is seen for ES-downregulated genes, which transition between SNAIL-OCT4 contacts in ES cells to GATA-SNAIL contacts in XEN cells. These results reveal that the two closely related embryonic lineages, ES and XEN cells, undergo strong transitions in chromatin organisation of specifically activated or repressed genes. The switch from one conformation to another, in which a gene locus is restructured, guided by differential binding of TFs, may in turn influence the expression state and potential of the restructured locus and thereby help lock in the two cellular fates.

### Snai1 is upregulated in ICM cells transitioning to PrE

SNAIL is known as a transcriptional repressor important for the down-regulation of E-cadherin through recruitment of histone deacetylases (Peinado et al., 2004) and the initiation of a program of epithelial-to-mesenchymal transitions (EMT) (Yang et al., 2020). To better understand the expression patterns of *Snai1* during early development, we mined published single-cell RNA-seq (scRNA-seq) data from mouse blastocyst stage (E3.5-E4.5) embryos (Nowotschin et al., 2019). Force-directed layout projections of these embryo scRNA-seq datasets show the developmental trajectory between cells of the ICM, to the Epi, the PrE or the trophectoderm (**Figure 6H**), and their maturation timeline between E3.5 and E4.5 (**Figure 6I**). Overlapping *Pou5f1* and *Gata6* expression shows the expression of *Pou5f1* in the ICM, Epi and, at low levels, in the PrE (**Figure 6J**), whereas *Gata6* is expressed in the ICM, and in early and committed PrE cells (**Figure 6K**). In contrast, *Snai1* is expressed weakly in the ICM and upregulated as cells exit the ICM towards the primitive endoderm lineage (**Figure 6L**). While *Snai1* is strongly expressed in E3.5 embryos, its expression becomes reduced in primitive endoderm cells of the late stage (E4.5) blastocyst suggesting that uncommitted ICM cells may undergo a partial and transient EMT-like event in their transition to the extra-embryonic identity. Finally, *Lama1* begins to be expressed in cells that show *Snai1* upregulation and is maximal in the primitive endoderm cells that weakly express *Snai1* (**Figure 6M**).

### ES-specific Lama1 contacts contain putative binding sites for SNAIL and OCT4 and contact other distant repressed genes

To further understand the presence of putative binding sites for SNAIL in ES-specific contacts at repressed extra-embryonic (primitive) endoderm genes in ES cells, we plotted a higher-resolution 4 Mb contact matrix which includes the *Lama1* locus (**Figure 6N**). We found that *Lama1*, especially its TES, was involved in many ES-specific contacts, often connecting *Lama1* with other repressed neighbouring genes and genes more than 1 Mb apart, such as *Ptprm* and *Mtcl1* and that many of ES-specific contacts established by the *Lama1* gene body contain motifs for OCT4 and SNAIL. These observations suggest that SNAIL expression during the transition of ICM cells to the PrE lineage may evoke active dissociation of ES-specific repressive chromatin contacts (**Figure 6O**).

### *Lama1* is less expressed and less decondensed in E4.5 PrE cells than in XEN cells

Finally, we were interested in exploring reports that XEN cells share features of parietal endoderm (ParE) cells which are a derivative of the PrE in the early post-implantation embryo (**Figure S6A**; Kunath et al., 2005). First, we plotted the profiles of gene expression of lineage markers and of *Lama1* and its neighbouring genes, in ES, XEN, Epi, PrE and ParE cells (Nowotschin et al., 2019; **Figure 7A,B**). Comparisons of gene expression profiles in XEN and PrE showed differences for example in *Pou5f1*, which is expressed at low levels in PrE, but not expressed in the XEN cell line used here or in the ParE (**Figures 3A** and **7A**). *Lama1* and its downstream adjacent gene *Arhgap28*, which encodes the Rho GTPase activating protein 28, are more highly expressed in ParE than PrE cells, whereas upstream genes like *Mtcl1* are equally expressed in PrE and ParE cells (**Figure 7B**). The observation that *Lama1* and its neighbouring genes have expression patterns more similar between XEN and ParE, than with PrE, where they are less robustly expressed, led us to hypothesise that *Lama1* might be less decondensed in PrE than XEN cells.

**Figure 7.**
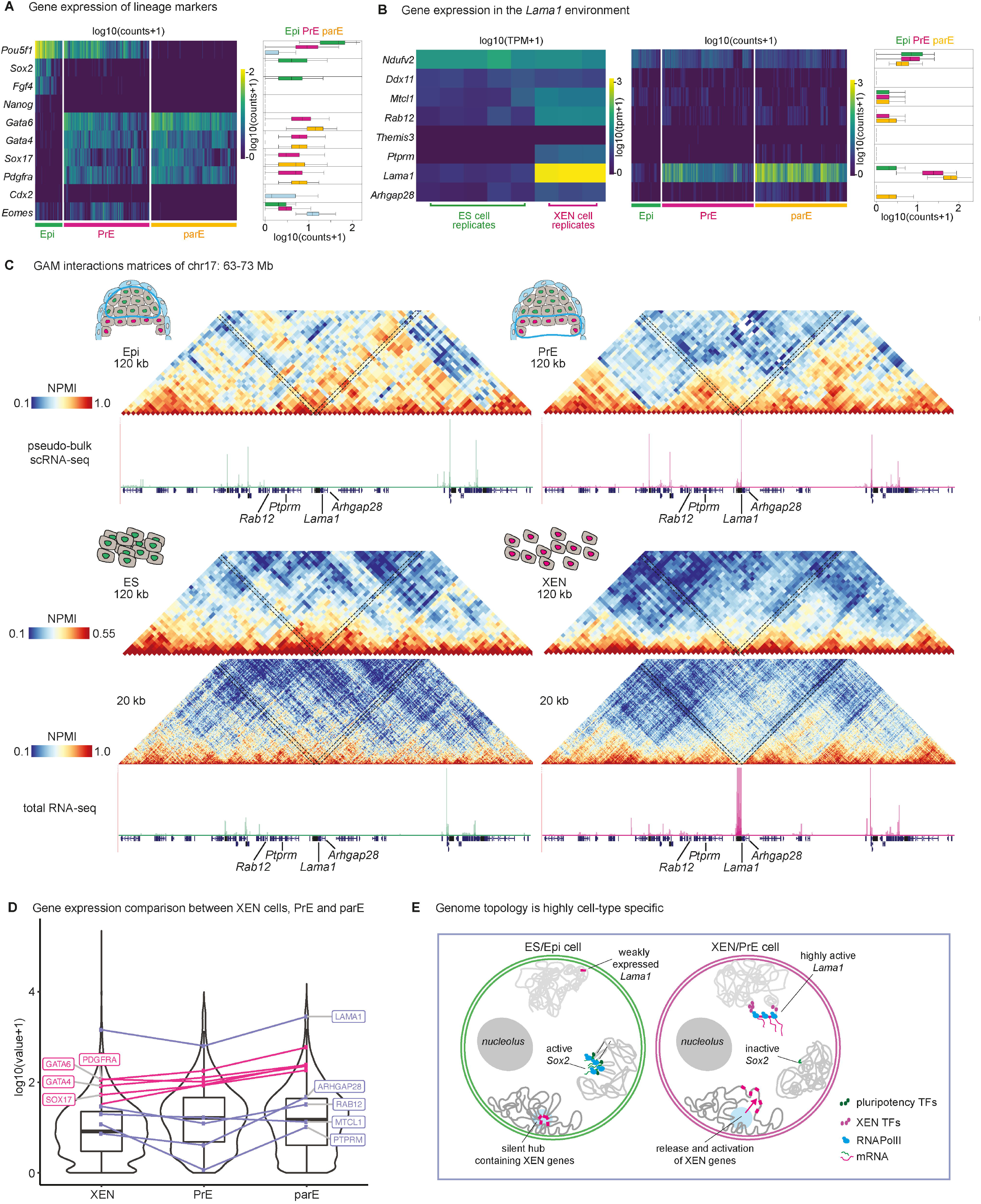
Subtle shifts in cellular identity are reflected in genome topology. A. Heat maps show expression of lineage marker genes in scRNA-seq data from Nowotschin et al. (2019) of cells classified as Epi, PrE and ParE. Count values are log10-scaled with a pseudo-count of 1 added to avoid negative values. Boxplots show distribution of values in single cells of the embryo. Boxplots show distribution of counts in scRNA-seq data across all cells of Epi, PrE and ParE. ParE cells are randomly subsampled from original data, to match number of cells classified as PrE. B. Heat maps show expression of genes in the *Lama1* environment in total RNA-seq of ES and XEN cells and in scRNA-seq of cells classified as Epi, PrE and ParE in Nowotschin et al. (2019). TPM and count values are log10-scaled with a pseudo-count of 1 added to avoid negative values. Boxplots show distribution of values in single cells of the embryo. Boxplots show distribution of counts in scRNA-seq data across all cells of Epi, PrE and TE. C. Interaction heat maps show NPMI-normalised GAM data for the region chr17:63-73 Mb, centred around the *Lama1* gene, at 120 kb resolution in Epi and PrE cells and at 20 kb and 120 kb in ES and XEN cells. Underneath the GAM matrices, pseudo-bulk tracks of scRNA-seq or RPKM-normalised total RNA-seq tracks show gene expression. D. Violin plots show the distribution of TPMs or pseudo-bulk expression values in XEN, PrE and ParE cells. Genes of interest are marked. Boxplots show distribution of expression values, whiskers indicate 1.5IQR. E. Summary of changes in genome topology between embryonic and extra-embryonic cell types, for *Sox2* and *Lama1* loci.

To compare the topology of the *Lama1* locus in ES, XEN, Epi and PrE cells, we visualised GAM contact matrices from ES and XEN cells at resolutions of 20 and 120 kb, the latter to aid comparison with the 120 kb matrices from Epi and PrE cells. Strong contacts between repressed *Lama1* and repressed surrounding genes are seen in both ES and Epi cells, at both 20 and 120 kb resolutions. PrE matrices at 120 kb resolution show loss of chromatin contacts throughout the whole locus, and especially at the *Lama1* gene, although to larger extent in XEN than PrE cells, which included retention of contacts with upstream genes *Ptprm* and *Rab12* (**Figure 7C**), coinciding with their proportionally lower state of activation.

Finally, comparisons of the expression of *Lama1* and its neighbouring genes between XEN, PrE and ParE cells, show the upregulation of *Lama1* and its neighbouring genes in XEN cells, is more similar among XEN and ParE than PrE cells (**Figure 7D**). For example, like XEN cells, ParE cells have higher levels of *Lama1* and *Arhgap28*. While *Lama1* is among the top 1% of genes ranked by expression in XEN and ParE (rank 16 and 62, respectively), it is less highly transcribed in PrE (top 3% or rank 251). Overall, our results show that *Lama1* is highly upregulated in XEN, PrE and ParE cells, where its genomic region is extensively decondensed compared to ES and Epi cells, and that its decondensation is highest in XEN cells where *Lama1* and several neighbouring genes are most upregulated. This suggests that intermediate levels of transcription and decondensation of *Lama1* in PrE cells, may further develop into full decondensation with almost complete loss of contacts chromosome-wide, when *Lama1* becomes more highly expressed in PrE derivatives.

## Discussion

Specification of the three cardinal lineages, two extra-embryonic and one pluripotent embryonic lineage, occurs at the pre-implantation stages and is indispensable for successful development. Despite major technological developments in the field of genome architecture, the changes in chromatin folding that occur during lineage specification in pre-implantation mammalian embryos had remained experimentally inaccessible to genome-wide techniques due to their small size and low cell numbers. Former genomics-based studies showed the establishment of TADs in merged ICMs of blastocysts (Du et al., 2017; Ke et al., 2017), while high resolution imaging revealed differences in chromatin compaction between cells of the Epi and PrE lineages (Ahmed et al., 2010). With the aim of connecting 3D genome structure with emergent cellular identity during early embryonic development, we adapted and applied the immunoGAM method (Winick-Ng et al., 2021) to E4.5 mouse embryos, to precisely distinguish and assign the Epi and PrE cell lineages within the ICM. To our knowledge, no unbiased genome-wide method has so far mapped genome topology in the distinct cells of the ICM of a mammalian blastocyst stage embryo.

Our approach revealed extensive differences in chromatin folding patterns between Epi and PrE cells, showing the importance of distinguishing between the lineages arising within the ICM. Key lineage drivers *Gata6, Nanog* and *Sox2* undergo large genome architecture rearrangements, which resemble those found in *in vitro* datasets of ES and XEN cells (**Figure 7E**). For detailed analyses of genome folding in embryonic (Epi) and extra-embryonic (PrE) lineages, we focused on the high-quality GAM datasets collected from ES and XEN cells, the Epi and PrE *in vitro* counterparts. ES and XEN cells exhibited extensive differences in their gene expression programs, which were often reflected in changes in genome topology. We found extensive rewiring of topological domains (41% and 30% of all TAD borders detected are ES and XEN cell specific, respectively). Importantly, many ES-or XEN-specific TAD boundaries were found to coincide with lineage-specific genes, pointing toward local decompaction of upregulated loci and gene activation creating insulation between the two neighbouring regions of the active gene.

The collected GAM data from XEN and PrE cells shows that the transcriptionally inactive *Sox2* locus establishes extensive intra-chromosomal contacts with other repressed regions. In contrast, the active *Sox2* locus loses interactions with all its adjacent genomic regions in both Epi and ES cells. This result agrees with live imaging studies that showed that *Sox2* transcription does not depend on its local proximity to its downstream super-enhancer, the Sox2-control-region (SCR) (Alexander et al., 2019), even though the SCR is responsible for approximately 90% of *Sox2* expression (Li et al., 2014). Albeit depleted of local contacts, the active *Sox2* locus forms long-range interactions with other active genes and regions marked by H3K27ac, and bound by pluripotency associated TFs, across its whole chromosome, in ES cells. Similar long-range interactions have been reported before for the *Nanog* and *Oct4* loci (Apostolou et al., 2013; de Wit et al., 2013). Together, these findings support the notion that *Sox2* may be regulated through the creation of a specific micro-environment, rather than solely due to enhancer-promoter proximity, potentially involving liquid-liquid phase separation (Boija et al., 2018; Sabari et al., 2018).

*Lama1*, a major component of the Reichert’s membrane, is indispensable for embryonic development, as *Lama1* knock-out embryos die at early post-implantation (Ueda et al., 2020). In its active state in XEN cells, the *Lama1* locus is extensively depleted of contacts both locally and throughout its whole chromosome, concurrent with the establishment of a XEN-specific TAD boundary that coincides with the whole gene. In ES cells, where *Lama1* is marginally expressed, it engages in a dense contact network, characterised by the presence of accessible binding sites of the transcription repressor SNAIL. Whether the extensive *Lama1* decondensation and boundary formation occur first to trigger gene expression, or vice versa, remains to be clarified. While similar topological differences are found *in vivo, Lama1* is less decondensed in PrE cells than in XEN cells, which coincides with lower relative expression of *Lama1* and its neighbouring genes in PrE than XEN cells. These relative differences in expression between *Lama1* and its neighbouring genes suggest that XEN cells have an identity intermediate between PrE and ParE, as previously indicated (Kunath et al., 2005), and that the subtle shift in identity is reflected in differences in genome topology.

SNAIL (*Snai1*), is a known repressor of *Cdh1* (E-Cadherin) and inducer of epithelial-to-mesenchymal transition (EMT) (Cano et al., 2000), which is involved in many important developmental processes, such as gastrulation (Carver Ethan et al., 2001). Over-expression of SNAIL in ES cells induces mesendodermal genes, such as *Foxa2*, and exit from pluripotent, indicating an involvement of SNAIL in early embryonic development (Galvagni et al., 2015). SNAIL binds to promoters of pluripotency-associated genes (Galvagni et al., 2015), pointing to its direct effect as a transcriptional repressor. Conversely, embryoid bodies from *Snai1-*knockout mutant cells showed up regulation of pluripotency genes, and down regulation of endodermal genes such as *Gata4*/6 and *Sox7* (Lin et al., 2014), suggesting multiple cell-type dependent roles. The discovery of SNAIL binding motifs in the ES-specific contacts in our data further suggests that SNAIL may assist in the transition from uncommitted ICM to extra-embryonic (PrE) fate. The co-occurrence of putative SNAIL and OCT4 binding sites points to the interplay of OCT4 and SNAIL in maintaining or dismantling important chromatin interactions. SNAIL binding may help disrupt ES-specific contacts, for example by displacing pluripotency factors, such as OCT4, leading to repression or activation of underlying embryonic (Epi) and extra-embryonic (PrE) genes, respectively. For example, the *Lama1* locus establishes strong ES-specific contacts that contain putative SNAIL and OCT4 binding sites, which are lost in XEN cells, raising SNAIL as a candidate factor involved in the disruption or subsequent prevention of ES-specific contacts. It is worthwhile noting that SNAIL has also been shown to enhance reprogramming efficiency in generating induced pluripotent stem cells, or iPSCs, when adding a SNAIL-NANOG cocktail to the culture, illustrating the complex and likely context- and/or dose-dependent role of SNAIL in differentiation versus pluripotency acquisition (Gingold et al., 2014; Unternaehrer et al., 2014). We suggest that SNAIL may be important for opening up the embryonic genome, and thereby creating accessibility for other TFs to establish new chromatin contacts in transitions to other cell types. Whether SNAIL can act as a pioneering TF, or requires other TFs remains to be investigated.

The emergence of the extra-embryonic (primitive) endoderm states *in vivo* and *in vitro* relies on the TFs GATA4 and GATA6 (Fujikura et al., 2002; Schrode et al., 2014; Shimosato et al., 2007; Wamaitha et al., 2015). Several other GATA-family transcription factors have been shown to help rewire chromatin contacts (Jing et al., 2008), while GATA-family TFs have also been shown to dimerise in homo- and heterotypic pairs, when bound to DNA (Bates et al., 2008). In our motif enrichment analysis, GATA-family TFs show the highest discriminatory power between contacts specific to ES or XEN cells, as well as the highest enrichment in XEN cell-specific contacts. Known characteristics of GATA-family TFs, together with our results, support the notion that GATA4 and GATA6 are involved in structuring the genome to regulate expression of genes important for the PrE state.

Our work demonstrates the power of the immunoGAM technology in assessing 3D genome folding in delicate samples with limited cell numbers, such as early mammalian embryos, while retaining their structural organisation by distinguishing cell types in intact tissues. In this way, GAM provides opportunities for detailed analyses of 3D genome dynamics across state transitions within developing embryos, which will be facilitated by the development of multiome-GAM technologies. Finally, the extensive rewiring of 3D genome folding observed between Epi and PrE lineages is often detected at lineage specification genes, such as *Sox2* and *Gata6*, revealing that the specialisation of genome folding is tightly related with lineage specification.

## STAR Methods

### RESOURCE AVAILABILITY

#### Lead contact

Further information and requests for resources and reagents should be directed to and will be fulfilled by the lead contact, Prof. Ana Pombo.

#### Materials availability

This study did not generate new unique reagents.

#### Data and code availability

Raw fastq sequencing files for all samples from XEN, Epi and PrE GAM datasets, together with non-normalised co-segregation matrices, normalised pair-wise NPMI chromatin contacts maps and raw GAM segregation tables have been submitted to the GEO repository under accession number GSE195485 and also available from the 4DN data portal (https://data.4dnucleome.org/) under the accession numbers 4DNESTG7CNKF, 4DNESF3VXIMV and 4DNESUENC64Y. Raw fastq sequencing files for 3NP mES cell GAM datasets are available from https://data.4dnucleome.org/ under the accession number 4DNESALAVZ67 (Beagrie et al., 2021).

Raw fastq sequencing files for ES and XEN cell ATAC-seq datasets, together with bigwig files for displaying read density tracks, as well as consensus peak files called from 3 biological replicates have been submitted to the GEO repository under the accession number GSE196080.

Raw fastq sequencing files for XEN cell RNA-seq, together with bigwig files for displaying read density tracks have been submitted to the GEO repository under the accession number GSE196389.

All deposited data will be publicly available as of the date of publication. Availability of published data is indicated in Table S8.

This paper does not report original code, the code used in this study was published in Beagrie et al. (2021) and Winick-Ng et al. (2021), and used with minor modifications, as described in the Methods.

### EXPERIMENTAL MODEL AND SUBJECT DETAILS

#### Cell culture

XEN cell clone IM8A1 was a gift by Janet Rossant (Kunath et al., 2005). ES cell clone 46C (Ying et al., 2003), a Sox1–GFP derivative of E14tg2a, was a gift by Domingos Henrique.

#### Animals

All work with live animals was performed in the laboratory of Anna-Katerina Hadjantonakis at Memorial Sloan Kettering Cancer Center (MSKCC) in New York City, in accordance with guidelines from MSKCC Institutional Animal Care and Use Committee (IACUC) under protocol no. 03-12-017 (principal investigator AKH).

This study used wild type 4-to 12-week-old CD1 females and CD1 stud males, strain code 022, by Charles River.

### METHOD DETAILS

#### Cell culture

XEN cells (clone IM8A1) were grown as previously described (Stock et al., 2007). ES cells (clone 46C) were grown as described (Beagrie et al., 2017).

#### Total RNA-seq

For RNA isolation, ES or XEN cells were grown to 70-80%, washed in culture media while adherent to the dish, and then lysed and washed off the dish with TRIzol reagent (Invitrogen, 15596026) at room temperature in three independent biological replicates per cell type. Lysates were snap-frozen in liquid nitrogen, and stored at -80 °C until further processing. Samples were incubated at room temperature for 5 min and homogenised with 200 µl chloroform per 1 ml TRIzol, by shaking for 15 s and incubating 3 min at room temperature. Samples were centrifuged at 12,000 x g for 15 min at 4 °C and the upper aqueous phase was transferred to a new tube and RNA was precipitated by adding 500 µl HPLC-grade isopropanol, incubating 10 min at room temperature and pelleting RNA by centrifugation at 12,000 x g for 10 min at 4 °C. Supernatant was removed and pellet was washed with 75% ethanol, air-dried for 10 min and finally resuspended in RNase-free water and incubated at 55 °C for 10 min. DNA was removed by DNase treatment (Turbo DNase, AM2238, ThermoFisher) according to manufacturer’s protocol. RNA was stored at -80 °C until further processing.

Purified RNA was tested for sufficient quality, ensuring intact, non-degraded RNA, by using the Bioanalyzer with the Agilent RNA 6000 Nano kit (Agilent Technologies, 5067-1511). RNA-seq libraries were generated from 1 µg of clean RNA using the TruSeq Stranded total RNA library preparation kit (Illumina, 20020596) according to the manufacturer’s protocol. Libraries were analysed on the Bioanalyzer using the Agilent High Sensitivity DNA kit, giving out the average fragment size of the libraries. Molarity was estimated according to:

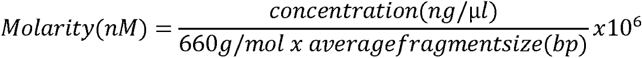

DNA concentrations were measured by Qubit Quant IT kit according to manufacturer’s protocol. ES cell RNA libraries were sequenced using the HiSeq4000 system at 150 bp length, paired-end, according to the manufacturer’s protocol. XEN cell RNA libraries were sequenced on the NextSeq, at 75 bp length, paired end, according to the manufacturer’s protocol. Two additional ES cell RNA-seq replicates were downloaded from the GEO repository under the accession number GSE148052 and differed only in read length, 100 and 75 bp and were sequenced in HiSeq2000, respectively. Sequencing depth ranged from 54-126 million reads in ES and XEN cell replicates 1 to 3 and between 163-187 million reads in ES cells replicates 4 and 5.

Therefore ES cell replicates 4 and 5 were excluded from visual comparisons of genome browser tracks (**Figure S2A**).

#### Assay for Transposase-Accessible Chromatin using sequencing (ATAC-seq)

For ATAC-seq, XEN or ES cells were grown to 60-70% confluency in 10 cm diameter dishes. Cells were placed on ice, gently scraped and resuspended in culture media (serum+LIF for ES cells and serum only for XEN cells). Approximately 50,000-75,000 cells were aliquoted on ice and directly used as input for the ATAC-seq experiment. ATAC-seq was done using the published protocol for Omni-ATAC (Ackermann et al., 2016) (https://www.med.upenn.edu/kaestnerlab/protocols.html) with the minor modification of using a Tn5 enzyme produced by the MDC Protein Purification and Characterisation platform. Briefly, cells were lysed and nuclei were extracted as described in the original protocol. Tn5 mediated transposition was done in intact nuclei. DNA was then purified and used to generate Illumina sequencing libraries using the Nextera XT kit or an in-house protocol. Samples were sequenced at a depth of 50-70 million reads with the NextSeq 500/550 v2.5 kit. ES replicate 3, produced in the exact same manner as one and two, was already published in Winick-Ng et al., 2021 and can be accessed through the GEO repository with the accession number GSE174024 and for the sake of completeness is also included in the GEO repository of this publication under the accession number GSE196080.

#### Tn5 loading for ATAC-seq and sequencing library preparations

Tn5 (Picelli et al., 2014) was loaded with short DNA oligonucleotides as follows. Lyophilised oligo primer stocks were reconstituted to 200 µM with 1x Tris-EDTA (TE) buffer. In two reactions, equal volumes of Tn5MErev and Tn5ME-A, and Tn5MErev and Tn5ME-B (key resources table) were mixed and incubated for 5 min at 95 °C and gradually cooled down to 25 °C, at 0.1 °C/s. Next, adapter reagents were mixed at a 1:1 ratio to generate a 100 µM adapter mix. Tn5 (1.85 µg/µl) was loaded by adding 1 volume of Tn5 to 0.143 volumes of 100 µM adapter mix. The reaction was incubated for 1 hour at 25 °C and stored at -20 °C until further use.

#### Karyotyping of XEN cell clone IM8A1 and ES cell clone 46C

XEN and ES cells were grown, trypsinised and washed as described above. Genomic DNA was extracted by washing the cell pellet in PBS, centrifuging (5 min at 190 x g) removing supernatant and adding 500 μl lysis buffer (0.6 % SDS, 150 mM NaCl, 10 mM EDTA, 10 mM Tris-HCl (pH 8.0)) plus 10 μl Proteinase K (10 mg/ml, Sigma Aldrich #03115836001). The sample was incubated over night at 50 °C and DNA was recovered using phenol-chloroform extraction, by adding 500 μl phenol-chlorophorm-isoamylalcohol (Merck, #KP31757), mixing by inverting, and centrifuging at 21,000 x g for 10 min. The upper aqueous phase was transferred to a new micro-centrifuge tube, 500 μl chloroform were added and mixed by inverting. The sample was centrifuged for 10 min at 21,000 x g and the upper aqueous phase was transferred to a new tube and 1 ml ice-cold Ethanol was added to precipitate DNA. After centrifugation for 5 min at 21,000 x g, the supernatant was discarded and the DNA pellet was washed in 70 % ethanol. Then the supernatant was removed and the pellet was dried at 37 °C until visually dry. DNA was resuspended in 1x TE buffer or water. Array-comparative genome hybridisations (array-CGH) on genomic DNA were carried out by Atlas Biolabs (Berlin, Germany) as a commercially available service, using the Agilent SurePrint G3 Mouse CGH Microarray, 4×180K system (**Figure S6B**). Figure S6B was plotted using the Agilent Genomic Workbench (https://www.agilent.com/en/download-software-agilent-genomic-workbench) and shows XEN replicate 3. Reports of all XEN and ES samples can be found in supplementary data file 1. Inspection of array CGH results showed amplification of chromosomes 1, 6, 11, 12, 14, 15, 16 and 19. Their analyses are possible within XEN contact maps due to the fact that NPMI normalisation accounts for the number with which each genomic window is detected. To avoid possible biases in the differential contact analyses comparing ES and XEN cell matrices (shown in **Figure 6**), the amplified chromosomes were excluded from those analyses.

M-FISH was performed as previously described (Azawi et al., 2020). Briefly, metaphase chromosomes of cytogenetically prepared XEN cells were stained by applying all 21 murine whole chromosome painting probes using a commercially available kit (Applied Spectral Imaging, Edingen-Neckarhausen, Germany; SKY Paint DNA Mouse #FPRPR0030). The probes were hybridised and evaluated according to manufacturer’s instructions and as previously described (Azawi et al., 2020). Evaluation was done in 20 metaphase spreads using a fluorescence microscope (Axioplan 2 mot, Zeiss) equipped with appropriate filter sets to discriminate between all five fluorochromes (SpectrumOrange, SpectrumGreen, TexasRed, SpectrumAqua, Cyanine 5) and the counterstain DAPI (Diaminophenylindol). Image capturing and processing were carried out using an ISIS mFISH imaging system (MetaSystems, Altlussheim, Germany) and showed a translocation between chromosomes 6 and 14 (**Figure S6C**), which were subsequently removed from differential contact analysis. G-banding was performed on three replicates of XEN cells cytogenetically prepared as for M-FISH, and evaluation of 100 metaphase spreads per replicate showed that 7-12 % of karyotypes were tetraploid.

#### Immunofluorescence of whole cells

Cells were grown on ethanol-washed and autoclaved glass coverslips for 1-3 days under regular conditions. For fixation, cells were rinsed in 4% EM-grade Paraformaldehyde (Alfa Aesar #43368) in 250 mM HEPES-NaOH (pH 7.6) and fixed for 10 min at room temperature in fresh 4% EM-grade Paraformaldehyde in 250 mM HEPES-NaOH. Samples were washed in PBS for 3 min (3 × 10 min), incubated in 20 mM glycine in PBS and permeabilised for 10 min in 0.5% Triton X-100 in PBS (w/v). Samples were incubated in blocking solution for 1 h (1% BSA, 0.1% Casein, 0.2% fish skin gelatin, in PBS, pH 7.8). Primary antibodies were diluted in blocking solution and incubated overnight at 4 °C in a humid chamber. Dilutions of primary antibodies were 1:100 for GATA6, SOX17, SOX2 and OCT4; and 1:500 for NANOG. After primary antibody incubation, cells were washed in blocking solution for one hour (3 × 20 min) and incubated with secondary antibody (dilution 1:1000 in blocking solution) for 1 h at room temperature (all antibodies indicated in key resource table). Finally, samples were washed for 1 h in blocking solution (3 × 20 min) at room temperature, before incubation (3 min) with 0.5 µg/ml DAPI in PBS, washed in PBS (3x) and mounted in Vectashield.

Immunostained ES or XEN cells grown on glass cover slips were imaged using the Leica SP8 laser-scanning confocal microscope. Images were acquired using a 63 x oil objective (numerical aperture (NA) 1.4), and a pinhole equivalent to one airy disk. Separate fluorescence channels were always collected subsequently to avoid bleed-through. Raw images were contrast stretched using ImageJ. Images in comparative experiments were always recorded and contrast-stretched using the same parameters.

#### Animal husbandry and mouse embryo collections

All following experiments used wild-type CD1 mice (Charles River), housed in a pathogen-free facility under a 12 h light/dark cycle in the MSKCC mouse facility.

Embryos were obtained from natural matings of 4-to 12-week-old females to stud males, with noon of the day at which a vaginal plug was detected considered E0.5. Embryos were collected at E4.5. For embryo collection, mice were sacrificed by dislocation of the neck, uteri were removed and uterine horns were flushed with M2 media (MTI GlobalStem, GSM-5120). Embryos were washed through three drops of M2 media and briefly kept in M2 at room temperature until fixation.

#### Immunofluorescence of whole embryos

For immunofluorescence of embryos, sequential steps were carried out in a soft-plastic U-bottom 96 well plate, the bottom of all wells were covered with agarose solution (1% Agar, 0.95% NaCl). Embryos were washed in 0.1% Triton X-100 in PBS, for 5 min, followed by permeabilisation for 5 min in 0.5% Triton X-100 in PBS, containing 100 mM glycine in PBS and again washed in 0.1% Triton X-100 in PBS. Then they were incubated in blocking solution (2% horse serum in PBS, Sigma Aldrich, H0146) for one hour at room temperature. The primary antibody was incubated overnight at 4 °C, diluted in blocking solution. During all incubations longer than 10 min, solutions were covered in mineral oil to prevent drying. After primary antibody incubation, embryos were washed 3 × 5 min in 0.1% Triton X-100 in PBS, and again blocked for 1 h at room temperature in blocking solution.

Secondary antibodies were incubated for 1 h, at 4 °C, diluted in blocking solution. Finally, the embryos were washed 2 × 5 min in 0.1% Triton X-100 in PBS and stored in PBS at 4 °C until imaging.

Immunostained whole embryos were imaged using the Zeiss LSM700 confocal microscope on glass-bottom dishes. Images were acquired using a 63x oil objective (numerical aperture (NA) 1.4) or 40 x objective (NA 1.4), and a pinhole equivalent to one airy disk. Separate fluorescence channels were collected subsequently to avoid bleed-through. Raw images were contrast stretched using ImageJ. Images in comparative experiments were collected and contrast-stretched using the same parameters.

#### Processing of embryos for GAM

Embryo samples were chemically fixed directly after collection from the mother animal by washing them in 250 mM HEPES solution (pH 7.6) at room temperature and incubating them (for 2 h, 4 °C) in 4% EM-grade Paraformaldehyde in 250 mM HEPES, followed by transfer and incubation (30 min, 4 °C) in 8% EM-grade Paraformaldehyde in 250 mM HEPES (pH 7.6). Embryos were stored in 1% EM-grade Paraformaldehyde in 250 mM HEPES (pH 7.6) at 4 °C until further processing, typically 7-21 days, while 1% EM-grade Paraformaldehyde in 250 mM HEPES was renewed once per week. For cryoblock preparation, embryos were embedded in 12% gelatin (Merck, 104070) in PBS. After heating up to 42 °C, 15 µl gelatin solution were placed in a 1.5 ml low-binding micro-centrifuge tube and kept warm at 37 °C until further processing. Embryos were washed 3x 5 min in 0.5% BSA, 0.1% glycine in PBS, to quench free aldehydes. Then, batches of 20-30 embryos were manually transferred into the warm gelatin solution, using a long, L-shaped glass pipette, pulled from a Pasteur pipet after melting in an open flame. The embryos were embedded in the gelatin by incubation at 37 °C for 2 h, centrifuged for 2 min at 50 x g and then left to solidify at 4 °C overnight. The tip of the tube was cut off with a razor blade and a drop of 2.1 M sucrose in PBS was added on top of the gelatin block. The solid gelatin block was detached from the walls of the tube with a wooden tooth pick that was autoclaved and flattened with a scalpel. Once detached, the gelatin block was flushed out into a small dish by injecting 2.1 M sucrose in PBS with a pipette behind the gelatin. The sample was then incubated in 2.1 M sucrose in PBS for 24 h, under gentle shaking in a humid chamber. After incubation, the gelatin was cut with a scalpel to remove excessive gelatin. The remainder smaller blocks of gelatin containing embryos were mounted on copper stubs. Gelatin blocks on copper stubs were frozen in liquid nitrogen under constant shaking and stored submersed under liquid nitrogen indefinitely.

Cryosections were cut at -110 °C with an ultracryomicrotome (Leica Biosystems EM UC7) using glass knives. Sections were cut at ~220 nm thickness, assessed by refractive index, and captured in a drop of 2.1 M sucrose in PBS, suspended in a copper wire loop. The drops with the sections were transferred to 4 µm thick polyethylene naphthalate (PEN) membrane metal frame slides (Leica Biosystems, 11600289), for GAM, or on ethanol-washed and autoclaved glass coverslips for confocal microscopy.

#### Immunofluorescence staining of embryo cryosections

Embryo cryosections were stained by immunofluorescence using SOX2, OCT4, GATA6 and SOX17 antibodies. After washing, primary antibodies recognising SOX2 and OCT4 were indirectly immunostained with fluorochromes with similar emission wavelength (to mark Epi cells) but distinct from the fluorochromes used to indirectly immunolabel GATA6 and SOX17 (key resources table). The overall immunofluorescence and confocal imaging protocols for cryosections are similar to what is described above for cryosections from ES and XEN cells, with minor modifications. Cryosections from gelatin-embedded embryo samples were washed (5 min) in PBS (heated to 37 °C in a water bath), followed by washes (3 × 10 min) in 2% gelatin in PBS (heated to 37 °C in a water bath) and (2 × 10 min) in warm PBS. Furthermore, for embryo sections, blocking buffer was changed to PBS+HS (2% horse serum, 0.1% casein, 0.2% fish skin gelatin). Antibodies were diluted in respective blocking solutions and primary antibody concentrations were increased to a 1:50 dilution.

### GAM data production

#### Staining of cryosections and laser microdissection

XEN cell samples were produced for cryosectioning as previously described (Beagrie et al., 2017) and XEN cell cryosections were stained using cresyl-violet as described in Beagrie et al., 2021.

Embryos were cryosectioned and stained as described above, with minor modifications. Compared to staining for confocal imaging which uses primary and secondary antibodies, also tertiary antibodies, recognising the secondary antibodies, were used to amplify fluorescent signal further. This was done similar to the secondary antibody, by washing in PBS+HS (3x 20 min) and incubating in PBS+HS with tertiary antibody (1:1000) for 1h. Further, glass coverslips were exchanged for PEN membrane slides, on which sections were finally washed 3 times in H_2_O and air-dried for 10 min after immunofluorescence staining, before laser micro-dissection.

Samples that were stained for collection of GAM samples were treated exclusively with sterile-filtered solutions (0.22 µm filter).

Nuclear profiles (NPs) of choice were laser micro-dissected from the PEN membrane using the Leica LMD7000 laser micro-dissection microscope. NPs were collected into adhesive PCR caps (AdhesiveStrip 8C opaque; Carl Zeiss Microscopy #415190-9161-000) and presence of NPs in caps was confirmed with a 5x objective using a 420-480 nm emission filter. Controls without nuclear profiles (water controls) were included for each dataset collection. The example image of stained E4.5 embryo section on PEN-membrane shown in **Figure 2F** was collected at 63x magnification with the laser-microdissection microscope (LMD700, Leica).

#### Whole genome amplification, library preparation and next generation sequencing of GAM samples

XEN cell GAM samples were amplified using the Yikon genomics MALBAC Single Cell WGA kit (Yikon Genomics, #EK100101210) or an in-house developed whole-genome amplification (WGA) approach (**Table S7**; Winick-Ng et al., 2021)). DNA from embryo samples was amplified using an adaptation of the in-house protocol, in which lysis was prolonged to 24 h, at 60 °C and the volume of Qiagen Protease solution added was increased to 3 µl. After WGA samples were cleaned up using SPRI beads (1.7x beads to sample volume ration).

Sequencing libraries of XEN and embryo GAM samples were produced using the Illumina Nextera XT library preparation kit (Illumina #FC-131-1096) following the manufacturer’s instructions with either full or an 80% reduced volume of reagents or with an in-house tagmentation-based library preparation protocol (Picelli et al., 2014 with modifications), as indicated in Table S7. Primer removal and average fragment size of libraries was tested using a DNA High Sensitivity on-chip electrophoresis on an Agilent 2100 Bioanalyzer, according to the manufacturer’s protocols. Concentration was measured using the Quant-iT® Pico Green dsDNA assay kit (Invitrogen #P7589). Then, 192 samples were pooled together by taking the same amount of DNA (5-8 ng) from every sample. Pooled libraries were cleaned up two times using 1.7 x SPRI beads. Samples were sequenced in batches of 192 samples per run, using the NextSeq500 system with the NextSeq 500/550 High Output v2 kit (75 cycles) at approximately 2 million reads per sample.

### GAM computational analyses

#### Sequence read alignment and sequencing quality control

Fastq files were demultiplexed, Illumina and WGA adapters were removed and reads were mapped to the mouse reference genome NCBI build 38/mm10 with Bowtie2 (version 2.3.4.3) using default settings. All non-uniquely mapped reads, reads with mapping quality <20 and PCR duplicates were excluded from further analyses as previously described (Beagrie et al., 2017).

#### Calling positive windows, quality controls and contact matrix resolutions

Processing of GAM data was as described in Winick-Ng et al., 2021 (https://github.com/pombo-lab/WinickNg_Kukalev_Harabula_Nature_2021), and applied with minor modifications, for example in cut-offs for exclusion of low-quality samples. To call positive windows, the genome was split into equally sized bins of 40 kb resolution.

Each batch of samples that was processed in the same whole-genome amplification reaction in one common 96-well reaction plate, was subjected to a cross-contamination analysis. To that end, we calculated a Jaccard similarity index for each pair of nuclear profiles, based on their positively called windows. All sample pairs with an index > 0.4 were sorted out and excluded from further processing. All remaining samples were assessed for quality by determining the percentage of orphan windows (positive windows without other directly adjacent positive windows), and number of uniquely mapped reads in each sample. GAM samples from XEN cells with < 20,000 uniquely mapped reads or > 70% orphan windows were excluded from further analyses. Similarly, PrE and Epi samples with < 20,000 uniquely mapped reads or > 60% orphan windows were excluded. The publicly available ES cell dataset was produced with a different WGA protocol, therefore was quality controlled using the same strategy, but required cut-offs of < 250,000 uniquely mapped reads (due to increased WGA noise) and > 70% orphan windows.

#### Visualisation of pairwise chromatin contact matrices

The normalised point wise mutual information (NPMI) measure is used to transform raw GAM co-segregation matrices into pairwise contact matrices while normalising for local differences in detection efficiency (Winick-Ng et al., 2021), https://github.com/pombo-lab/WinickNg_Kukalev_Harabula_Nature_2021). Briefly, point-wise mutual information (PMI) describes the difference between the probability of finding two genomic windows in a common nuclear profile (NP) and their individual distributions across all NPs. It assumes that the probability of finding one window is independent from finding a second one. Normalised PMI (NPMI) values are obtained by adding a correction factor bounding the PMI values to -1 and 1.

For visualisation, NPMI values are plotted as heat maps, where colour scales are adjusted to range between 0 value and the 98th percentile of NPMI values for each matrix.

#### A/B compartment calling

A/B compartments were identified as previously described (Winick-Ng et al., 2021, https://github.com/pombo-lab/WinickNg_Kukalev_Harabula_Nature_2021).

#### TAD boundary calling using the insulation square method

The insulation score method was applied to identify topologically associating domains (TADs) and TAD boundaries genome wide in GAM data (Crane et al., 2015). A slightly modified version was used, adapted to consider negative values in GAM data (Winick-Ng et al., 2021; https://github.com/pombo-lab/WinickNg_Kukalev_Harabula_Nature_2021).

The insulation square method is preferred for TAD calling from GAM data, as it was previously shown to identify TAD boundaries also found in Hi-C data (Beagrie et al., 2021). To call TAD boundaries, data with a 40 kb resolution and an insulation square size (iss) of 10x the resolution (400 kb) was used. Additional parameters used were boundary margin of error (bmoe) = 1, resulting in a boundary size of minimum 120 kb, comparable to Crane et al. (2015), and insulation delta span (ids) = 80 kb.

Boundaries that the insulation square algorithm calls separately but that actually overlap or at least touch, were merged using bedtools (v 2.29.2; Quinlan and Hall, 2010) merge function.

The insulation square pipeline was also used to create contact density heat maps using different square sizes (from 3 x to 30 x the resolution, in steps of 3 x resolution, ie from 120 kb to1200 kb), as previously done in Winick-Ng et al. (2021). Resulting insulation scores are plotted as heatmaps and indicate the contact density of genomic regions.

#### Calling cell-type specific contacts

Cell-type specific contacts were called as previously described (Beagrie et al., 2021). In brief, windows with a detection frequency of less than 4% or more than 32% in XEN cells or less than 6% or more than 40% in ES cells were removed. NPMI contact frequencies at each genomic distance were normalised by z-score transformation, and a differential contact matrix was calculated by subtracting the two z-score normalised matrices. For feature enrichments (**Figure 5, S4**) whole-chromosome data was considered. For TF motif enrichment analysis, a 5 Mb distance threshold was applied to the differential matrices (**Figure 6**). Cell-type specific contacts were determined for each pair of datasets by fitting a normal distribution to the observed distribution of differential z-scores and selecting contacts with differential intensities stronger than the upper and lower 5% threshold from the fitted curve. To obtain a set of strong and common contacts, differential z-scores within one standard deviation of the mean were sorted by the lower z-score value from the two original datasets and the top 10% of contacts from each chromosome were extracted. Further, for TF motif enrichment analysis only strong cell-type specific contacts with NPMI > mean NPMI of that dataset were selected. All differential contact analyses were limited to chromosomes that were not duplicated as determined by M-FISH and array-CGH (chromosomes 2, 3, 4, 5, 7, 8, 9,10, 13, 17, 18; **Figure S6B**,**C**).

#### Feature enrichment analysis in ES- and XEN-specific contacts by permutation test

For calculation of feature enrichments in differential contacts established by *Lama1, Sox2* and *Gata6* (**Figures 5, S4**), we mapped genomic and epigenomic read data to the NCBI Build 38/mm10 reference genome using Bowtie2 v2.1.0 (Langmead et al., 2009). We excluded replicated reads (i.e., identical reads, mapped to the same genomic location) found more often than the 95th percentile of the frequency distribution of each dataset. We obtained peaks using BCP v1.1 (Xing et al., 2012) in transcription factor mode or histone modification mode with default settings. A full list of published data used in this analysis is given in Table S8. We computed the presence of features for all genomic 40 kb windows using the bedtools (v 2.29.2) (Quinlan and Hall, 2010) window and intersect functions. Genomic windows of interest were then overlapped with windows positive for the feature of interest.

Features were permuted cyclically while preserving feature organisation. Permutation was repeated 10,000 times to create a random genomic background, and for each permutation the overlap of features to the windows of interest was calculated. Empirical p-values for feature depletion or enrichment were calculated against the distribution of feature overlaps in the genomic background.

#### Transcription factor binding motif enrichments in cell-type specific contacts

Enrichments of binding motifs of chosen transcription factors were calculated as described in Winick-Ng et al., 2021 (https://github.com/pombo-lab/WinickNg_Kukalev_Harabula_Nature_2021). Transcription factors were chosen based on differential expression (TPM > 2 in at least one cell type, p-adj of log_2_FC < 1e^-10^) and coverage in windows engaged in cell-type specific interactions (> 20%).

#### Evaluation of small GAM datasets collected from Epi and PrE cells from E4.5 embryos

To quantitatively assess the robustness of contacts detected with smaller GAM datasets, we subsampled a published, high-quality ES-cell dataset, composed of 408 single nuclear slices (Beagrie et al., 2017), by randomly drawing three independent subsets of 59 and 111 single GAM samples (**Figure S6D**). First, we quantified the extent of genomic sampling in the collected and *in silico* subsampled datasets, and found that 99.9% of all 120-kb genomic windows are sampled at least once in the smaller PrE and Epi datasets, and to a similar extent of the full and *in silico* subsampled ES datasets (**Figure S6E**). Second, we measured the co-segregation of all possible intra-chromosomal pairs of genomic windows in the collections of nuclear slices, and found that 86.5% or 93.5% of all possible pairs of 120-kb genomic windows are detected at least once in the Epi and PrE datasets (or 93.5% and 97.8% for 240 kb), to the same extent found in the smaller subsampled ES cell datasets (**Figure S6F**). Slightly higher frequencies of detection of all possible window co-segregations were achieved in the full ES cell dataset samples (99.84 and 99.96% at 120 and 240 kb, respectively). Lastly, to directly compare the contacts obtained from 59 and 111 GAM samples with those obtained from the full dataset of 408 samples, we correlated subsampled with complete contact matrices, and found median correlations of 0.63-0.65 across all chromosomes for 59-sample subsets, and 0.77-0.78 for the 111-sample subsets (for contacts within 5 Mb, at 120 kb resolution; **Figure S6G**). These analyses suggest that GAM datasets with smaller numbers of nuclear slices from Epi and PrE cells have very good sampling quality and capture the broader patterns of chromatin folding at resolutions of 120 kb or 240 kb, which have sampling parameters that are equivalent to the ES and XEN GAM datasets at 20 kb.

#### Gene ontology enrichment analysis of genes in TAD boundaries and contact groups

Gene ontology (GO) enrichments were calculated using GOElite version 1.2.5 (Gladstone Institutes; http://genmapp.org/go_elite). Default parameters were used: z-score threshold > 1.96, permutation-derived p-value < 0.05, number of genes changed > 2. Over-representation analysis (ORA) was performed using the ‘permute p-value’ setting, with 2,000 permutations. For GO analysis, conversion of gene symbols to Ensemble Gene IDs was required, and performed using the KnownToEnsembl table, downloaded from the UCSC Table browser (http://genome.ucsc.edu/cgi-bin/hgTables). Results were filtered for redundancy of terms. For GO enrichment analysis of genes in TAD boundaries in **Figure 3G**, all upregulated genes in the respective cell type were used as background set and terms were chosen from the top 10 most significantly enriched GO Terms, or if more than 10 terms had the lowest p-value, terms were chosen among those with the lowest p-value. For GO enrichment analysis of genes in contact groups of **Figure 6F**,**G**, up-or downregulated genes in the respective cell type were used, depending whether enrichment was calculated for up-or downregulated genes in contacts of interest.

### Total RNA-seq computational analysis

Published and newly produced RNA-seq data was mapped to mouse reference genome mm10 using STAR (Spliced transcript alignment to a reference, v2.6.0c) (Dobin et al., 2013) and processed with RSEM (RNA-Seq by Expectation-Maximisation, v1.3.0) (Li and Dewey, 2011). Genome annotation used was the patch 6 (2017 release) of the Genome Reference Consortium m38 build for improved accuracy (mm10). RefSeq assembly accession GCF_000001635.26 was downloaded from NCBI. RPKM normalised genomic tracks (bigwigs) were generated using the UCSC bamCoverage tool (v3.1.3).

Differential expression analysis was conducted using the DESeq2 (v1.26.0) pipeline (Love et al., 2014). We considered protein coding and long non-coding RNAs, which were detected at over 1 TPM in at least one biological replicate, for differential expression analysis with DESeq2 v1.26.0 (Love et al., 2014). Genes were considered differentially expressed when their adjusted p-value was below 0.05 and their absolute log_2_(fold change) was greater than 1.

Gene ontology enrichments in expression data were calculated using the GSEA pipeline (Gene set enrichment analysis; Subramanian et al., 2005). As input, all differentially expressed genes were ranked according to their p-value (Benjamini-Hochberg multiple testing corrected), and split by the sign of their fold change.

### Single cell RNA-seq data analyses

Single-cell RNA sequencing reads of E4.5 and E7.5 embryos were downloaded from ENA (experiment accessions: SRX5074030, SRX5074031). Single cell reads were aligned to the Gencode vM23/Ensembl 98 mm10 reference with Cellranger v5.0.1. Identifiers of single cells, previously annotated as Epi, PrE or ParE by Nowotschin et al. (2019) were downloaded from www.endoderm-explorer, and used to subset the BAM files, using the 10X Genomics subset-bam v1.0 tool to obtain pseudo-bulk tracks. Gene expression values were obtained by aggregating counts by feature, dividing by the number of counts over all features and multiplying with 100,000. E3.5 and E4.5 scRNA-seq force directed layouts in **Figure 6H-M** were plotted using www.endoderm-explorer.com.

### ATAC-seq computational analysis

Fastq files were mapped to the NCBI build 38/mm10 mouse reference genome using bowtie2 (v2.3.4.3). Reads mapping the mitochondrial genome and low-quality reads (MQ < 30) were filtered out and sequencing file (BAM) was sorted using Sambamba (v0.6.8). Duplicated reads were also removed using Sambamba markdup with default parameters. RPKM normalised genomic tracks (bigwigs) were generated using the UCSC bamCoverage tool (v3.1.3).

ATAC-seq peaks were called using MACS2 (v2.1.1.20160309) (Zhang et al., 2008). To generate a consensus peak dataset among the three biological replicates, a peak file was called using the sequencing files (sorted and deduplicated BAM) for all biological replicates, after merging technical replicates as input.

## Acknowledgements

The authors thank Bettina Purfürst, Minjung Kang, Néstor Saiz for support in adapting block making and staining of E4.5 embryos, Jennifer Giannini for advice on the manuscript, Ibai Irastorza Azcarate, Ehsan Irani and Warren Winick-Ng for advice on computational analysis, Izabela Harabula for support with WGA of embryo samples and all the Pombo lab members for helpful discussions. The authors thank the Protein Purification and Characterisation platform, the Systems Biology Imaging platform and the Genomics platform at the Max-Delbrück-Center for Molecular Medicine in the Helmholtz Association (MDC), Berlin, Germany for technical support and assistance in this work. AP acknowledges support from the Helmholtz Association (Germany), and EU-Horizon International Training Network PEP-NET - 813282 and National Institutes of Health Common Fund 4D Nucleome Program grants U54DK107977. AP and LZR acknowledge support by the Deutsche Forschungsgemeinschaft (DFG; German Research Foundation) International Research Training Group (IRTG2403). GL and RK acknowledge the MDC International PhD Program. TMS and GL acknowledge the MDC-NYU PhD exchange program. SC is a GABBA PhD fellow supported by the FCT (Fundação para a Ciência e Tecnologia; PD/BD/135453/2017). GL acknowledges the Boehringer Ingelheim Fonds travel grant program. Work in AKH’s lab is supported by the NIH (R01HD094868, R01DK127821, R01HD086478, and P30CA008748). LW and YZ acknowledge the Ohio University GERB Program.

## Author contributions

AP, GL and AKH designed the concept for this work; GL and VG collected animal tissues; GL and RK grew cells; GL and RK produced GAM datasets; GL developed the GAM experimental protocol for embryos; GL, AK and CJT developed computational pipelines for bioinformatics and quality control analyses of GAM data; GL performed quality control analyses of GAM data; GL and AK performed bioinformatics analyses of GAM data; DS performed the computational analyses of bulk and single-cell RNA-seq data; GL performed the differential contact analysis, supervised by CJT; GL performed feature enrichment analysis of differential contacts, supervised by CJT; TMS performed ATAC-seq experiments in XEN cells; LZR performed ATAC-seq experiments in ES cells and computational analysis of all ATAC-seq data; GL and SC grew cells for RNA-seq data collection; GL, SC and AK performed RNA-seq experiments; YZ performed TF motif finding enrichments; AKH supervised animal tissue collection and provided animal samples; AP supervised GAM, bulk RNA, ATAC-seq and confocal microscopy experiments and bioinformatics analyses; AP supervised the scRNA-seq analysis; LW supervised the TF motif analyses; MB performed M-FISH and g-banding experiments; AW analysed M-FISH and g-banding experiments, TL supervised M-FISH and g-banding experiments; GL, AP, AKH, VG, DS and LW contributed to the interpretation of the results; GL wrote the first draft of the manuscript and designed the figures; GL, AP and AKH wrote the manuscript. All authors provided critical feedback and helped revise the manuscript.

## Supplementary table titles and legends

**Table S1**. Expression in all replicates of ES and XEN cells. Differential gene expression between the two cell types, where positive log2(fold change) indicates an upregulation in XEN cells, and negative values and upregulation in ES cells. Genes were ranked by the logarithmized adjusted p-value and sign of the fold change. Gene set enrichment analysis on the ranked list was performed with GSEA. Gene sets enriched at the top of the ranked list (higher expression in ES cells) are displayed on sheet 3, gene sets enriched at the bottom of the ranked list (higher expression in XEN cells) are displayed on sheet 4. Related to Figure 3.

**Table S2**. Coordinates for TAD boundaries in ES and XEN cells. Boundaries were called with an adaptation of the insulation square method (Crane et al., 2015) as described in methods, at 40 kb resolution with a square size of 400 kb. Sheets with GO enrichment analysis describe GO terms for genes in cell-type-specific boundaries that are also upregulated in that cell type. Analysis was done using GOElite, with standard parameters, as indicated in methods. Background gene sets for enrichment were all expressed genes (> 1 TPM) in the respective cell type. Related to Figure 3.

**Table S3**. Master table for all genes, showing TAD boundary presence, compartment A/B status, expression and differential expression. Related to Figure 3.

**Table S4**. Compartment states and Eigenvalues of the first principal component in ES and XEN cells, for all 250 kb windows in the genome. Related to Figure 4.

**Table S5**. TF motif feature pair analysis with differential expression analysis for XEN and ES cells. Related to Figure 6.

**Table S6**. List of GO terms enriched in cell type-specific contacts, containing accessible binding sites for specific TF pairs. Related to Figure 6.

**Table S7**. Experimental, sequencing and QC metrics for every sample in GAM dataset on XEN cells (clone IM8A1) or PrE and Epi cells from mouse blastocyst E4.5. Related to STAR methods.

**Table S8**. Overview and availability of published data used in this study. Related to STAR methods.

