## Supplementary material for "3D genome topologies distinguish pluripotent epiblast and primitive endoderm cells in the mouse blastocyst": SI Data File

### Aberration Summary Report

#### Sample Properties

| | | | |
|---|---|---|---|
| Array type | : | ArraySet | : |
| Red Sample | : XEN R1 | Green Sample | : |

#### Analysis Settings

| | |
|---|---|
| Genome | : mm9 |
| Aberration Algorithm | : ADM-2 |
| Threshold | : 6.0 |
| Fuzzy Zero | : ON |
| GC Correction | : ON |
| Window Size | : 2Kb |
| Centralization (legacy) | : OFF |
| Diploid Peak Centralization | : ON |
| Manually Reassign Peaks | : OFF |
| Combine Replicates (Intra Array) | : ON |
| Feature Level Filter | : glsSaturated = true OR rIsSaturated = true OR glsFeatNonUnifOL = true OR rIsFeatNonUnifOL = true OR LogRatio = 0 |
| Design Level Filter | : Homology = 0 OR IsPseudoautosomal = 1 |
| Genomic Boundary | : OFF |
| Aberration Filter | : ( Minimum Number of Probes for Amplification >= 3 AND Minimum Size (Kb) of Region for Amplification >= 0.0 AND Minimum Avg. Absolute Log Ratio for Amplification >= 0.25 ) OR ( Minimum Number of Probes for Deletion >= 3 AND Minimum Size (Kb) of Region for Deletion >= 0.0 AND Minimum Avg. Absolute Log Ratio for Deletion >= 0.25 ) |

#### Genome Overview

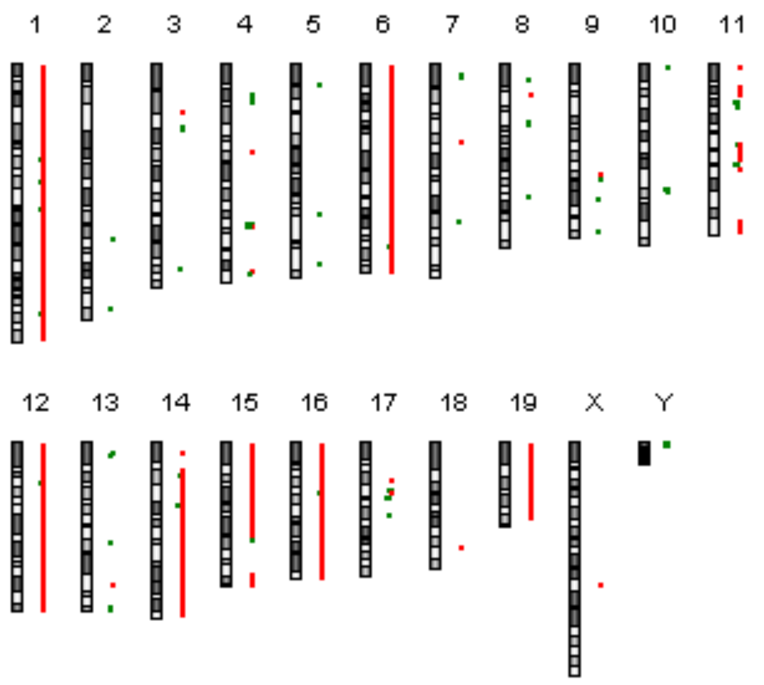

#### Comments

Technician: Date:__/__/__ Supervisor Date:__/__/__

#### Text Summary Report for Sample US81403230_252741111547_S01_CGH_107_Sep09_SureTag_1_1

| Event No | Chr | Cytoband | #Probes | Amp/Del | P-value | Annotations |
|---|---|---|---|---|---|---|
| 1 | chr1:3192257-34606839 | qA1 - qB | 1898 | 0.445036 | NA | Xkr4, Rp1, Sox17... |
| 2 | chr1:34606840-34639821 | qB | 5 | -0.254251 | NA | Tesp2, Fam123c |
| 3 | chr1:34639822-66327175 | qB - qC3 | 2137 | 0.445036 | NA | Fam123c, Arhgef4, Fam168b... |
| 4 | chr1:66327176-66341987 | qC3 | 3 | -1.056267 | NA | Mtap2 |
| 5 | chr1:66341988-67682928 | qC3 | 103 | 0.445036 | NA | Mtap2, Unc80, Rpe... |
| 6 | chr1:67682929-67817197 | qC3 | 4 | -0.849167 | NA | |
| 7 | chr1:67817198-82907979 | qC3 - qC5 | 1022 | 0.445036 | NA | Erbb4, Ikzf2, Spag16... |
| 8 | chr1:82907980-82938344 | qC5 | 3 | -0.610730 | NA | |
| 9 | chr1:82938345-101783951 | qC5 - qE1.1 | 1136 | 0.445036 | NA | Slc19a3, Ccl20, Wdr69... |
| 10 | chr1:101783952-101807547 | qE1.1 | 4 | -0.502192 | NA | Cntnap5b |
| 11 | chr1:101807548-141582303 | qE1.1 - qF | 2354 | 0.445036 | NA | Cntnap5b, Cdh20, Rnf152... |
| 12 | chr1:141582304-141678786 | qF | 9 | -0.409605 | NA | EG214403 |
| 13 | chr1:141678787-173457379 | qF - qH3 | 2076 | 0.445036 | NA | BC026782, Cfh, Kcnt2... |
| 14 | chr1:173457380-173490445 | qH3 | 3 | 1.867157 | NA | Itln1, Cd244 |
| 15 | chr1:173490446-175715422 | qH3 | 183 | 0.445036 | NA | Cd244, Ly9, Slamf7... |
| 16 | chr1:175715423-175796142 | qH3 | 6 | -0.890102 | NA | |
| 17 | chr1:175796143-176330988 | qH3 | 41 | 0.445036 | NA | Mnda, Ifi203, Ifi202b... |
| 18 | chr1:176330989-176369910 | qH3 | 6 | -0.423652 | NA | Olfr414 |
| 19 | chr1:176369911-197141706 | qH3 - qH6 | 1482 | 0.445036 | NA | Fmn2, Grem2, Rgs7... |
| 20 | chr2:122573363-123648375 | qE5 | 36 | -0.284777 | NA | Pldn, Sqrdl |
| 21 | chr2:172762838-172797294 | qH3 | 4 | -1.179027 | NA | Bmp7 |
| 22 | chr3:33423132-36370782 | qA3 - qB | 161 | 0.361692 | NA | Ttc14, Ccdc39, Fxr1... |
| 23 | chr3:43932117-49397682 | qB - qC | 181 | -0.398144 | NA | Pcdh10, Pabpc4l, 1700018B24Rik |
| 24 | chr3:144424171-144490871 | qH2 | 6 | -0.738190 | NA | Clca2, Clca4 |
| 25 | chr4:22522479-30426687 | qA3 - qA5 | 341 | -0.301657 | NA | F730047E07Rik, Klhl32, Ndufaf4... |
| 26 | chr4:62164667-62182581 | qB3 | 3 | 1.568056 | NA | Alad |
| 27 | chr4:111726014-111771908 | qD1 | 4 | -2.096744 | NA | Skint4 |
| 28 | chr4:111771909-112147017 | qD1 | 30 | -4.014340 | NA | Skint4, Skint3, Skint9 |
| 29 | chr4:112147018-112202834 | qD1 | 1 | -2.096744 | NA | |
| 30 | chr4:112202835-112273763 | qD1 | 4 | -4.092874 | NA | |
| 31 | chr4:112273764-112289417 | qD1 | 1 | -2.096744 | NA | Skint2 |
| 32 | chr4:112289418-112460956 | qD1 | 14 | -0.311672 | NA | Skint2, Skint10 |
| 33 | chr4:112460957-112480583 | qD1 | 1 | -2.096744 | NA | Skint6 |
| 34 | chr4:112480584-112504314 | qD1 | 4 | -1.114290 | NA | Skint6 |
| 35 | chr4:112504315-112525047 | qD1 | 1 | -2.096744 | NA | Skint6 |
| 36 | chr4:112525048-112564300 | qD1 | 5 | -4.428579 | NA | Skint6 |
| 37 | chr4:112564301-112664999 | qD1 | 6 | -2.096744 | NA | Skint6 |
| 38 | chr4:112665000-112785680 | qD1 | 11 | -4.256351 | NA | Skint6 |
| 39 | chr4:112785681-113251939 | qD1 | 26 | -3.221405 | NA | Skint6 |

| Event No | Chr | Cytoband | #Probes | Amp/Del | P-value | Annotations |
|---|---|---|---|---|---|---|
| 40 | chr4:113286238-113453826 | qD1 | 9 | -0.857491 | NA | |
| 41 | chr4:113453827-113491779 | qD1 | 3 | 0.261225 | NA | |
| 42 | chr4:113491780-113518070 | qD1 | 1 | -0.857491 | NA | |
| 43 | chr4:113518071-113549532 | qD1 | 3 | -2.696584 | NA | |
| 44 | chr4:113549533-113574763 | qD1 | 2 | -0.857491 | NA | |
| 45 | chr4:145018403-145192926 | qE1 | 8 | 0.978098 | NA | Vmn2r-ps14 |
| 46 | chr4:146804312-146935370 | qE1 | 7 | -0.603303 | NA | Gm13152, Gm13154 |
| 47 | chr5:15684053-15872543 | qA2 | 21 | -0.965443 | NA | Cacna2d1 |
| 48 | chr5:105142362-105202268 | qE5 | 4 | -0.905103 | NA | |
| 49 | chr5:140239345-140265361 | qG2 | 4 | -0.706556 | NA | Ints1 |
| 50 | chr6:3131098-3400603 | qA1 | 9 | 0.554883 | NA | Samd9l |
| 51 | chr6:3400604-34506915 | qA1 - qB1 | 2033 | 0.748930 | NA | Hepacam2, Ccdc132, Calcr... |
| 52 | chr6:34506916-34541940 | qB1 | 3 | -0.279601 | NA | |
| 53 | chr6:34541941-52841579 | qB1 - qB3 | 1422 | 0.748930 | NA | Cald1, Agbl3, Tmem140... |
| 54 | chr6:52841580-70150127 | qB3 - qC1 | 1171 | 0.554883 | NA | Jazf1, Creb5, 1200009O22Rik... |
| 55 | chr6:70150128-70264552 | qC1 | 4 | 1.381556 | NA | |
| 56 | chr6:70264553-128726560 | qC1 - qF3 | 4147 | 0.554883 | NA | Rpia, Eif2ak3, 1700011F03Rik... |
| 57 | chr6:128726561-128746179 | qF3 | 3 | -1.126642 | NA | Klrb1c |
| 58 | chr6:128746180-149494692 | qF3 - qG3 | 1445 | 0.554883 | NA | Klrb1b, Clec2i, Clec2g... |
| 59 | chr7:7089757-12236949 | qA1 | 62 | -0.311757 | NA | Zfp418, BC023179, Vmn2r29... |
| 60 | chr7:54844755-54883844 | qB4 | 3 | 1.014187 | NA | Mrgpra3 |
| 61 | chr7:110691909-110709040 | qE3 | 3 | -1.016216 | NA | Olfr614, Olfr615 |
| 62 | chr7:111656562-111683983 | qE3 | 3 | -1.016724 | NA | Gm6577 |
| 63 | chr8:11316586-11382229 | qA1.1 | 9 | -1.128457 | NA | Col4a2 |
| 64 | chr8:22113295-22233176 | qA2 | 5 | 1.099103 | NA | Defa21, Defa-rs7, Defa23... |
| 65 | chr8:40187843-40274870 | qA4 | 6 | -0.578776 | NA | Tusc3 |
| 66 | chr8:43127913-43149700 | qA4 | 3 | -1.200864 | NA | |
| 67 | chr8:93508957-93530058 | qC5 | 3 | -1.018710 | NA | Chd9 |
| 68 | chr9:78512136-78574163 | qE1 | 8 | 0.493691 | NA | Cd109 |
| 69 | chr9:81761824-83221489 | qE1 - qE2 | 75 | -0.323158 | NA | 4930486G11Rik, Irak1bp1, Phip... |
| 70 | chr9:95556843-95572710 | qE3.3 | 3 | -1.046355 | NA | Pcolce2 |
| 71 | chr9:117877147-117905398 | qF3 | 4 | -0.862698 | NA | |
| 72 | chr10:3143971-3173641 | qA1 | 5 | -0.713938 | NA | Cnksr3 |
| 73 | chr10:88379728-88397518 | qC1 | 3 | -2.137453 | NA | Slc5a8 |
| 74 | chr10:90492952-90513111 | qC2 | 3 | -1.151012 | NA | Apaf1 |
| 75 | chr11:3107005-3567716 | qA1 | 40 | 0.661151 | NA | Eif4enif1, Drg1, Patz1... |
| 76 | chr11:15932992-17933933 | qA2 - qA3.1 | 108 | 0.361972 | NA | Vstm2a, Sec61g, Egfr... |
| 77 | chr11:17988148-21712069 | qA3.1 | 209 | 0.808196 | NA | Meis1, Spred2, Actr2... |
| 78 | chr11:21712070-21877950 | qA3.1 - qA3.2 | 12 | 0.374159 | NA | AV249152 |
| 79 | chr11:21877951-21932468 | qA3.2 | 4 | 0.808196 | NA | Otx1, Ehbp1 |
| 80 | chr11:27574653-27632740 | qA3.3 | 3 | -3.886607 | NA | |
| 81 | chr11:30540062-30582473 | qA4 | 5 | -0.979768 | NA | Acyp2 |
| 82 | chr11:31049571-31227838 | qA4 | 5 | -0.926768 | NA | |
| 83 | chr11:57072524-57096542 | qB1.3 | 4 | -0.578572 | NA | Gria1 |
| 84 | chr11:57384950-57578028 | qB1.3 | 14 | 0.687359 | NA | Galnt10 |
| 85 | chr11:57578029-58533135 | qB1.3 | 83 | 0.404437 | NA | Galnt10, Sap30l, Hand1... |
| 86 | chr11:58533136-58660441 | qB1.3 | 13 | 0.687359 | NA | Olfr318, Olfr317, Olfr316... |
| 87 | chr11:58660442-62245721 | qB1.3 - qB2 | 330 | 0.927527 | NA | 2810021J22Rik, Zfp39, Butr1... |
| 88 | chr11:62245722-64526710 | qB2 - qB3 | 161 | 0.687359 | NA | Ncor1, Pigl, Cenpv... |
| 89 | chr11:64526711-67055280 | qB3 | 179 | 0.469273 | NA | Elac2, AU040829, Myocd... |
| 90 | chr11:67055281-67286644 | qB3 | 20 | 0.687359 | NA | Myh4, Myh8, Myh13 |
| 91 | chr11:70989862-71107103 | qB4 | 11 | -3.306733 | NA | Nlrp1b, Nlrp1c |
| 92 | chr11:74184906-74199333 | qB5 | 3 | 1.073781 | NA | Garnl4 |
| 93 | chr11:110879483-111518645 | qE2 | 24 | 0.303829 | NA | Kcnj16, Kcnj2 |

| Event No | Chr | Cytoband | #Probes | Amp/Del | P-value | Annotations |
|---|---|---|---|---|---|---|
| 94 | chr11:113062032-121797041 | qE2 | 797 | 0.477446 | NA | Slc39a11, Sstr2, Cog1... |
| 95 | chr12:3162616-16354595 | qA1.1 | 732 | 0.463887 | NA | Rab10, Kif3c, Asxl2... |
| 96 | chr12:16354596-16528555 | qA1.1 | 5 | -0.422117 | NA | |
| 97 | chr12:16528556-28429346 | qA1.1 - qA2 | 320 | 0.463887 | NA | Lpin1, Ntsr2, Greb1... |
| 98 | chr12:28429347-28483451 | qA2 | 3 | -1.718620 | NA | |
| 99 | chr12:28483452-114670168 | qA2 - qF1 | 5643 | 0.463887 | NA | Allc, Colec11, Rps7... |
| 100 | chr12:114670169-115298715 | qF1 | 48 | 0.881950 | NA | Adam6b, Adam6a |
| 101 | chr12:115298716-116840010 | qF1 - qF2 | 85 | 0.463887 | NA | |
| 102 | chr12:116840011-117047397 | qF2 | 7 | 1.423474 | NA | |
| 103 | chr12:117047398-121221931 | qF2 | 291 | 0.463887 | NA | Zfp386, Vipr2, Wdr60... |
| 104 | chr13:7593763-7910087 | qA1 | 20 | -0.441211 | NA | |
| 105 | chr13:8915529-8959470 | qA1 | 3 | -0.994944 | NA | Idi2 |
| 106 | chr13:9326106-9346851 | qA1 | 3 | -1.664423 | NA | Dip2c |
| 107 | chr13:70119197-70195757 | qC1 | 5 | -0.779343 | NA | |
| 108 | chr13:101110785-101183591 | qD1 | 6 | 0.789514 | NA | Naip1 |
| 109 | chr13:115711622-115732085 | qD2.2 | 3 | -0.998802 | NA | Itga2 |
| 110 | chr13:118460047-118476029 | qD2.3 | 3 | -1.113086 | NA | Hcn1 |
| 111 | chr14:8572189-10278956 | qA1 | 118 | 0.453607 | NA | Flnb, Dnase1l3, Abhd6... |
| 112 | chr14:20445376-23364371 | qA3 | 296 | 0.402276 | NA | Nid2, 2700060E02Rik, Gng2... |
| 113 | chr14:23364372-23383481 | qA3 | 3 | -0.532767 | NA | 1700112E06Rik |
| 114 | chr14:23383482-23726436 | qA3 | 39 | 0.402276 | NA | 1700112E06Rik |
| 115 | chr14:23726437-23748305 | qA3 | 4 | -0.536554 | NA | 1700112E06Rik |
| 116 | chr14:23748306-44542852 | qA3 - qC1 | 1409 | 0.402276 | NA | 1700112E06Rik, Kcnma1, Dlg5... |
| 117 | chr14:44542853-44579789 | qC1 | 6 | -2.039362 | NA | Ang6 |
| 118 | chr14:44579790-125143137 | qC1 - qE5 | 5237 | 0.402276 | NA | Ang6, Ear2, Ear12... |
| 119 | chr15:3114799-35590381 | qA1 - qB3.1 | 1938 | 0.644631 | NA | Sepp1, Ghr, Fbxo4... |
| 120 | chr15:35590382-35711964 | qB3.1 | 13 | 1.152264 | NA | Vps13b |
| 121 | chr15:35711965-36017337 | qB3.1 | 27 | 0.644631 | NA | Vps13b, Cox6c |
| 122 | chr15:36017338-36108577 | qB3.1 | 8 | 1.198952 | NA | Fbxo43, Polr2k |
| 123 | chr15:36108578-36181161 | qB3.1 | 7 | 0.644631 | NA | Spag1, Rnf19a |
| 124 | chr15:36181162-41292216 | qB3.1 | 367 | 1.162520 | NA | Rnf19a, Ankrd46, Snx31... |
| 125 | chr15:41292217-51624140 | qB3.1 - qC | 528 | 0.644631 | NA | Oxr1, Abra, Angpt1... |
| 126 | chr15:51624141-68932446 | qC - qD3 | 1059 | 0.844171 | NA | Eif3h, Utp23, Rad21... |
| 127 | chr15:68932447-69378937 | qD3 | 14 | 0.644631 | NA | |
| 128 | chr15:69400090-70779311 | qD3 | 43 | -0.313395 | NA | |
| 129 | chr15:93512848-103432826 | qE3 - qF3 | 860 | 0.571316 | NA | Adamts20, Pus7l, Irak4... |
| 130 | chr16:3284525-6149147 | qA1 | 214 | 0.531833 | NA | Olfr161, Mefv, Zfp263... |
| 131 | chr16:6149148-6168655 | qA1 | 3 | -0.255792 | NA | A2bp1 |
| 132 | chr16:6168656-36245660 | qA1 - qB3 | 2338 | 0.531833 | NA | A2bp1, Tmem114, BC024814... |
| 133 | chr16:36245661-36336077 | qB3 | 7 | -0.837443 | NA | 2010005H15Rik, Stfa1, BC117090 |
| 134 | chr16:36336078-44835804 | qB3 - qB4 | 651 | 0.531833 | NA | BC100530, Stfa2, Stfa3... |
| 135 | chr16:44835805-44872692 | qB4 | 4 | -0.444304 | NA | Cd200r4, Cd200r2 |
| 136 | chr16:44872693-98242899 | qB4 - qC4 | 3048 | 0.531833 | NA | Cd200r2, Cd200r3, Ccdc80... |
| 137 | chr17:27451543-27582091 | qA3.3 | 7 | 0.865584 | NA | Grm4 |
| 138 | chr17:33297519-33319404 | qB1 | 3 | -1.821836 | NA | Olfr55 |
| 139 | chr17:34117148-34494339 | qB1 | 38 | -0.345732 | NA | AA388235, H2-K1, Ring1... |
| 140 | chr17:35387186-35477935 | qB1 | 9 | 0.773480 | NA | Bat1a, H2-D1, LOC547349... |
| 141 | chr17:38449974-38501757 | qB1 | 5 | -3.078815 | NA | Olfr136 |
| 142 | chr17:52145888-52211123 | qC | 6 | -0.814203 | NA | |
| 143 | chr18:73767484-73976871 | qE2 | 22 | 0.606166 | NA | Smad4, Elac1, Me2 |
| 144 | chr19:3260956-39989067 | qA - qC3 | 2801 | 0.606183 | NA | Ighmbp2, Mrpl21, Cpt1a... |
| 145 | chr19:39989068-50318633 | qC3 - qD1 | 889 | 0.902154 | NA | Cyp2c37, Cyp2c54, Cyp2c50... |
| 146 | chr19:50318634-51030714 | qD1 | 56 | 0.606183 | NA | Sorcs1 |
| 147 | chr19:51030715-53393768 | qD1 - qD2 | 92 | 0.876968 | NA | Ins1, Xpnpep1, Add3... |

| Event No | Chr | Cytoband | #Probes | Amp/Del | P-value | Annotations |
|---|---|---|---|---|---|---|
| 148 | chr19:53393769-53401312 | qD2 | 1 | 0.606183 | NA | Mxi1 |
| 149 | chrX:100008370-100023414 | qD | 3 | 1.069696 | NA | |
| 150 | chrY:37653-139871 | qA1 | 8 | -1.848036 | NA | Zfy1 |
| 151 | chrY:139872-249095 | qA1 | 1 | -2.930083 | NA | Ube1y1, Kdm5d |
| 152 | chrY:249096-306546 | qA1 | 8 | -4.236148 | NA | Kdm5d |
| 153 | chrY:306547-346877 | qA1 | 2 | -2.930083 | NA | |
| 154 | chrY:346878-374359 | qA1 | 5 | -4.702396 | NA | Eif2s3y |
| 155 | chrY:374360-434832 | qA1 | 3 | -2.930083 | NA | Tspy-ps, Uty |
| 156 | chrY:434833-520092 | qA1 | 12 | -3.710355 | NA | Uty |
| 157 | chrY:520093-535058 | qA1 | 3 | -2.178633 | NA | Uty |
| 158 | chrY:535059-581689 | qA1 | 6 | -3.710355 | NA | Uty |
| 159 | chrY:581690-602081 | qA1 | 3 | -6.257580 | NA | Uty, Ddx3y |
| 160 | chrY:602082-731591 | qA1 | 16 | -3.710355 | NA | Ddx3y, Usp9y |
| 161 | chrY:731592-801911 | qA1 | 9 | -2.930083 | NA | Usp9y |
| 162 | chrY:801912-1365243 | qA1 | 14 | -4.439327 | NA | Zfy2 |
| 163 | chrY:1365244-1707093 | qA1 | 16 | -2.930083 | NA | Zfy2 |
| 164 | chrY:1707094-1824601 | qA1 | 4 | -5.043808 | NA | |
| 165 | chrY:1824602-2506308 | qA1 | 6 | -2.930083 | NA | Gm16501, Sry, Rbmy1a1 |
| 166 | chrY:2506309-2546621 | qA1 | 3 | -1.437846 | NA | |
| 167 | chrY:2546622-2635638 | qA1 | 3 | -2.930083 | NA | |

Amp=Amplification
Del=Deletion

### Aberration Summary Report

#### Sample Properties

| | | | |
|---|---|---|---|
| Array type | : | ArraySet | : |
| Red Sample | : XEN R2 | Green Sample | : |

#### Analysis Settings

| | |
|---|---|
| Genome | : mm9 |
| Aberration Algorithm | : ADM-2 |
| Threshold | : 6.0 |
| Fuzzy Zero | : ON |
| GC Correction | : ON |
| Window Size | : 2Kb |
| Centralization (legacy) | : OFF |
| Diploid Peak Centralization | : ON |
| Manually Reassign Peaks | : OFF |
| Combine Replicates (Intra Array) | : ON |
| Feature Level Filter | : glsSaturated = true OR rIsSaturated = true OR glsFeatNonUnifOL = true OR rIsFeatNonUnifOL = true OR LogRatio = 0 |
| Design Level Filter | : Homology = 0 OR IsPseudoautosomal = 1 |
| Genomic Boundary | : OFF |
| Aberration Filter | : ( Minimum Number of Probes for Amplification >= 3 AND Minimum Size (Kb) of Region for Amplification >= 0.0 AND Minimum Avg. Absolute Log Ratio for Amplification >= 0.25 ) OR ( Minimum Number of Probes for Deletion >= 3 AND Minimum Size (Kb) of Region for Deletion >= 0.0 AND Minimum Avg. Absolute Log Ratio for Deletion >= 0.25 ) |

#### Genome Overview

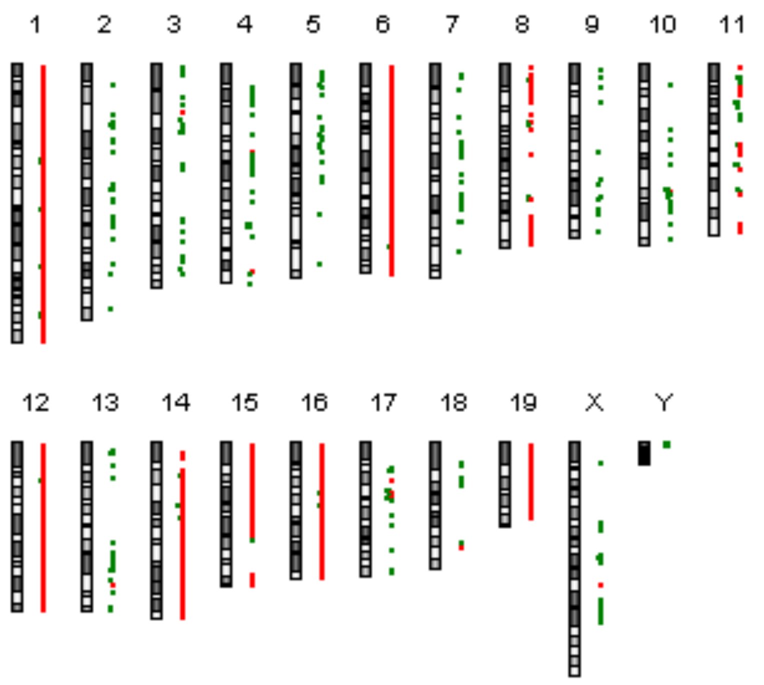

#### Comments

Technician: Date:__/__/__ Supervisor Date:__/__/__

#### Text Summary Report for Sample US81403230_252741111547_S01_CGH_107_Sep09_SureTag_1_2

| Event No | Chr | Cytoband | #Probes | Amp/Del | P-value | Annotations |
|---|---|---|---|---|---|---|
| 1 | chr1:3192257-4399504 | qA1 | 70 | 0.398807 | NA | Xkr4, Rp1 |
| 2 | chr1:4399505-5428296 | qA1 | 67 | 0.571347 | NA | Sox17, Mrpl15, Lypla1... |
| 3 | chr1:5428297-9701171 | qA1 - qA2 | 240 | 0.398807 | NA | Oprk1, Npbwr1, Rb1cc1... |
| 4 | chr1:9701172-10212198 | qA2 | 54 | 0.642228 | NA | Vcpip1, Sgk3, 6030422M02Rik... |
| 5 | chr1:10212199-33521532 | qA2 - qB | 1364 | 0.398807 | NA | Arfgef1, Cpa6, Prex2... |
| 6 | chr1:33521533-34606839 | qB | 104 | 0.546802 | NA | Prim2, Rab23, Bag2... |
| 7 | chr1:34606840-34639821 | qB | 5 | -0.274618 | NA | Tesp2, Fam123c |
| 8 | chr1:34639822-38205065 | qB | 255 | 0.546802 | NA | Fam123c, Arhgef4, Fam168b... |
| 9 | chr1:38205066-42915177 | qB | 298 | 0.398807 | NA | Aff3, Lonrf2, Chst10... |
| 10 | chr1:42915178-44213429 | qB - qC1.1 | 98 | 0.563811 | NA | Mrps9, Gpr45, Tgfbrap1... |
| 11 | chr1:44213430-44479791 | qC1.1 | 11 | 0.398807 | NA | Ercc5, 4832428D23Rik |
| 12 | chr1:44479792-50968806 | qC1.1 | 316 | 0.281847 | NA | Gulp1, Col3a1, Col5a2... |
| 13 | chr1:50968807-51807078 | qC1.1 | 60 | 0.398807 | NA | Tmeff2, Sdpr, Obfc2a... |
| 14 | chr1:51807079-65425886 | qC1.1 - qC2 | 1063 | 0.483546 | NA | Myo1b, Stat4, Stat1... |
| 15 | chr1:65425887-66327175 | qC2 - qC3 | 37 | 0.398807 | NA | Pth2r, Crygf, Mtap2 |
| 16 | chr1:66327176-66341987 | qC3 | 3 | -1.242175 | NA | Mtap2 |
| 17 | chr1:66341988-66422996 | qC3 | 8 | 0.398807 | NA | Mtap2 |
| 18 | chr1:66422997-67242362 | qC3 | 78 | 0.298884 | NA | Mtap2, Unc80, Rpe... |
| 19 | chr1:67242363-67682928 | qC3 | 17 | 0.398807 | NA | Cps1 |
| 20 | chr1:67682929-67817197 | qC3 | 4 | -0.883460 | NA | |
| 21 | chr1:67817198-68366731 | qC3 | 35 | 0.398807 | NA | Erbb4 |
| 22 | chr1:68366732-68442987 | qC3 | 8 | -0.474544 | NA | Erbb4 |
| 23 | chr1:68442988-71462737 | qC3 | 250 | 0.398807 | NA | Erbb4, Ikzf2, Spag16... |
| 24 | chr1:71462738-75727412 | qC3 - qC4 | 329 | 0.563828 | NA | Atic, Fn1, Gm8883... |
| 25 | chr1:75727413-82293232 | qC4 - qC5 | 338 | 0.398807 | NA | Epha4, Pax3, Sgpp2... |
| 26 | chr1:82293233-82883180 | qC5 | 61 | 0.590390 | NA | Rhbdd1, Col4a4, Col4a3... |
| 27 | chr1:82883181-87670433 | qC5 | 170 | 0.398807 | NA | Agfg1, Slc19a3, Ccl20... |
| 28 | chr1:87670434-96276975 | qC5 - qD | 704 | 0.527483 | NA | Cab39, Itm2c, Gpr55... |
| 29 | chr1:96276976-99520288 | qD | 124 | 0.398807 | NA | Fam174a, St8sia4, Slco4c1... |
| 30 | chr1:99520289-100088911 | qD | 53 | 0.359928 | NA | D1Ertd622e, Hisppd1, Zh2c2... |
| 31 | chr1:100088912-101783951 | qD - qE1.1 | 89 | 0.398807 | NA | Cntnap5b |
| 32 | chr1:101783952-101807547 | qE1.1 | 4 | -0.802121 | NA | Cntnap5b |
| 33 | chr1:101807548-106752971 | qE1.1 - qE2.1 | 198 | 0.398807 | NA | Cntnap5b, Cdh20 |
| 34 | chr1:106752972-108915126 | qE2.1 | 160 | 0.410400 | NA | Cdh20, Rnf152, Pign... |
| 35 | chr1:108915127-120196860 | qE2.1 - qE2.3 | 415 | 0.398807 | NA | Serpinb3a, Serpinb3d, Serpinb3b... |
| 36 | chr1:120196861-121730969 | qE2.3 | 127 | 0.572933 | NA | Tsn, Mki67ip, Clasp1... |
| 37 | chr1:121730970-129786859 | qE2.3 - qE3 | 514 | 0.398807 | NA | Ptpn4, Tmem177, Gm101... |
| 38 | chr1:129786860-130209706 | qE3 - qE4 | 45 | 0.607139 | NA | Rab3gap1, Zranb3, R3hdm1... |
| 39 | chr1:130209707-133814573 | qE4 | 275 | 0.398807 | NA | Lct, Mcm6, Dars... |
| 40 | chr1:133814574-139249649 | qE4 | 471 | 0.516799 | NA | Nucks1, Slc45a3, Elk4... |

| Event No | Chr | Cytoband | #Probes | Amp/Del | P-value | Annotations |
|---|---|---|---|---|---|---|
| 41 | chr1:139249650-141582303 | qE4 - qF | 151 | 0.398807 | NA | Ptprc, Atp6v1g3, Nek7... |
| 42 | chr1:141582304-141678786 | qF | 9 | -0.649900 | NA | EG214403 |
| 43 | chr1:141678787-145084897 | qF | 136 | 0.398807 | NA | BC026782, Cfh, Kcnt2 |
| 44 | chr1:145084898-145972385 | qF | 49 | 0.494918 | NA | Cdc73, B3galt2, Glrx2... |
| 45 | chr1:145972386-152209584 | qF - qG1 | 246 | 0.398807 | NA | Rgs13, Rgs1, Rgs18... |
| 46 | chr1:152209585-159533067 | qG1 - qH1 | 597 | 0.485097 | NA | BC003331, Tpr, Prg4... |
| 47 | chr1:159533068-161172087 | qH1 | 104 | 0.307765 | NA | Fam5b, Astn1, Pappa2... |
| 48 | chr1:161172088-173457379 | qH1 - qH3 | 948 | 0.485097 | NA | Rfwd2, Tnr, 4930523C07Rik... |
| 49 | chr1:173457380-173490445 | qH3 | 3 | 1.827782 | NA | Itln1, Cd244 |
| 50 | chr1:173490446-174399001 | qH3 | 85 | 0.485097 | NA | Cd244, Ly9, Slamf7... |
| 51 | chr1:174399002-175197600 | qH3 | 62 | 0.398807 | NA | Slamf9, Igsf9, Tagln2... |
| 52 | chr1:175197601-175253384 | qH3 | 5 | 0.785997 | NA | Olfr1403 |
| 53 | chr1:175253385-175715422 | qH3 | 32 | 0.398807 | NA | Darc, Cadm3, Aim2... |
| 54 | chr1:175715423-175796142 | qH3 | 6 | -0.976640 | NA | |
| 55 | chr1:175796143-176330988 | qH3 | 41 | 0.398807 | NA | Mnda, Ifi203, Ifi202b... |
| 56 | chr1:176330989-176369910 | qH3 | 6 | -0.568452 | NA | Olfr414 |
| 57 | chr1:176369911-178642395 | qH3 - qH4 | 181 | 0.398807 | NA | Fmn2, Grem2, Rgs7... |
| 58 | chr1:178642396-185579220 | qH4 - qH5 | 535 | 0.518644 | NA | Cep170, Sdccag8, Akt3... |
| 59 | chr1:185579221-186615069 | qH5 | 46 | 0.326262 | NA | Dusp10, Hlx, Mosc1 |
| 60 | chr1:186615070-195211037 | qH5 - qH6 | 629 | 0.518644 | NA | Mosc1, Mosc2, C130074G19Rik... |
| 61 | chr1:195211038-197141706 | qH6 | 93 | 0.398807 | NA | Plxna2, Cd34, Cd46... |
| 62 | chr2:15419962-16681542 | qA2 - qA3 | 68 | -0.297373 | NA | Plxdc2 |
| 63 | chr2:36632200-37035082 | qB | 31 | -0.262221 | NA | Olfr348, Olfr50, Olfr3... |
| 64 | chr2:40338308-41998467 | qB | 155 | -0.334850 | NA | Lrp1b |
| 65 | chr2:41998468-42793923 | qB | 65 | -0.517561 | NA | Lrp1b |
| 66 | chr2:42793924-48588194 | qB - qC1.1 | 279 | -0.334850 | NA | Kynu, Arhgap15, Gtdc1... |
| 67 | chr2:53267193-57876956 | qC1.1 | 215 | -0.272814 | NA | Rprm, Galnt13, Kcnj3... |
| 68 | chr2:62511116-63901532 | qC1.3 | 89 | -0.351281 | NA | Gca, Kcnh7, Fign |
| 69 | chr2:85474534-87646245 | qD | 203 | -0.283583 | NA | Olfr1002, Olfr154, Olfr1006... |
| 70 | chr2:87646246-87715562 | qD | 9 | -0.758228 | NA | Olfr1145, Olfr1148, Olfr1151... |
| 71 | chr2:87715563-89361730 | qD - qE1 | 146 | -0.283583 | NA | Olfr1153, Olfr1154, Olfr1155... |
| 72 | chr2:94617766-100925085 | qE1 | 241 | -0.384044 | NA | Lrrc4c |
| 73 | chr2:106920480-108044707 | qE3 | 34 | -0.368873 | NA | Kcna4 |
| 74 | chr2:110453088-111608853 | qE3 | 93 | -0.287613 | NA | Slc5a12, Ano3, Muc15... |
| 75 | chr2:113987129-115123590 | qE4 | 58 | -0.332060 | NA | Aqr, Zfp770, Atpbd4 |
| 76 | chr2:122422995-122519791 | qE5 | 11 | 0.433521 | NA | Gatm, AA467197, Slc30a4 |
| 77 | chr2:122564835-123648375 | qE5 | 37 | -0.380736 | NA | Pldn, Sqrdl |
| 78 | chr2:141242323-141802072 | qG1 | 60 | -0.332324 | NA | Macrod2 |
| 79 | chr2:146839968-146859942 | qG2 | 3 | -0.815058 | NA | Xrn2 |
| 80 | chr2:172762838-172797294 | qH3 | 4 | -1.175991 | NA | Bmp7 |
| 81 | chr3:3000029-7509145 | qA1 | 168 | -0.326381 | NA | Hnf4g, Zfhx4, Pxmp3... |
| 82 | chr3:7509146-7526401 | qA1 | 3 | 0.435681 | NA | Fam164a |
| 83 | chr3:7526402-7684715 | qA1 | 14 | -0.326381 | NA | Fam164a, Il7 |
| 84 | chr3:23598530-24786343 | qA3 | 54 | -0.447973 | NA | |
| 85 | chr3:29078425-29590168 | qA3 | 48 | -0.290790 | NA | |
| 86 | chr3:33423132-36370782 | qA3 - qB | 161 | 0.364580 | NA | Ttc14, Ccdc39, Fxr1... |
| 87 | chr3:39103938-39624118 | qB | 16 | -0.502940 | NA | |
| 88 | chr3:41968881-48042610 | qB - qC | 205 | -0.376212 | NA | Pcdh10, Pabpc4l |
| 89 | chr3:48042611-48847869 | qC | 25 | -0.706654 | NA | 1700018B24Rik |
| 90 | chr3:48847870-50781754 | qC | 99 | -0.376212 | NA | Pcdh18, Slc7a11 |
| 91 | chr3:70282299-78033680 | qE2 - qE3 | 350 | -0.340274 | NA | 2010204N08Rik, Slitrk3, Bche... |
| 92 | chr3:109382158-115114935 | qF3 - qG1 | 273 | -0.288139 | NA | Vav3, Ntng1, Prmt6... |
| 93 | chr3:117639466-119398618 | qG1 | 114 | -0.267789 | NA | Dpyd |
| 94 | chr3:124284465-125772210 | qG1 | 79 | -0.327869 | NA | Ndst4, Ugt8a |

| Event No | Chr | Cytoband | #Probes | Amp/Del | P-value | Annotations |
|---|---|---|---|---|---|---|
| 95 | chr3:135613406-136266549 | qG3 | 43 | -0.372300 | NA | Bank1 |
| 96 | chr3:139363736-141390026 | qH1 | 82 | -0.349296 | NA | B930007M17Rik, Pdha2, Unc5c |
| 97 | chr3:144424171-144490871 | qH2 | 6 | -0.718439 | NA | Clca2, Clca4 |
| 98 | chr3:146923617-148336789 | qH2 - qH3 | 39 | -0.314882 | NA | |
| 99 | chr4:16589785-21505981 | qA2 - qA3 | 279 | -0.299042 | NA | Mmp16, Cngb3, Cpne3... |
| 100 | chr4:22522479-31664269 | qA3 - qA5 | 377 | -0.376693 | NA | F730047E07Rik, Klhl32, Ndufaf4... |
| 101 | chr4:35694352-38779036 | qA5 | 190 | -0.335865 | NA | Lingo2 |
| 102 | chr4:49920677-52425349 | qB1 - qB2 | 75 | -0.355381 | NA | Cylc2 |
| 103 | chr4:62164667-62182581 | qB3 | 3 | 1.644301 | NA | Alad |
| 104 | chr4:63714708-80409927 | qC1 - qC3 | 817 | -0.324688 | NA | Pappa, Astn2, Trim32... |
| 105 | chr4:89889641-93258506 | qC5 | 115 | -0.314010 | NA | Zfp352, Elavl2, Tusc1 |
| 106 | chr4:96155813-97094906 | qC5 - qC6 | 48 | -0.357298 | NA | Cyp2j8, Cyp2j6, Cyp2j9... |
| 107 | chr4:111726014-111810350 | qD1 | 8 | -2.148940 | NA | Skint4 |
| 108 | chr4:111810351-112147017 | qD1 | 26 | -4.085631 | NA | Skint4, Skint3, Skint9 |
| 109 | chr4:112147018-112202834 | qD1 | 1 | -2.148940 | NA | |
| 110 | chr4:112202835-112273763 | qD1 | 4 | -3.864877 | NA | |
| 111 | chr4:112273764-112289417 | qD1 | 1 | -2.148940 | NA | Skint2 |
| 112 | chr4:112289418-112460896 | qD1 | 13 | -0.636102 | NA | Skint2, Skint10 |
| 113 | chr4:112460897-112496839 | qD1 | 5 | -1.477116 | NA | Skint6 |
| 114 | chr4:112496840-112504314 | qD1 | 1 | -0.636102 | NA | Skint6 |
| 115 | chr4:112504315-112525047 | qD1 | 1 | -2.148940 | NA | Skint6 |
| 116 | chr4:112525048-112564300 | qD1 | 5 | -4.311910 | NA | Skint6 |
| 117 | chr4:112564301-112664999 | qD1 | 6 | -2.148940 | NA | Skint6 |
| 118 | chr4:112665000-112785680 | qD1 | 11 | -4.312820 | NA | Skint6 |
| 119 | chr4:112785681-113251939 | qD1 | 26 | -3.297857 | NA | Skint6 |
| 120 | chr4:113286238-113518070 | qD1 | 13 | -0.735915 | NA | |
| 121 | chr4:113518071-113549532 | qD1 | 3 | -2.923769 | NA | |
| 122 | chr4:113549533-113658273 | qD1 | 6 | -0.735915 | NA | |
| 123 | chr4:120886429-122365647 | qD2.2 | 42 | -0.313040 | NA | Gm12888, Gm12886, Gm12887 |
| 124 | chr4:145018403-145192926 | qE1 | 8 | 1.011855 | NA | Vmn2r-ps14 |
| 125 | chr4:146804312-146935370 | qE1 | 7 | -0.614439 | NA | Gm13152, Gm13154 |
| 126 | chr4:154192251-154237986 | qE2 | 4 | -0.646273 | NA | B230396O12Rik |
| 127 | chr5:6936780-7393355 | qA1 | 44 | -0.431666 | NA | Zfp804b, 4921511H03Rik |
| 128 | chr5:9629608-15684052 | qA1 - qA2 | 256 | -0.326788 | NA | Grm3, Gm6455, Speer1-ps1... |
| 129 | chr5:15684053-15975104 | qA2 | 26 | -0.912313 | NA | Cacna2d1 |
| 130 | chr5:15975105-17756918 | qA2 - qA3 | 95 | -0.326788 | NA | Hgf, Speer4f, Sema3c... |
| 131 | chr5:22360303-22889017 | qA3 | 44 | -0.381943 | NA | Lhfpl3 |
| 132 | chr5:36745322-36771300 | qB3 | 3 | -1.041515 | NA | |
| 133 | chr5:44646131-48958933 | qB3 | 247 | -0.337460 | NA | Ldb2, Qdpr, Lap3... |
| 134 | chr5:48958934-51788154 | qB3 - qC1 | 144 | -0.505875 | NA | Kcnip4, Gpr125 |
| 135 | chr5:51788155-56750552 | qC1 | 236 | -0.337460 | NA | Ppargc1a, Dhx15, Sod3... |
| 136 | chr5:56750553-61534535 | qC1 - qC3.1 | 186 | -0.484351 | NA | Gm8121, Pcdh7 |
| 137 | chr5:61534536-63245794 | qC3.1 | 60 | -0.337460 | NA | G6pd2, Arap2 |
| 138 | chr5:68910011-72352879 | qC3.1 - qC3.2 | 181 | -0.263881 | NA | Kctd8, Yipf7, Guf1... |
| 139 | chr5:78753157-86355637 | qD - qE1 | 298 | -0.381980 | NA | Lphn3, Tecrl, Epha5 |
| 140 | chr5:105142362-105237345 | qE5 | 6 | -0.709908 | NA | |
| 141 | chr5:140230570-140265361 | qG2 | 5 | -0.740265 | NA | Ints1 |
| 142 | chr6:3131098-3407796 | qA1 | 10 | 0.497535 | NA | Samd9l, Hepacam2 |
| 143 | chr6:3407797-9159380 | qA1 | 424 | 0.671151 | NA | Hepacam2, Ccdc132, Calcr... |
| 144 | chr6:9159381-27894005 | qA1 - qA3.2 | 1054 | 0.555761 | NA | Nxph1, Ndufa4, Phf14... |
| 145 | chr6:27894006-28017316 | qA3.2 | 10 | 0.671151 | NA | Grm8 |
| 146 | chr6:28017317-34506915 | qA3.2 - qB1 | 545 | 0.778975 | NA | Grm8, Zfp800, Gcc1... |
| 147 | chr6:34506916-34541940 | qB1 | 3 | -0.437801 | NA | |
| 148 | chr6:34541941-36235264 | qB1 | 108 | 0.778975 | NA | Cald1, Agbl3, Tmem140... |

| Event No | Chr | Cytoband | #Probes | Amp/Del | P-value | Annotations |
|---|---|---|---|---|---|---|
| 149 | chr6:36235265-36824823 | qB1 | 50 | 0.586718 | NA | Chrm2, Ptn, Dgki |
| 150 | chr6:36824824-40461947 | qB1 | 325 | 0.778975 | NA | Dgki, Creb3l2, Akr1d1... |
| 151 | chr6:40461948-41616443 | qB1 - qB2.1 | 94 | 0.671151 | NA | 1700016G05Rik, Olfr461, Olfr460... |
| 152 | chr6:41616444-41948741 | qB2.1 | 28 | 0.422524 | NA | Trpv5, 1700034O15Rik, Kel... |
| 153 | chr6:41948742-44104379 | qB2.1 | 165 | 0.671151 | NA | Sva, Tas2r139, Tas2r144... |
| 154 | chr6:44104380-46691153 | qB2.1 - qB2.3 | 195 | 0.483134 | NA | Cntnap2 |
| 155 | chr6:46691154-47525468 | qB2.3 | 65 | 0.671151 | NA | Cntnap2, Cul1, Ezh2 |
| 156 | chr6:47525469-52841579 | qB2.3 - qB3 | 392 | 0.816817 | NA | Ezh2, Rn4.5s, Pdia4... |
| 157 | chr6:52841580-52959167 | qB3 | 13 | 0.671151 | NA | Jazf1 |
| 158 | chr6:52959168-55745250 | qB3 | 228 | 0.497535 | NA | Jazf1, Creb5, 1200009O22Rik... |
| 159 | chr6:55745251-64261786 | qB3 - qC1 | 605 | 0.318060 | NA | Ccdc129, Gsbs, Pde1c... |
| 160 | chr6:64261787-67829764 | qC1 | 229 | 0.434402 | NA | Grid2, Atoh1, Smarcad1... |
| 161 | chr6:67829765-69953594 | qC1 | 86 | 0.318060 | NA | Rprl1 |
| 162 | chr6:69953595-73408961 | qC1 | 278 | 0.575145 | NA | Rpia, Eif2ak3, 1700011F03Rik... |
| 163 | chr6:73408962-81670376 | qC1 - qC3 | 388 | 0.318060 | NA | 4931417E11Rik, Ctnna2, Lrrtm1... |
| 164 | chr6:81670377-95929197 | qC3 - qD3 | 1169 | 0.497535 | NA | AW146020, Mrpl19, Fam176a... |
| 165 | chr6:95929198-96881726 | qD3 | 83 | 0.281142 | NA | Fam19a1, 1700123L14Rik, Fam19a4 |
| 166 | chr6:96881727-107513770 | qD3 - qE1 | 651 | 0.497535 | NA | Fam19a4, A130022J15Rik, Tmf1... |
| 167 | chr6:107513771-108820964 | qE1 - qE2 | 94 | 0.520587 | NA | Lrrn1, Setmar, Sumf1... |
| 168 | chr6:108820965-128726560 | qE2 - qF3 | 1507 | 0.497535 | NA | Grm7, 1700054K19Rik, Lmcd1... |
| 169 | chr6:128726561-128746179 | qF3 | 3 | -1.141174 | NA | Klrb1c |
| 170 | chr6:128746180-129795762 | qF3 | 93 | 0.497535 | NA | Klrb1b, Clec2i, Clec2g... |
| 171 | chr6:129795763-129861126 | qF3 | 8 | 0.636298 | NA | Klra17, Klra5 |
| 172 | chr6:129861127-135300756 | qF3 - qG1 | 282 | 0.497535 | NA | Klra5, Klra22, Klra15... |
| 173 | chr6:135300757-148030100 | qG1 - qG3 | 950 | 0.414057 | NA | Emp1, Grin2b, E330021D16Rik... |
| 174 | chr6:148030101-149494692 | qG3 | 115 | 0.497535 | NA | Far2, Ergic2, Tmtc1... |
| 175 | chr7:7089757-12330733 | qA1 | 68 | -0.429552 | NA | Zfp418, BC023179, Vmn2r29... |
| 176 | chr7:18441712-18574960 | qA2 | 10 | -0.488279 | NA | Ceacam11 |
| 177 | chr7:37637857-37719766 | qB2 | 3 | -1.306158 | NA | |
| 178 | chr7:47377405-47468404 | qB3 | 4 | -0.978053 | NA | |
| 179 | chr7:54844755-54883844 | qB4 | 3 | 0.917504 | NA | Mrgpra3 |
| 180 | chr7:54920080-57322672 | qB4 - qB5 | 184 | -0.301383 | NA | Mrgpra4, Mrgprx1, Mrgprb5... |
| 181 | chr7:57322673-58101311 | qB5 | 84 | -0.456805 | NA | Nell1, 4933405O20Rik |
| 182 | chr7:58101312-69509710 | qB5 - qC | 641 | -0.301383 | NA | Nell1, Ano5, Slc17a6... |
| 183 | chr7:77735091-79161612 | qD1 | 49 | -0.345022 | NA | |
| 184 | chr7:83564899-85168573 | qD2 - qD3 | 70 | -0.387994 | NA | |
| 185 | chr7:91816857-100582827 | qD3 - qE1 | 679 | -0.259943 | NA | Zfand6, Olfr291, Olfr290... |
| 186 | chr7:100582828-102901936 | qE1 | 85 | -0.422197 | NA | |
| 187 | chr7:102901937-103146283 | qE1 | 7 | -0.259943 | NA | |
| 188 | chr7:110685282-111656561 | qE3 | 92 | -0.315371 | NA | Olfr614, Olfr615, Olfr616... |
| 189 | chr7:111656562-111683983 | qE3 | 3 | -1.148189 | NA | Gm6577 |
| 190 | chr7:111683984-112282633 | qE3 | 48 | -0.315371 | NA | Olfr651, Olfr652, Olfr653... |
| 191 | chr7:131026888-131119092 | qF3 | 5 | -0.749452 | NA | |
| 192 | chr8:3130823-5088880 | qA1.1 | 140 | 0.360986 | NA | Insr, A430078G23Rik, Arhgef18... |
| 193 | chr8:8152850-11316585 | qA1.1 | 221 | 0.353650 | NA | Efnb2, Arglu1, Fam155a... |
| 194 | chr8:11316586-11382229 | qA1.1 | 9 | -1.159926 | NA | Col4a2 |
| 195 | chr8:11382230-11388567 | qA1.1 | 1 | 0.353650 | NA | Col4a2 |
| 196 | chr8:11388568-14374806 | qA1.1 | 250 | 0.493702 | NA | Col4a2, Rab20, Carkd... |
| 197 | chr8:14374807-16530927 | qA1.1 - qA1.2 | 159 | 0.353650 | NA | Dlgap2, Cln8, Arhgef10... |
| 198 | chr8:16530928-16630580 | qA1.2 | 11 | -0.320738 | NA | Csmd1 |
| 199 | chr8:16630581-18568094 | qA1.2 - qA1.3 | 124 | 0.353650 | NA | Csmd1 |

| Event No | Chr | Cytoband | #Probes | Amp/Del | P-value | Annotations |
|---|---|---|---|---|---|---|
| 200 | chr8:18568095-18919606 | qA1.3 | 35 | 0.430019 | NA | Mcph1, Angpt2, Agpat5 |
| 201 | chr8:18919607-22148453 | qA1.3 - qA2 | 53 | 0.353650 | NA | Xkr5, Defb40, Defb37... |
| 202 | chr8:22148454-22233176 | qA2 | 3 | 1.363272 | NA | Defa21, Defa-rs7, Defa23... |
| 203 | chr8:22233177-25961591 | qA2 | 264 | 0.353650 | NA | Defa22, Defa-rs7, Defa23... |
| 204 | chr8:25961592-27189553 | qA2 | 107 | 0.496848 | NA | Adam32, Adam9, Tm2d2... |
| 205 | chr8:27189554-28804365 | qA2 | 95 | 0.353650 | NA | Hook3, Rnf170, Thap1... |
| 206 | chr8:35968865-36427098 | qA4 | 22 | 0.498302 | NA | Tnks |
| 207 | chr8:40187843-40274870 | qA4 | 6 | -0.702161 | NA | Tusc3 |
| 208 | chr8:41436380-42014895 | qA4 | 52 | 0.403250 | NA | Efha2, Zdhhc2, Cnot7... |
| 209 | chr8:43127913-43149700 | qA4 | 3 | -1.238168 | NA | |
| 210 | chr8:45984820-49694529 | qB1.1 - qB1.2 | 322 | 0.389803 | NA | Fat1, Mtnr1a, F11... |
| 211 | chr8:63020940-64466092 | qB3.1 | 121 | 0.381031 | NA | Aadat, Mfap3l, 2700029M09Rik... |
| 212 | chr8:93508957-93530058 | qC5 | 3 | -0.729472 | NA | Chd9 |
| 213 | chr8:95561100-95630883 | qC5 | 7 | -0.533038 | NA | Gm4976, Gm5158, Es1 |
| 214 | chr8:95949644-98319347 | qC5 - qD1 | 212 | 0.262346 | NA | Ces7, Gnao1, 4930488L21Rik... |
| 215 | chr8:106778160-130303538 | qD3 - qE2 | 1844 | 0.288870 | NA | Cklf, Cmtm2a, Cmtm2b... |
| 216 | chr9:3543177-6342549 | qA1 | 169 | -0.258462 | NA | Gucy1a2, Aasdhppt, Kbtbd3... |
| 217 | chr9:9531013-11730807 | qA1 | 157 | -0.372565 | NA | Cntn5 |
| 218 | chr9:16124571-19375718 | qA2 | 156 | -0.252578 | NA | Fat3, Chordc1, Naalad2... |
| 219 | chr9:27633405-29264679 | qA4 | 162 | -0.310124 | NA | Opcml, Ntm |
| 220 | chr9:62273455-62295798 | qB | 3 | -0.777700 | NA | Coro2b |
| 221 | chr9:80346215-83221489 | qE1 - qE2 | 137 | -0.340179 | NA | Impg1, Htr1b, 4930486G11Rik... |
| 222 | chr9:84405744-84727295 | qE3.1 | 8 | -0.571476 | NA | |
| 223 | chr9:92695995-94437198 | qE3.3 | 50 | -0.331514 | NA | 1190002N15Rik |
| 224 | chr9:95556843-95572710 | qE3.3 | 3 | -1.119205 | NA | Pcolce2 |
| 225 | chr9:101271461-101724441 | qE4 - qF1 | 16 | -0.537492 | NA | |
| 226 | chr9:105390517-105439072 | qF1 | 5 | -0.697153 | NA | Atp2c1 |
| 227 | chr9:117884082-117905398 | qF3 | 3 | -1.068836 | NA | |
| 228 | chr10:14972794-16961662 | qA2 | 67 | -0.316296 | NA | |
| 229 | chr10:45840624-49920158 | qB2 - qB3 | 164 | -0.371452 | NA | Grik2 |
| 230 | chr10:53780358-55184710 | qB3 | 43 | -0.326431 | NA | Man1a |
| 231 | chr10:63752486-65681938 | qB4 - qB5.1 | 109 | -0.300647 | NA | Ctnna3 |
| 232 | chr10:71489416-74071450 | qB5.3 | 165 | -0.267996 | NA | Zwint, Pcdh15 |
| 233 | chr10:88379728-88397518 | qC1 | 3 | -2.136123 | NA | Slc5a8 |
| 234 | chr10:89120497-89257314 | qC2 | 16 | 0.258705 | NA | Scyl2, Actr6, Uhrf1bp1l |
| 235 | chr10:90492952-90513111 | qC2 | 3 | -1.101546 | NA | Apaf1 |
| 236 | chr10:91716385-92481070 | qC2 | 35 | -0.361899 | NA | Nedd1 |
| 237 | chr10:93709104-93752466 | qC2 | 4 | -1.106289 | NA | |
| 238 | chr10:97058389-106780747 | qC3 - qD1 | 516 | -0.326835 | NA | Kera, Epyc, 4921510H08Rik... |
| 239 | chr10:112396193-114354378 | qD2 | 89 | -0.295461 | NA | Trhde |
| 240 | chr10:122860752-124666967 | qD2 - qD3 | 85 | -0.375640 | NA | Fam19a2, 4930503E24Rik, Slc16a7 |
| 241 | chr11:3107005-3567716 | qA1 | 41 | 0.654618 | NA | Eif4enif1, Drg1, Patz1... |
| 242 | chr11:9255738-9522194 | qA1 | 25 | -0.269382 | NA | Abca13 |
| 243 | chr11:9556391-9638456 | qA1 | 6 | -0.871609 | NA | Abca13 |
| 244 | chr11:12559224-15881378 | qA1 - qA2 | 121 | -0.252464 | NA | Pom121l12 |
| 245 | chr11:15932992-17988207 | qA2 - qA3.1 | 110 | 0.320238 | NA | Vstm2a, Sec61g, Egfr... |
| 246 | chr11:18057980-21723783 | qA3.1 | 208 | 0.774421 | NA | Meis1, Spred2, Actr2... |
| 247 | chr11:21723784-21981226 | qA3.1 - qA3.2 | 19 | 0.408566 | NA | AV249152, Otx1, Ehbp1 |
| 248 | chr11:27574653-27632740 | qA3.3 | 3 | -3.987885 | NA | |
| 249 | chr11:30540062-30582473 | qA4 | 5 | -1.153551 | NA | Acyp2 |
| 250 | chr11:31049571-31227838 | qA4 | 5 | -1.045742 | NA | |
| 251 | chr11:36092369-36860048 | qA4 - qA5 | 70 | -0.283227 | NA | Odz2 |
| 252 | chr11:36860049-39728807 | qA5 | 86 | -0.444533 | NA | |
| 253 | chr11:39728808-42474020 | qA5 | 141 | -0.283227 | NA | Mat2b, Hmmr, Nudcd2... |

| Event No | Chr | Cytoband | #Probes | Amp/Del | P-value | Annotations |
|---|---|---|---|---|---|---|
| 254 | chr11:57072524-57096542 | qB1.3 | 4 | -0.675585 | NA | Gria1 |
| 255 | chr11:57299896-57333085 | qB1.3 | 4 | 0.667367 | NA | Fam114a2, Mfap3 |
| 256 | chr11:57333086-58533135 | qB1.3 | 100 | 0.384779 | NA | Mfap3, Galnt10, Sap30l... |
| 257 | chr11:58533136-58660441 | qB1.3 | 13 | 0.667367 | NA | Olfr318, Olfr317, Olfr316... |
| 258 | chr11:58660442-62358420 | qB1.3 - qB2 | 341 | 0.927516 | NA | 2810021J22Rik, Zfp39, Butr1... |
| 259 | chr11:62358421-63610764 | qB2 - qB3 | 83 | 0.667367 | NA | Ubb, Trpv2, BC046404... |
| 260 | chr11:63610765-63937028 | qB3 | 30 | 0.971816 | NA | Hs3st3b1, Cox10 |
| 261 | chr11:63937029-64526710 | qB3 | 38 | 0.667367 | NA | Hs3st3a1 |
| 262 | chr11:64526711-67286644 | qB3 | 199 | 0.438770 | NA | Elac2, AU040829, Myocd... |
| 263 | chr11:70989862-71107103 | qB4 | 11 | -3.323278 | NA | Nlrp1b, Nlrp1c |
| 264 | chr11:74184906-74199333 | qB5 | 3 | 1.036847 | NA | Garnl4 |
| 265 | chr11:88256203-88270300 | qC | 3 | -0.829791 | NA | Msi2 |
| 266 | chr11:90590119-92090603 | qD | 48 | -0.256353 | NA | Kif2b |
| 267 | chr11:92435900-92967823 | qD | 16 | 0.473176 | NA | Car10 |
| 268 | chr11:113240579-121797041 | qE2 | 780 | 0.426144 | NA | Slc39a11, Sstr2, Cog1... |
| 269 | chr12:3162616-3246953 | qA1.1 | 5 | 0.422408 | NA | |
| 270 | chr12:3246954-5205868 | qA1.1 | 197 | 0.561061 | NA | Rab10, Kif3c, Asxl2... |
| 271 | chr12:5205869-16354595 | qA1.1 | 530 | 0.422408 | NA | Apob, 1110057K04Rik, Gdf7... |
| 272 | chr12:16354596-16528555 | qA1.1 | 5 | -0.419462 | NA | |
| 273 | chr12:16528556-18231389 | qA1.1 - qA1.2 | 98 | 0.422408 | NA | Lpin1, Ntsr2, Greb1... |
| 274 | chr12:18231390-19146894 | qA1.2 | 4 | 1.134618 | NA | 5730507C01Rik |
| 275 | chr12:19146895-27313077 | qA1.2 - qA2 | 179 | 0.422408 | NA | Ddef2, Itgb1bp1, Cpsf3... |
| 276 | chr12:27313078-27467012 | qA2 | 6 | -0.477471 | NA | |
| 277 | chr12:27467013-31757591 | qA2 | 252 | 0.422408 | NA | Sox11, Allc, Colec11... |
| 278 | chr12:31757592-32901184 | qA2 - qA3 | 105 | 0.565692 | NA | Fam110c, Lamb1-1, Dld... |
| 279 | chr12:32901185-40547727 | qA3 - qB1 | 489 | 0.422408 | NA | 2010109K11Rik, Nampt, Sypl... |
| 280 | chr12:40547728-41934755 | qB1 | 115 | 0.392048 | NA | Arl4a, Scin, Gm889... |
| 281 | chr12:41934756-45282946 | qB1 - qB2 | 141 | 0.422408 | NA | Immp2l, Lrrn3 |
| 282 | chr12:45282947-45726673 | qB2 - qB3 | 46 | 0.500094 | NA | Dnajb9, Pnpla8, Nrcam |
| 283 | chr12:45726674-53805379 | qB3 - qC1 | 394 | 0.422408 | NA | Stxbp6, Foxg1, 3110039M20Rik... |
| 284 | chr12:53805380-57439363 | qC1 | 281 | 0.535484 | NA | Akap6, Npas3, Egln3... |
| 285 | chr12:57439364-66014678 | qC1 | 380 | 0.422408 | NA | Mbip, Nkx2-1, Nkx2-9... |
| 286 | chr12:66014679-66648264 | qC1 - qC2 | 41 | 0.541462 | NA | Gm527, Klhl28, Fam179b... |
| 287 | chr12:66648265-70225157 | qC2 | 173 | 0.422408 | NA | Rpl10l, Mdga2 |
| 288 | chr12:70225158-75264266 | qC2 - qC3 | 411 | 0.533065 | NA | Rps29, Ppil5, Rpl36al... |
| 289 | chr12:75264267-76488660 | qC3 | 76 | 0.333546 | NA | Syt16, Dbpht2, Kcnh5... |
| 290 | chr12:76488661-88813426 | qC3 - qD2 | 1086 | 0.533065 | NA | Rhoj, Gphb5, Ppp2r5e... |
| 291 | chr12:88813427-92240310 | qD2 - qD3 | 232 | 0.422408 | NA | Oog1, BB287469, 100039042... |
| 292 | chr12:92240311-93098347 | qD3 | 85 | 0.577087 | NA | 4930534B04Rik, Tshr, Gtf2a1... |
| 293 | chr12:93098348-99871850 | qD3 - qE | 222 | 0.422408 | NA | Flrt2, 1700019M22Rik, Galc... |
| 294 | chr12:99871851-114676625 | qE - qF1 | 1160 | 0.542274 | NA | Spata7, Ptpn21, Zc3h14... |
| 295 | chr12:114676626-114956321 | qF1 | 20 | 0.876512 | NA | Adam6b, Adam6a |
| 296 | chr12:114956322-115298715 | qF1 | 27 | 0.542274 | NA | |
| 297 | chr12:115298716-116840010 | qF1 - qF2 | 85 | 0.422408 | NA | |
| 298 | chr12:116840011-117047397 | qF2 | 7 | 1.351995 | NA | |
| 299 | chr12:117047398-121241648 | qF2 | 292 | 0.422408 | NA | Zfp386, Vipr2, Wdr60... |
| 300 | chr13:6207199-7685832 | qA1 | 82 | -0.253601 | NA | Pitrm1, Pfkp |
| 301 | chr13:7685833-7844615 | qA1 | 10 | -0.734233 | NA | |
| 302 | chr13:7844616-11664478 | qA1 | 277 | -0.253601 | NA | Adarb2, Wdr37, Idi1... |
| 303 | chr13:15999110-17318673 | qA1 - qA2 | 72 | -0.400671 | NA | Inhba, 5033411D12Rik |
| 304 | chr13:25939124-27806807 | qA3.1 | 95 | -0.422955 | NA | Hdgfl1, Prl, Prl3d1... |
| 305 | chr13:69891394-72004571 | qC1 | 93 | -0.267330 | NA | Med10, Adamts16 |
| 306 | chr13:78471787-89065536 | qC2 - qC3 | 439 | -0.262141 | NA | Arrdc3, Gpr98, Lysmd3... |
| 307 | chr13:89065537-89165484 | qC3 | 13 | -0.750011 | NA | Edil3 |

| Event No | Chr | Cytoband | #Probes | Amp/Del | P-value | Annotations |
|---|---|---|---|---|---|---|
| 308 | chr13:89165485-91142603 | qC3 | 133 | -0.262141 | NA | Edil3, Hapln1, Vcan... |
| 309 | chr13:96261788-96302487 | qD1 | 6 | -0.553436 | NA | F2rl1 |
| 310 | chr13:101149908-101183591 | qD1 | 4 | 0.860586 | NA | Naip1 |
| 311 | chr13:106104950-107447137 | qD1 - qD2.1 | 50 | -0.335013 | NA | Htr1a |
| 312 | chr13:115711622-115732085 | qD2.2 | 3 | -0.970358 | NA | Itga2 |
| 313 | chr13:118388635-118591947 | qD2.3 | 21 | -0.539469 | NA | Hcn1 |
| 314 | chr14:8549772-9925440 | qA1 | 108 | 0.440551 | NA | Flnb, Dnase1l3, Abhd6... |
| 315 | chr14:11468200-11487335 | qA1 | 3 | 0.738788 | NA | Fhit |
| 316 | chr14:20549623-20697618 | qA3 | 15 | 0.348761 | NA | Nid2, 2700060E02Rik, Gng2 |
| 317 | chr14:20697619-23206060 | qA3 | 263 | 0.479588 | NA | Gng2, 1810063B07Rik, Kcnk5... |
| 318 | chr14:23206061-23364371 | qA3 | 16 | 0.348761 | NA | 1700112E06Rik |
| 319 | chr14:23364372-23383481 | qA3 | 3 | -0.691484 | NA | 1700112E06Rik |
| 320 | chr14:23383482-23726436 | qA3 | 39 | 0.348761 | NA | 1700112E06Rik |
| 321 | chr14:23726437-23748305 | qA3 | 4 | -0.804506 | NA | 1700112E06Rik |
| 322 | chr14:23748306-25410667 | qA3 | 146 | 0.348761 | NA | 1700112E06Rik, Kcnma1, Dlg5... |
| 323 | chr14:25410668-28536308 | qA3 | 216 | 0.460733 | NA | Zmiz1, 4931406H21Rik, Ppif... |
| 324 | chr14:28536309-30429579 | qA3 - qB | 189 | 0.352574 | NA | Erc2, Wnt5a, Gm2670... |
| 325 | chr14:30429580-35824786 | qB | 501 | 0.460733 | NA | Cacna2d3, Selk, Actr8... |
| 326 | chr14:35824787-44542852 | qB - qC1 | 358 | 0.348761 | NA | Grid1, 4930596D02Rik, 4930474N05Rik... |
| 327 | chr14:44542853-44579789 | qC1 | 6 | -2.215660 | NA | Ang6 |
| 328 | chr14:44579790-52376882 | qC1 - qC2 | 533 | 0.348761 | NA | Ang6, Ear2, Ear12... |
| 329 | chr14:52376883-52415019 | qC2 | 5 | -0.749682 | NA | Ang4 |
| 330 | chr14:52415020-53088124 | qC2 | 67 | 0.348761 | NA | AY358078, Ear6, Ear7... |
| 331 | chr14:53088125-53111533 | qC2 | 3 | -0.615848 | NA | Olfr1507 |
| 332 | chr14:53111534-54556104 | qC2 | 44 | 0.348761 | NA | |
| 333 | chr14:54556105-54612181 | qC2 | 7 | -0.310851 | NA | |
| 334 | chr14:54612182-54617430 | qC2 | 1 | 0.348761 | NA | |
| 335 | chr14:54617431-80292310 | qC2 - qD3 | 2126 | 0.446452 | NA | Dad1, Abhd4, Olfr49... |
| 336 | chr14:80292311-87092375 | qD3 - qE1 | 236 | 0.348761 | NA | Olfm4, Pcdh17, Diap3 |
| 337 | chr14:87092376-88525938 | qE1 | 95 | 0.271586 | NA | Diap3, Tdrd3 |
| 338 | chr14:88525939-99375328 | qE1 - qE2.2 | 469 | 0.348761 | NA | Pcdh20, Gm5088, Pcdh9... |
| 339 | chr14:99375329-106747870 | qE2.2 - qE2.3 | 462 | 0.434612 | NA | 2410129H14Rik, 6720463M24Rik, Dis3... |
| 340 | chr14:106747871-117974231 | qE2.3 - qE4 | 524 | 0.348761 | NA | Slitrk1, Slitrk6, Slitrk5... |
| 341 | chr14:117974232-123397518 | qE4 - qE5 | 500 | 0.456929 | NA | Gpc6, Dct, Tgds... |
| 342 | chr14:123397519-124750164 | qE5 | 134 | 0.348761 | NA | Nalcn, Itgbl1, Fgf14 |
| 343 | chr15:3114799-3337852 | qA1 | 17 | 0.583793 | NA | Sepp1, Ghr |
| 344 | chr15:3337853-7922908 | qA1 | 296 | 0.471672 | NA | Ghr, Fbxo4, AW549877... |
| 345 | chr15:7922909-8175763 | qA1 | 24 | 0.748369 | NA | Wdr70, Nup155, 2410089E03Rik |
| 346 | chr15:8175764-11581282 | qA1 | 248 | 0.471672 | NA | 2410089E03Rik, Nipbl, Slc1a3... |
| 347 | chr15:11581283-14992021 | qA1 | 167 | 0.583793 | NA | Npr3, Sub1, Zfr... |
| 348 | chr15:14992022-15100240 | qA1 | 4 | 1.228714 | NA | |
| 349 | chr15:15100241-21864377 | qA1 - qA2 | 296 | 0.583793 | NA | Cdh9, Cdh10, Acot10... |
| 350 | chr15:21864378-23073506 | qA2 | 58 | 0.325306 | NA | Cdh18 |
| 351 | chr15:23073507-26005868 | qA2 - qB1 | 154 | 0.583793 | NA | Cdh18, Basp1, Gm5468... |
| 352 | chr15:26005869-27654391 | qB1 | 112 | 0.430821 | NA | March11, Fbxl7, Ank... |
| 353 | chr15:27654392-28290846 | qB1 | 53 | 0.626864 | NA | Trio, Dnahc5 |
| 354 | chr15:28290847-30597827 | qB1 - qB2 | 115 | 0.430821 | NA | Dnahc5, Ctnnd2 |
| 355 | chr15:30597828-31956972 | qB2 - qB3.1 | 98 | 0.583009 | NA | Ctnnd2, Dap, Ankrd33b... |
| 356 | chr15:31956973-35521477 | qB3.1 | 289 | 0.430821 | NA | Tas2r119, Sema5a, Sdc2... |
| 357 | chr15:35521478-35534833 | qB3.1 | 3 | -0.423397 | NA | Vps13b |
| 358 | chr15:35534834-35558378 | qB3.1 | 2 | 0.430821 | NA | Vps13b |
| 359 | chr15:35558379-35590381 | qB3.1 | 3 | 0.583793 | NA | Vps13b |
| 360 | chr15:35590382-35722602 | qB3.1 | 14 | 1.010444 | NA | Vps13b |

| Event No | Chr | Cytoband | #Probes | Amp/Del | P-value | Annotations |
|---|---|---|---|---|---|---|
| 361 | chr15:35722603-36017337 | qB3.1 | 26 | 0.583793 | NA | Vps13b, Cox6c |
| 362 | chr15:36017338-36123015 | qB3.1 | 10 | 1.129461 | NA | Fbxo43, Polr2k, Spag1 |
| 363 | chr15:36123016-36166193 | qB3.1 | 3 | 0.583793 | NA | Spag1 |
| 364 | chr15:36166194-39255557 | qB3.1 | 234 | 1.161813 | NA | Rnf19a, Ankrd46, Snx31... |
| 365 | chr15:39255558-39278355 | qB3.1 | 3 | 0.361411 | NA | Rims2 |
| 366 | chr15:39278356-40443102 | qB3.1 | 70 | 1.161813 | NA | Rims2, Tm7sf4, Dpys... |
| 367 | chr15:40443103-40967447 | qB3.1 | 51 | 0.895704 | NA | Zfpm2 |
| 368 | chr15:40967448-40995453 | qB3.1 | 1 | 1.161813 | NA | |
| 369 | chr15:40995454-41066930 | qB3.1 | 1 | 0.583793 | NA | |
| 370 | chr15:41066931-42021094 | qB3.1 | 62 | 0.797420 | NA | Oxr1, Abra |
| 371 | chr15:42021095-47406795 | qB3.1 - qB3.3 | 246 | 0.583793 | NA | Angpt1, Rspo2, Eif3e... |
| 372 | chr15:47406796-48844962 | qB3.3 | 128 | 0.454636 | NA | Csmd3 |
| 373 | chr15:48844963-51616643 | qB3.3 - qC | 101 | 0.583793 | NA | Trps1 |
| 374 | chr15:51616644-57547031 | qC - qD1 | 375 | 0.784892 | NA | Eif3h, Utp23, Rad21... |
| 375 | chr15:57547032-60354835 | qD1 | 198 | 0.954299 | NA | Zhx2, Derl1, Wdr67... |
| 376 | chr15:60354836-68932446 | qD1 - qD3 | 486 | 0.784892 | NA | D330050I23Rik, A1bg, Myc... |
| 377 | chr15:68932447-69301490 | qD3 | 11 | 0.583793 | NA | |
| 378 | chr15:69400090-71667211 | qD3 | 94 | -0.360124 | NA | Fam135b, Col22a1 |
| 379 | chr15:93512848-94672512 | qE3 | 74 | 0.568225 | NA | Adamts20, Pus7l, Irak4... |
| 380 | chr15:94672513-95533187 | qE3 - qF1 | 82 | 0.385770 | NA | Tmem117, Nell2, Dbx2 |
| 381 | chr15:95533188-103432826 | qF1 - qF3 | 705 | 0.568225 | NA | Ano6, D030018L15Rik, Gm4371... |
| 382 | chr16:3284525-3735145 | qA1 | 11 | 0.485570 | NA | Olfr161, Mefv |
| 383 | chr16:3735146-5173044 | qA1 | 146 | 0.676551 | NA | Zfp263, Olfr15, Zfp174... |
| 384 | chr16:5173045-6149147 | qA1 | 57 | 0.485570 | NA | Sec14l5, Nagpa, AU021092... |
| 385 | chr16:6149148-6168655 | qA1 | 3 | -0.400778 | NA | A2bp1 |
| 386 | chr16:6168656-10055020 | qA1 | 263 | 0.485570 | NA | A2bp1, Tmem114, BC024814... |
| 387 | chr16:10055021-21417877 | qA1 - qB1 | 900 | 0.570395 | NA | Rpl39l, Atf7ip2, Emp2... |
| 388 | chr16:21417878-22838140 | qB1 | 121 | 0.702217 | NA | Vps8, 2510009E07Rik, Ehhadh... |
| 389 | chr16:22838141-24652773 | qB1 | 143 | 0.570395 | NA | Tbccd1, Dnajb11, Ahsg... |
| 390 | chr16:24652774-28432126 | qB1 - qB2 | 242 | 0.420462 | NA | Lpp, Tprg, Trp63... |
| 391 | chr16:28432127-35493113 | qB2 - qB3 | 598 | 0.570395 | NA | Fgf12, 1600021P15Rik, Hrasls... |
| 392 | chr16:35493114-35550180 | qB3 | 6 | 1.301132 | NA | Sema5b |
| 393 | chr16:35550181-36245660 | qB3 | 65 | 0.570395 | NA | Sema5b, Dirc2, Hspbap1... |
| 394 | chr16:36245661-36336077 | qB3 | 7 | -0.862213 | NA | 2010005H15Rik, Stfa1, BC117090 |
| 395 | chr16:36336078-39028255 | qB3 - qB4 | 247 | 0.570395 | NA | BC100530, Stfa2, Stfa3... |
| 396 | chr16:39028256-39691375 | qB4 | 15 | 0.485570 | NA | |
| 397 | chr16:39691376-41932174 | qB4 | 147 | 0.269949 | NA | Lsamp |
| 398 | chr16:41932175-44835804 | qB4 | 243 | 0.485570 | NA | Lsamp, Gap43, BC002163... |
| 399 | chr16:44835805-44872692 | qB4 | 4 | -0.610137 | NA | Cd200r4, Cd200r2 |
| 400 | chr16:44872693-51117600 | qB4 - qB5 | 370 | 0.485570 | NA | Cd200r2, Cd200r3, Ccdc80... |
| 401 | chr16:51117601-54955230 | qB5 - qC1.1 | 163 | 0.289440 | NA | Cblb, Alcam |
| 402 | chr16:54955231-58719596 | qC1.1 - qC1.2 | 261 | 0.485570 | NA | Zpld1, Nfkbiz, Fam55c... |
| 403 | chr16:58719597-71929163 | qC1.2 - qC3.1 | 673 | 0.332128 | NA | Cldnd1, Olfr172, Olfr173... |
| 404 | chr16:71929164-73447781 | qC3.1 | 86 | 0.604049 | NA | Robo1 |
| 405 | chr16:73447782-77823044 | qC3.1 | 237 | 0.332128 | NA | Robo2, Lipi, Rbm11... |
| 406 | chr16:77823045-78557990 | qC3.1 | 30 | 0.602684 | NA | Cxadr, Btg3, Gm7334... |
| 407 | chr16:78557991-84667318 | qC3.1 - qC3.3 | 225 | 0.332128 | NA | D16Ertd472e, Chodl, Prss7... |
| 408 | chr16:84667319-89465513 | qC3.3 | 310 | 0.485570 | NA | Mrpl39, Jam2, Atp5j... |
| 409 | chr16:89465514-98242899 | qC3.3 - qC4 | 697 | 0.606589 | NA | Krtap8-1, Krtap7-1, Krtap11-1... |
| 410 | chr17:17962623-20196705 | qA3.2 | 155 | -0.285904 | NA | 4930546H06Rik, Has1, Fpr1... |
| 411 | chr17:20196706-20405825 | qA3.2 | 15 | -0.665866 | NA | Fpr-rs6, Vmn2r105, Vmn2r106 |
| 412 | chr17:20405826-21089565 | qA3.2 | 59 | -0.285904 | NA | Vmn2r106, Vmn2r107, Vmn2r108... |

| Event No | Chr | Cytoband | #Probes | Amp/Del | P-value | Annotations |
|---|---|---|---|---|---|---|
| 413 | chr17:27451543-27582091 | qA3.3 | 7 | 0.811291 | NA | Grm4 |
| 414 | chr17:33297519-33319404 | qB1 | 3 | -2.177957 | NA | Olfr55 |
| 415 | chr17:34117148-34226193 | qB1 | 10 | -0.669018 | NA | AA388235, H2-K1, Ring1... |
| 416 | chr17:35387186-35490032 | qB1 | 10 | 0.708256 | NA | Bat1a, H2-D1, LOC547349... |
| 417 | chr17:36860348-36909276 | qB1 | 3 | -1.041324 | NA | |
| 418 | chr17:36989803-37003487 | qB1 | 3 | -0.932040 | NA | Trim26, Trim15 |
| 419 | chr17:38449974-38501757 | qB1 | 5 | -3.019689 | NA | Olfr136 |
| 420 | chr17:38979148-40017193 | qB1 | 22 | -0.400681 | NA | |
| 421 | chr17:40098323-40241053 | qB1 - qB2 | 4 | 0.623693 | NA | |
| 422 | chr17:40403526-40427177 | qB2 | 3 | 0.688018 | NA | |
| 423 | chr17:40458506-42686993 | qB2 - qB3 | 120 | -0.272732 | NA | Gm6084, Crisp2, Rhag... |
| 424 | chr17:52145888-53134014 | qC | 76 | -0.430951 | NA | Kcnh8 |
| 425 | chr17:58712803-62513769 | qD - qE1.1 | 113 | -0.355355 | NA | 2610034M16Rik, Nudt12 |
| 426 | chr17:75853574-78545530 | qE2 | 91 | -0.313298 | NA | Rasgrp3, Fam98a |
| 427 | chr17:89557989-94887164 | qE5 | 246 | -0.280610 | NA | Fshr, Nrxn1, Adcyap1 |
| 428 | chr18:15575902-20007455 | qA1 - qA2 | 182 | -0.276425 | NA | Chst9, Cdh2 |
| 429 | chr18:26022540-30327315 | qA2 - qB1 | 138 | -0.292425 | NA | |
| 430 | chr18:30327316-30434872 | qB1 | 5 | 0.313067 | NA | Pik3c3 |
| 431 | chr18:30434873-31254038 | qB1 | 41 | -0.292425 | NA | Pik3c3, Rit2 |
| 432 | chr18:70848146-73277131 | qE2 | 149 | -0.311607 | NA | Dcc |
| 433 | chr18:73747729-73976871 | qE2 | 24 | 0.638753 | NA | Mex3c, Smad4, Elac1... |
| 434 | chr19:3271730-11022625 | qA | 641 | 0.561504 | NA | Ighmbp2, Mrpl21, Cpt1a... |
| 435 | chr19:11022626-11937866 | qA | 93 | 0.335452 | NA | Ccdc86, Ms4a10, Ms4a15... |
| 436 | chr19:11937867-12069297 | qA | 14 | 0.694347 | NA | Olfr1419, Olfr1420, Patl1... |
| 437 | chr19:12069298-16162409 | qA | 264 | 0.335452 | NA | Olfr1423, Olfr1424, Olfr1425... |
| 438 | chr19:16162410-16542761 | qA | 39 | 0.583938 | NA | Gnaq, Gna14 |
| 439 | chr19:16542762-23112523 | qA - qB | 452 | 0.335452 | NA | Gna14, Vps13a, Foxb2... |
| 440 | chr19:23112524-30749270 | qB - qC1 | 599 | 0.561504 | NA | Klf9, Smc5, Mamdc2... |
| 441 | chr19:30749271-31727978 | qC1 | 104 | 0.429406 | NA | Prkg1, Cstf2t, 8430431K14Rik |
| 442 | chr19:31727979-32932832 | qC1 | 105 | 0.573937 | NA | Prkg1, A1cf, Asah2... |
| 443 | chr19:32932833-35837222 | qC1 - qC2 | 167 | 0.429406 | NA | Rnls, AI747699, Lipf... |
| 444 | chr19:35837223-40105401 | qC2 - qC3 | 328 | 0.561504 | NA | Htr7, Rpp30, Ankrd1... |
| 445 | chr19:40105402-48428122 | qC3 - qD1 | 789 | 0.914169 | NA | Cyp2c54, Cyp2c50, Cyp2c70... |
| 446 | chr19:48428123-48666638 | qD1 | 24 | 0.561504 | NA | Sorcs3 |
| 447 | chr19:48666639-50467493 | qD1 | 87 | 0.719099 | NA | Sorcs3, Sorcs1 |
| 448 | chr19:50467494-52371103 | qD1 - qD2 | 79 | 0.561504 | NA | Sorcs1, Ins1 |
| 449 | chr19:52371104-53393768 | qD2 | 55 | 0.944465 | NA | Xpnpep1, Add3, Mxi1 |
| 450 | chr19:53393769-53401312 | qD2 | 1 | 0.561504 | NA | Mxi1 |
| 451 | chrX:14415255-16905124 | qA1.1 - qA1.2 | 131 | -0.323641 | NA | 4930403L05Rik, Cypt6, Cypt8... |
| 452 | chrX:56064117-65261153 | qA6 - qA7.1 | 392 | -0.302050 | NA | 4930550L24Rik, Fgf13, F9... |
| 453 | chrX:78578186-80644939 | qB - qC1 | 98 | -0.319678 | NA | 4930595M18Rik, Dmd |
| 454 | chrX:80644940-80732798 | qC1 | 11 | -0.776549 | NA | Dmd |
| 455 | chrX:80732799-88491732 | qC1 | 520 | -0.319678 | NA | Dmd, 1600014K23Rik, Map3k7ip3... |
| 456 | chrX:100008370-100023414 | qD | 3 | 0.969478 | NA | |
| 457 | chrX:110500894-129457222 | qE1 - qE3 | 702 | -0.267296 | NA | Dach2, Klhl4, Ube2dnl... |
| 458 | chrY:37653-139871 | qA1 | 8 | -1.640423 | NA | Zfy1 |
| 459 | chrY:139872-253206 | qA1 | 1 | -2.779504 | NA | Ube1y1, Kdm5d |
| 460 | chrY:253207-273823 | qA1 | 4 | -4.764123 | NA | Kdm5d |
| 461 | chrY:273824-346877 | qA1 | 5 | -2.779504 | NA | Kdm5d |
| 462 | chrY:346878-374359 | qA1 | 5 | -4.599603 | NA | Eif2s3y |
| 463 | chrY:374360-434832 | qA1 | 3 | -2.779504 | NA | Tspy-ps, Uty |
| 464 | chrY:434833-520092 | qA1 | 12 | -3.528248 | NA | Uty |
| 465 | chrY:520093-535058 | qA1 | 3 | -1.977978 | NA | Uty |
| 466 | chrY:535059-581689 | qA1 | 6 | -3.528248 | NA | Uty |

| Event No | Chr | Cytoband | #Probes | Amp/Del | P-value | Annotations |
|---|---|---|---|---|---|---|
| 467 | chrY:581690-602081 | qA1 | 3 | -5.756958 | NA | Uty, Ddx3y |
| 468 | chrY:602082-731591 | qA1 | 16 | -3.528248 | NA | Ddx3y, Usp9y |
| 469 | chrY:731592-1021262 | qA1 | 11 | -2.779504 | NA | Usp9y |
| 470 | chrY:1021263-1365243 | qA1 | 12 | -4.320993 | NA | Zfy2 |
| 471 | chrY:1365244-1707093 | qA1 | 16 | -2.779504 | NA | Zfy2 |
| 472 | chrY:1707094-1824601 | qA1 | 4 | -4.690825 | NA | |
| 473 | chrY:1824602-2506308 | qA1 | 6 | -2.779504 | NA | Gm16501, Sry, Rbmy1a1 |
| 474 | chrY:2506309-2546621 | qA1 | 3 | -1.161927 | NA | |
| 475 | chrY:2546622-2635638 | qA1 | 3 | -2.779504 | NA | |

Amp=Amplification
Del=Deletion

### Aberration Summary Report

#### Sample Properties

| | | | |
|---|---|---|---|
| Array type | : | ArraySet | : |
| Red Sample | : XEN R3 | Green Sample | : |

#### Analysis Settings

| | |
|---|---|
| Genome | : mm9 |
| Aberration Algorithm | : ADM-2 |
| Threshold | : 6.0 |
| Fuzzy Zero | : ON |
| GC Correction | : ON |
| Window Size | : 2Kb |
| Centralization (legacy) | : OFF |
| Diploid Peak Centralization | : ON |
| Manually Reassign Peaks | : OFF |
| Combine Replicates (Intra Array) | : ON |
| Feature Level Filter | : glsSaturated = true OR rlsSaturated = true OR glsFeatNonUnifOL = true OR rlsFeatNonUnifOL = true OR LogRatio = 0 |
| Design Level Filter | : Homology = 0 OR IsPseudoautosomal = 1 |
| Genomic Boundary | : OFF |
| Aberration Filter | : ( Minimum Number of Probes for Amplification >= 3 AND Minimum Size (Kb) of Region for Amplification >= 0.0 AND Minimum Avg. Absolute Log Ratio for Amplification >= 0.25 ) OR ( Minimum Number of Probes for Deletion >= 3 AND Minimum Size (Kb) of Region for Deletion >= 0.0 AND Minimum Avg. Absolute Log Ratio for Deletion >= 0.25 ) |

#### Genome Overview

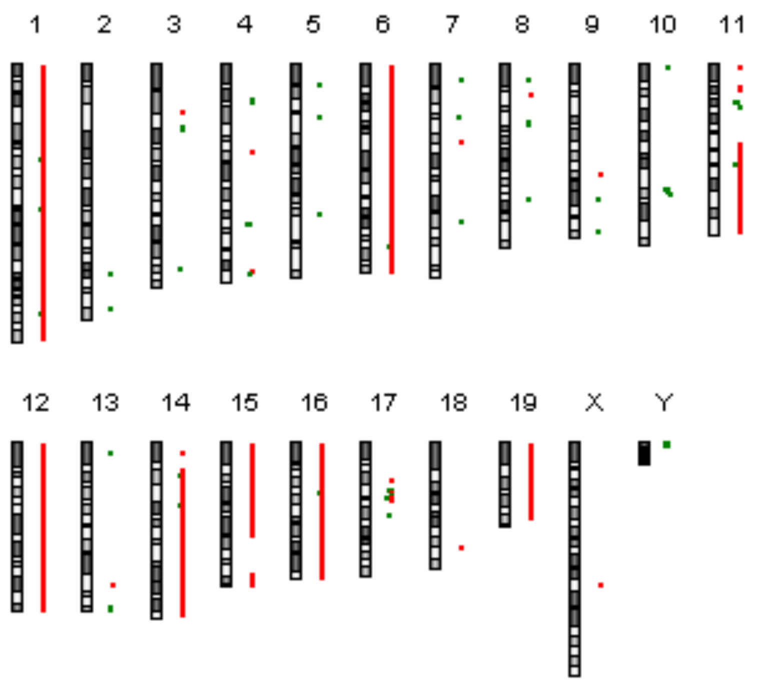

#### Comments

Technician: Date:__/__/__ Supervisor Date:__/__/__

#### Text Summary Report for Sample US81403230_252741111547_S01_CGH_107_Sep09_SureTag_1_3

| Event No | Chr | Cytoband | #Probes | Amp/Del | P-value | Annotations |
|---|---|---|---|---|---|---|
| 1 | chr1:3192257-34606839 | qA1 - qB | 1900 | 0.450556 | NA | Xkr4, Rp1, Sox17... |
| 2 | chr1:34606840-34639821 | qB | 5 | -0.255231 | NA | Tesp2, Fam123c |
| 3 | chr1:34639822-66327175 | qB - qC3 | 2138 | 0.450556 | NA | Fam123c, Arhgef4, Fam168b... |
| 4 | chr1:66327176-66341987 | qC3 | 3 | -1.079504 | NA | Mtap2 |
| 5 | chr1:66341988-67682928 | qC3 | 103 | 0.450556 | NA | Mtap2, Unc80, Rpe... |
| 6 | chr1:67682929-67817197 | qC3 | 4 | -0.831888 | NA | |
| 7 | chr1:67817198-101783951 | qC3 - qE1.1 | 2160 | 0.450556 | NA | Erbb4, Ikzf2, Spag16... |
| 8 | chr1:101783952-101807547 | qE1.1 | 4 | -0.550085 | NA | Cntnap5b |
| 9 | chr1:101807548-141590068 | qE1.1 - qF | 2357 | 0.450556 | NA | Cntnap5b, Cdh20, Rnf152... |
| 10 | chr1:141590069-141678786 | qF | 8 | -0.445433 | NA | EG214403 |
| 11 | chr1:141678787-173457379 | qF - qH3 | 2080 | 0.450556 | NA | BC026782, Cfh, Kcnt2... |
| 12 | chr1:173457380-173490445 | qH3 | 3 | 1.880279 | NA | Itln1, Cd244 |
| 13 | chr1:173490446-175715422 | qH3 | 183 | 0.450556 | NA | Cd244, Ly9, Slamf7... |
| 14 | chr1:175715423-175796142 | qH3 | 6 | -0.895050 | NA | |
| 15 | chr1:175796143-176330988 | qH3 | 41 | 0.450556 | NA | Mnda, Ifi203, Ifi202b... |
| 16 | chr1:176330989-176369910 | qH3 | 6 | -0.368642 | NA | Olfr414 |
| 17 | chr1:176369911-197141706 | qH3 - qH6 | 1483 | 0.450556 | NA | Fmn2, Grem2, Rgs7... |
| 18 | chr2:146839968-146859942 | qG2 | 3 | -0.849333 | NA | Xrn2 |
| 19 | chr2:172762838-172797294 | qH3 | 4 | -1.170266 | NA | Bmp7 |
| 20 | chr3:33423132-36370782 | qA3 - qB | 161 | 0.347478 | NA | Ttc14, Ccdc39, Fxr1... |
| 21 | chr3:43801528-50155490 | qB - qC | 233 | -0.358425 | NA | Pcdh10, Pabpc4l, 1700018B24Rik... |
| 22 | chr3:144424171-144490871 | qH2 | 6 | -0.736920 | NA | Clca2, Clca4 |
| 23 | chr4:24591285-30426687 | qA3 - qA5 | 235 | -0.328707 | NA | Klhl32, Ndufaf4, Gpr63... |
| 24 | chr4:62164667-62182581 | qB3 | 3 | 1.613883 | NA | Alad |
| 25 | chr4:111726014-111810350 | qD1 | 8 | -2.073411 | NA | Skint4 |
| 26 | chr4:111810351-112147017 | qD1 | 26 | -3.998887 | NA | Skint4, Skint3, Skint9 |
| 27 | chr4:112147018-112202834 | qD1 | 1 | -2.073411 | NA | |
| 28 | chr4:112202835-112273763 | qD1 | 4 | -4.113665 | NA | |
| 29 | chr4:112273764-112289417 | qD1 | 1 | -2.073411 | NA | Skint2 |
| 30 | chr4:112289418-112460896 | qD1 | 13 | -0.482256 | NA | Skint2, Skint10 |
| 31 | chr4:112460897-112496839 | qD1 | 5 | -1.408479 | NA | Skint6 |
| 32 | chr4:112496840-112504314 | qD1 | 1 | -0.482256 | NA | Skint6 |
| 33 | chr4:112504315-112525047 | qD1 | 1 | -2.073411 | NA | Skint6 |
| 34 | chr4:112525048-112564300 | qD1 | 5 | -4.224428 | NA | Skint6 |
| 35 | chr4:112564301-112664999 | qD1 | 6 | -2.073411 | NA | Skint6 |
| 36 | chr4:112665000-112688115 | qD1 | 3 | -3.261429 | NA | Skint6 |
| 37 | chr4:112688116-112785680 | qD1 | 8 | -4.203950 | NA | Skint6 |
| 38 | chr4:112785681-113251939 | qD1 | 26 | -3.261429 | NA | Skint6 |
| 39 | chr4:113286238-113574763 | qD1 | 18 | -0.889315 | NA | |
| 40 | chr4:145001092-145192926 | qE1 | 9 | 0.904507 | NA | Vmn2r-ps14 |

| Event No | Chr | Cytoband | #Probes | Amp/Del | P-value | Annotations |
|---|---|---|---|---|---|---|
| 41 | chr4:146804312-146935370 | qE1 | 7 | -0.627181 | NA | Gm13152, Gm13154 |
| 42 | chr5:15684053-15975104 | qA2 | 26 | -0.859304 | NA | Cacna2d1 |
| 43 | chr5:36745322-36771300 | qB3 | 3 | -1.025885 | NA | |
| 44 | chr5:105142362-105237345 | qE5 | 6 | -0.692652 | NA | |
| 45 | chr6:3131098-34506915 | qA1 - qB1 | 2042 | 0.557445 | NA | Samd9l, Hepacam2, Ccdc132... |
| 46 | chr6:34506916-34541940 | qB1 | 3 | -0.359129 | NA | |
| 47 | chr6:34541941-70150127 | qB1 - qC1 | 2594 | 0.557445 | NA | Cald1, Agbl3, Tmem140... |
| 48 | chr6:70150128-70264552 | qC1 | 4 | 1.451905 | NA | |
| 49 | chr6:70264553-128726560 | qC1 - qF3 | 4154 | 0.557445 | NA | Rpia, Eif2ak3, 1700011F03Rik... |
| 50 | chr6:128726561-128746179 | qF3 | 3 | -1.172006 | NA | Klrb1c |
| 51 | chr6:128746180-149494692 | qF3 - qG3 | 1448 | 0.557445 | NA | Klrb1b, Clec2i, Clec2g... |
| 52 | chr7:10751851-12236949 | qA1 | 48 | -0.350564 | NA | Vmn2r52, V1re11, V1re10... |
| 53 | chr7:37637857-37719766 | qB2 | 3 | -1.285762 | NA | |
| 54 | chr7:54844755-54883844 | qB4 | 3 | 1.036325 | NA | Mrgpra3 |
| 55 | chr7:110617416-111918496 | qE3 | 120 | -0.280034 | NA | Olfr608, Olfr609, Olfr610... |
| 56 | chr8:11316586-11382229 | qA1.1 | 9 | -1.151274 | NA | Col4a2 |
| 57 | chr8:22113295-22233176 | qA2 | 5 | 1.147382 | NA | Defa21, Defa-rs7, Defa23... |
| 58 | chr8:40187843-40274870 | qA4 | 6 | -0.652912 | NA | Tusc3 |
| 59 | chr8:43127913-43149700 | qA4 | 3 | -1.086773 | NA | |
| 60 | chr8:95561100-95630883 | qC5 | 7 | -0.481129 | NA | Gm4976, Gm5158, Es1 |
| 61 | chr9:78512136-78574163 | qE1 | 8 | 0.557865 | NA | Cd109 |
| 62 | chr9:95556843-95572710 | qE3.3 | 3 | -1.036121 | NA | Pcolce2 |
| 63 | chr9:117877147-117905398 | qF3 | 4 | -0.867181 | NA | |
| 64 | chr10:3143971-3173641 | qA1 | 5 | -0.732514 | NA | Cnksr3 |
| 65 | chr10:88379728-88397518 | qC1 | 3 | -2.087244 | NA | Slc5a8 |
| 66 | chr10:90492952-90513111 | qC2 | 3 | -1.105483 | NA | Apaf1 |
| 67 | chr10:91368915-92481070 | qC2 | 57 | -0.291140 | NA | Rmst, Nedd1 |
| 68 | chr11:3107005-3578854 | qA1 | 42 | 0.624740 | NA | Eif4enif1, Drg1, Patz1... |
| 69 | chr11:16004019-18032410 | qA2 - qA3.1 | 109 | 0.375723 | NA | Vstm2a, Sec61g, Egfr... |
| 70 | chr11:18032411-21981226 | qA3.1 - qA3.2 | 227 | 0.661650 | NA | Meis1, Spred2, Actr2... |
| 71 | chr11:27574653-27632740 | qA3.3 | 3 | -3.727140 | NA | |
| 72 | chr11:30540062-31602928 | qA4 | 63 | -0.259127 | NA | Acyp2, Psme4, Gpr75... |
| 73 | chr11:57104514-58538776 | qB1.3 | 115 | 0.254116 | NA | Gria1, Fam114a2, Mfap3... |
| 74 | chr11:58538777-62365505 | qB1.3 - qB2 | 354 | 0.818669 | NA | Olfr317, Olfr316, Olfr315... |
| 75 | chr11:62365506-63571284 | qB2 - qB3 | 83 | 0.508715 | NA | Ubb, Trpv2, BC046404... |
| 76 | chr11:63571285-64495531 | qB3 | 68 | 0.818669 | NA | Hs3st3b1, Cox10, Hs3st3a1 |
| 77 | chr11:64495532-64564589 | qB3 | 1 | 0.254116 | NA | |
| 78 | chr11:64564590-67286644 | qB3 | 198 | 0.475439 | NA | Elac2, AU040829, Myocd... |
| 79 | chr11:67286645-70989861 | qB3 - qB4 | 357 | 0.254116 | NA | Gas7, Rcvrn, Glp2r... |
| 80 | chr11:70989862-71107103 | qB4 | 11 | -3.198334 | NA | Nlrp1b, Nlrp1c |
| 81 | chr11:71107104-74184905 | qB4 - qB5 | 255 | 0.254116 | NA | Wscd1, Aipl1, 6720460F02Rik... |
| 82 | chr11:74184906-74199333 | qB5 | 3 | 1.129249 | NA | Garnl4 |
| 83 | chr11:74199334-92175924 | qB5 - qD | 1518 | 0.254116 | NA | Garnl4, E130309D14Rik, 1300001I01Rik... |
| 84 | chr11:92175925-92967823 | qD | 25 | 0.490055 | NA | Car10 |
| 85 | chr11:92967824-110879482 | qD - qE2 | 1638 | 0.254116 | NA | Car10, Utp18, Mbtd1... |
| 86 | chr11:110879483-111518645 | qE2 | 24 | 0.296428 | NA | Kcnj16, Kcnj2 |
| 87 | chr11:111518646-113528440 | qE2 | 107 | 0.254116 | NA | BC006965, Sox9, Slc39a11... |
| 88 | chr11:113528441-121797041 | qE2 | 753 | 0.467528 | NA | D11Wsu99e, D11Wsu47e, Cpsf4l... |
| 89 | chr12:3102541-114676625 | qA1.1 - qF1 | 6713 | 0.465221 | NA | Rab10, Kif3c, Asxl2... |
| 90 | chr12:114676626-115298715 | qF1 | 47 | 0.907466 | NA | Adam6b, Adam6a |
| 91 | chr12:115298716-116840010 | qF1 - qF2 | 85 | 0.465221 | NA | |
| 92 | chr12:116840011-117047397 | qF2 | 7 | 1.443553 | NA | |
| 93 | chr12:117047398-121241648 | qF2 | 292 | 0.465221 | NA | Zfp386, Vipr2, Wdr60... |

| Event No | Chr | Cytoband | #Probes | Amp/Del | P-value | Annotations |
|---|---|---|---|---|---|---|
| 94 | chr13:7593763-7910087 | qA1 | 20 | -0.463247 | NA | |
| 95 | chr13:101110785-101183591 | qD1 | 6 | 0.814464 | NA | Naip1 |
| 96 | chr13:115711622-115732085 | qD2.2 | 3 | -0.927392 | NA | Itga2 |
| 97 | chr13:115825573-115849137 | qD2.2 | 3 | -1.334267 | NA | Itga1 |
| 98 | chr13:118460047-118476029 | qD2.3 | 3 | -1.142198 | NA | Hcn1 |
| 99 | chr14:8549772-10151200 | qA1 | 115 | 0.464821 | NA | Flnb, Dnase1l3, Abhd6... |
| 100 | chr14:20445376-23364371 | qA3 | 298 | 0.404851 | NA | Nid2, 2700060E02Rik, Gng2... |
| 101 | chr14:23364372-23383481 | qA3 | 3 | -0.635439 | NA | 1700112E06Rik |
| 102 | chr14:23383482-23726436 | qA3 | 39 | 0.404851 | NA | 1700112E06Rik |
| 103 | chr14:23726437-23748305 | qA3 | 4 | -0.555034 | NA | 1700112E06Rik |
| 104 | chr14:23748306-44542852 | qA3 - qC1 | 1410 | 0.404851 | NA | 1700112E06Rik, Kcnma1, Dlg5... |
| 105 | chr14:44542853-44579789 | qC1 | 6 | -1.908522 | NA | Ang6 |
| 106 | chr14:44579790-125143137 | qC1 - qE5 | 5241 | 0.404851 | NA | Ang6, Ear2, Ear12... |
| 107 | chr15:3114799-9130920 | qA1 | 405 | 0.654331 | NA | Sepp1, Ghr, Fbxo4... |
| 108 | chr15:9130921-14992021 | qA1 | 346 | 0.428635 | NA | Ugt3a1, Ugt3a2, Capsl... |
| 109 | chr15:14992022-15100240 | qA1 | 4 | 1.331789 | NA | |
| 110 | chr15:15100241-35547960 | qA1 - qB3.1 | 1178 | 0.428635 | NA | Cdh9, Cdh10, Acot10... |
| 111 | chr15:35547961-35590381 | qB3.1 | 4 | 0.654331 | NA | Vps13b |
| 112 | chr15:35590382-35711964 | qB3.1 | 13 | 1.105356 | NA | Vps13b |
| 113 | chr15:35711965-36181161 | qB3.1 | 42 | 0.654331 | NA | Vps13b, Cox6c, Fbxo43... |
| 114 | chr15:36181162-41292216 | qB3.1 | 368 | 1.183582 | NA | Rnf19a, Ankrd46, Snx31... |
| 115 | chr15:41292217-69378937 | qB3.1 - qD3 | 1600 | 0.654331 | NA | Oxr1, Abra, Angpt1... |
| 116 | chr15:93512848-103432826 | qE3 - qF3 | 860 | 0.580047 | NA | Adamts20, Pus7l, Irak4... |
| 117 | chr16:3257712-6149147 | qA1 | 215 | 0.541171 | NA | Olfr161, Mefv, Zfp263... |
| 118 | chr16:6149148-7852010 | qA1 | 142 | 0.279726 | NA | A2bp1 |
| 119 | chr16:7852011-35483724 | qA1 - qB3 | 2127 | 0.541171 | NA | Tmem114, BC024814, Abat... |
| 120 | chr16:35483725-35535857 | qB3 | 6 | 1.197570 | NA | Pdia5 |
| 121 | chr16:35535858-36245660 | qB3 | 66 | 0.541171 | NA | Sema5b, Dirc2, Hspbap1... |
| 122 | chr16:36245661-36336077 | qB3 | 7 | -0.833467 | NA | 2010005H15Rik, Stfa1, BC117090 |
| 123 | chr16:36336078-98263712 | qB3 - qC4 | 3708 | 0.541171 | NA | BC100530, Stfa2, Stfa3... |
| 124 | chr17:27451543-27582091 | qA3.3 | 7 | 0.855195 | NA | Grm4 |
| 125 | chr17:33297519-33319404 | qB1 | 3 | -1.948270 | NA | Olfr55 |
| 126 | chr17:34117148-34494339 | qB1 | 38 | -0.318233 | NA | AA388235, H2-K1, Ring1... |
| 127 | chr17:35387186-35477935 | qB1 | 9 | 0.749227 | NA | Bat1a, H2-D1, LOC547349... |
| 128 | chr17:36860348-37003487 | qB1 | 16 | -0.427689 | NA | H2-M10.5, H2-M10.6, Trim26... |
| 129 | chr17:38449974-38501757 | qB1 | 5 | -2.807249 | NA | Olfr136 |
| 130 | chr17:40082709-40241053 | qB1 - qB2 | 5 | 0.708735 | NA | |
| 131 | chr17:40403526-40427177 | qB2 | 3 | 0.957391 | NA | |
| 132 | chr17:52145888-52211123 | qC | 6 | -0.886257 | NA | |
| 133 | chr18:73767484-73976871 | qE2 | 22 | 0.613737 | NA | Smad4, Elac1, Me2 |
| 134 | chr19:3260956-9785461 | qA | 537 | 0.613913 | NA | Ighmbp2, Mrpl21, Cpt1a... |
| 135 | chr19:9785462-9916174 | qA | 4 | 1.237048 | NA | |
| 136 | chr19:9916175-39989067 | qA - qC3 | 2260 | 0.613913 | NA | Incenp, Fth1, Best1... |
| 137 | chr19:39989068-48428122 | qC3 - qD1 | 794 | 0.919748 | NA | Cyp2c37, Cyp2c54, Cyp2c50... |
| 138 | chr19:48428123-52040633 | qD1 - qD2 | 179 | 0.613913 | NA | Sorcs3, Sorcs1 |
| 139 | chr19:52040634-53393768 | qD2 | 66 | 0.963160 | NA | Ins1, Xpnpep1, Add3... |
| 140 | chr19:53393769-53401312 | qD2 | 1 | 0.613913 | NA | Mxi1 |
| 141 | chrX:100008370-100023414 | qD | 3 | 1.090563 | NA | |
| 142 | chrY:37653-139871 | qA1 | 8 | -1.667212 | NA | Zfy1 |
| 143 | chrY:139872-253206 | qA1 | 1 | -2.747580 | NA | Ube1y1, Kdm5d |
| 144 | chrY:253207-273823 | qA1 | 4 | -5.023287 | NA | Kdm5d |
| 145 | chrY:273824-434832 | qA1 | 13 | -2.747580 | NA | Kdm5d, Eif2s3y, Tspy-ps... |
| 146 | chrY:434833-520092 | qA1 | 12 | -3.666406 | NA | Uty |
| 147 | chrY:520093-535058 | qA1 | 3 | -2.202840 | NA | Uty |

| Event No | Chr | Cytoband | #Probes | Amp/Del | P-value | Annotations |
|---|---|---|---|---|---|---|
| 148 | chrY:535059-581689 | qA1 | 6 | -3.666406 | NA | Uty |
| 149 | chrY:581690-602081 | qA1 | 3 | -6.066073 | NA | Uty, Ddx3y |
| 150 | chrY:602082-697491 | qA1 | 13 | -3.666406 | NA | Ddx3y, Usp9y |
| 151 | chrY:697492-801911 | qA1 | 12 | -2.747580 | NA | Usp9y |
| 152 | chrY:801912-1365243 | qA1 | 14 | -4.269301 | NA | Zfy2 |
| 153 | chrY:1365244-1707093 | qA1 | 16 | -2.747580 | NA | Zfy2 |
| 154 | chrY:1707094-1824601 | qA1 | 4 | -4.654709 | NA | |
| 155 | chrY:1824602-2506308 | qA1 | 6 | -2.747580 | NA | Gm16501, Sry, Rbmy1a1 |
| 156 | chrY:2506309-2546621 | qA1 | 3 | -1.172499 | NA | |
| 157 | chrY:2546622-2635638 | qA1 | 3 | -2.747580 | NA | |

Amp=Amplification
Del=Deletion

### Aberration Summary Report

#### Sample Properties

| | | | |
|---|---|---|---|
| Array type | : | ArraySet | : |
| Red Sample | : 46C WT | Green Sample | : |

#### Analysis Settings

| | |
|---|---|
| Genome | : mm9 |
| Aberration Algorithm | : ADM-2 |
| Threshold | : 6.0 |
| Fuzzy Zero | : ON |
| GC Correction | : ON |
| Window Size | : 2Kb |
| Centralization (legacy) | : OFF |
| Diploid Peak Centralization | : ON |
| Manually Reassign Peaks | : OFF |
| Combine Replicates (Intra Array) | : ON |
| Feature Level Filter | : glsSaturated = true OR rIsSaturated = true OR glsFeatNonUnifOL = true OR rIsFeatNonUnifOL = true OR LogRatio = 0 |
| Design Level Filter | : Homology = 0 OR IsPseudoautosomal = 1 |
| Genomic Boundary | : OFF |
| Aberration Filter | : ( Minimum Number of Probes for Amplification >= 3 AND Minimum Size (Kb) of Region for Amplification >= 0.0 AND Minimum Avg. Absolute Log Ratio for Amplification >= 0.25 ) OR ( Minimum Number of Probes for Deletion >= 3 AND Minimum Size (Kb) of Region for Deletion >= 0.0 AND Minimum Avg. Absolute Log Ratio for Deletion >= 0.25 ) |

#### Genome Overview

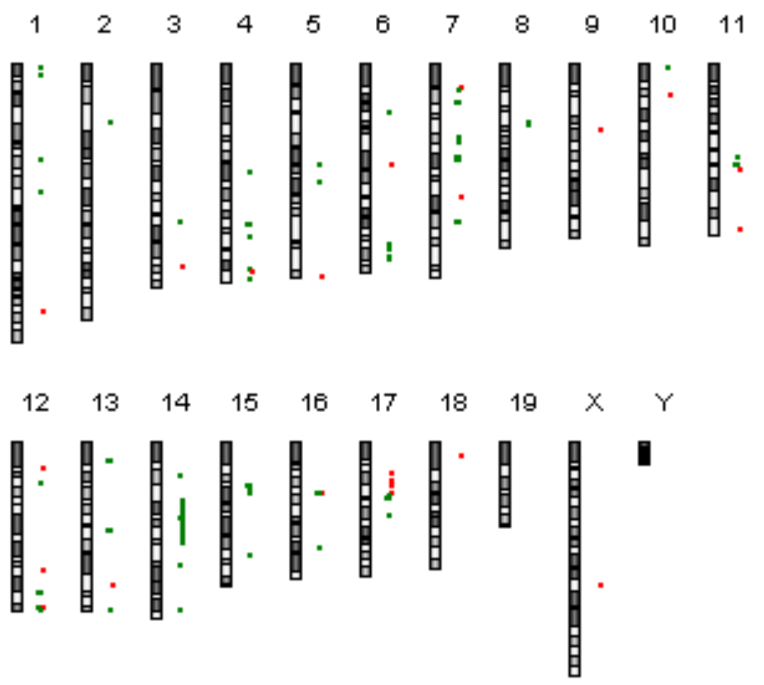

#### Comments

Technician: Date:__/__/__ Supervisor Date:__/__/__

#### Text Summary Report for Sample US81403230_252741111548_S01_CGH_107_Sep09_SureTag_1_2

| Event No | Chr | Cytoband | #Probes | Amp/Del | P-value | Annotations |
|---|---|---|---|---|---|---|
| 1 | chr1:3090501-3154658 | qA1 | 3 | -1.085352 | NA | |
| 2 | chr1:8714929-8749487 | qA1 | 4 | -0.853257 | NA | Sntg1 |
| 3 | chr1:66327176-66341987 | qC3 | 3 | -1.530259 | NA | Mtap2 |
| 4 | chr1:90115633-90193738 | qD | 10 | -1.171071 | NA | Ugt1a10, Ugt1a9, Ugt1a7c... |
| 5 | chr1:173457380-173511412 | qH3 | 5 | 1.875210 | NA | Itln1, Cd244 |
| 6 | chr2:40524011-40546414 | qB | 3 | -1.388911 | NA | Lrp1b |
| 7 | chr3:110878881-110995945 | qF3 | 4 | -1.313088 | NA | |
| 8 | chr3:142264983-142283409 | qH1 | 3 | 0.840475 | NA | Gbp1 |
| 9 | chr4:75901954-75926852 | qC3 | 3 | -1.622616 | NA | Ptprd |
| 10 | chr4:111726014-111819114 | qD1 | 9 | -2.285886 | NA | Skint4 |
| 11 | chr4:111819115-112147017 | qD1 | 25 | -3.985556 | NA | Skint4, Skint3, Skint9 |
| 12 | chr4:112147018-112202834 | qD1 | 1 | -2.285886 | NA | |
| 13 | chr4:112202835-112273763 | qD1 | 4 | -3.948666 | NA | |
| 14 | chr4:112273764-112289417 | qD1 | 1 | -2.285886 | NA | Skint2 |
| 15 | chr4:112289418-112460896 | qD1 | 13 | -0.780540 | NA | Skint2, Skint10 |
| 16 | chr4:112460897-112480643 | qD1 | 3 | -2.267761 | NA | Skint6 |
| 17 | chr4:112480644-112504314 | qD1 | 3 | -0.780540 | NA | Skint6 |
| 18 | chr4:112504315-112525047 | qD1 | 1 | -2.285886 | NA | Skint6 |
| 19 | chr4:112525048-112564300 | qD1 | 5 | -4.627134 | NA | Skint6 |
| 20 | chr4:112564301-112653307 | qD1 | 5 | -2.285886 | NA | Skint6 |
| 21 | chr4:112653308-112785680 | qD1 | 12 | -4.193987 | NA | Skint6 |
| 22 | chr4:112785681-112821288 | qD1 | 3 | -2.285886 | NA | Skint6 |
| 23 | chr4:112821289-112915923 | qD1 | 9 | -3.759207 | NA | Skint6 |
| 24 | chr4:112915924-112954400 | qD1 | 2 | -2.285886 | NA | Skint6 |
| 25 | chr4:112954401-113251939 | qD1 | 12 | -3.593805 | NA | Skint6 |
| 26 | chr4:113266186-113574763 | qD1 | 19 | -0.855060 | NA | |
| 27 | chr4:121649423-121767386 | qD2.2 | 5 | -1.074315 | NA | |
| 28 | chr4:144333817-144357834 | qE1 | 3 | -1.844878 | NA | |
| 29 | chr4:145001092-146912674 | qE1 | 22 | 0.912865 | NA | Vmn2r-ps14, Gm13242, Gm13251... |
| 30 | chr4:150448833-150467116 | qE2 | 3 | -1.028412 | NA | Camta1 |
| 31 | chr5:69891606-69920539 | qC3.1 | 4 | -1.080189 | NA | Yipf7 |
| 32 | chr5:83197619-83285960 | qE1 | 3 | -1.153869 | NA | |
| 33 | chr5:149852342-150181232 | qG3 | 29 | 0.565843 | NA | Hmgb1, Gm8615, Uspl1... |
| 34 | chr6:34506916-34541940 | qB1 | 3 | -1.310004 | NA | |
| 35 | chr6:70150128-70264552 | qC1 | 4 | 0.998098 | NA | |
| 36 | chr6:125808028-125822406 | qF3 | 3 | -0.962783 | NA | Ano2 |
| 37 | chr6:128726561-128746179 | qF3 | 3 | -1.540983 | NA | Klrb1c |
| 38 | chr6:130212984-130317661 | qF3 | 5 | -1.124837 | NA | Klra22, Klra15, Klra12... |
| 39 | chr6:136008799-136029331 | qG1 | 3 | -1.091817 | NA | Grin2b |
| 40 | chr6:137732018-137745184 | qG1 | 3 | -1.038703 | NA | Dera |

| Event No | Chr | Cytoband | #Probes | Amp/Del | P-value | Annotations |
|---|---|---|---|---|---|---|
| 41 | chr7:16481149-16495981 | qA2 | 3 | 0.918540 | NA | Crxos1, Egam-1c |
| 42 | chr7:18382929-18617208 | qA2 | 18 | -0.569678 | NA | Ceacam14, Ceacam11, Ceacam13 |
| 43 | chr7:26790284-26981899 | qA3 | 8 | -2.877568 | NA | Cyp2b13, Cyp2b9 |
| 44 | chr7:52197905-52217244 | qB4 | 3 | -1.023978 | NA | Tsks, Cpt1c |
| 45 | chr7:54758476-54961406 | qB4 | 9 | -1.692609 | NA | Mrgpra3 |
| 46 | chr7:66210072-66237278 | qC | 3 | -2.483995 | NA | |
| 47 | chr7:67364519-67731725 | qC | 14 | -2.336002 | NA | |
| 48 | chr7:93085633-93155866 | qD3 | 6 | 0.906114 | NA | Vmn2r74 |
| 49 | chr7:110667234-110709040 | qE3 | 5 | -1.433732 | NA | Olfr614, Olfr615 |
| 50 | chr7:110844593-110877929 | qE3 | 5 | -1.180215 | NA | Olfr627 |
| 51 | chr7:111430424-111459364 | qE3 | 3 | -3.132178 | NA | Trim12 |
| 52 | chr7:111656562-111683983 | qE3 | 3 | -4.864336 | NA | Gm6577 |
| 53 | chr8:40187843-40274870 | qA4 | 6 | -1.024480 | NA | Tusc3 |
| 54 | chr8:43127913-43149700 | qA4 | 3 | -1.508005 | NA | |
| 55 | chr9:46697499-46906183 | qA5.3 | 9 | 0.890701 | NA | |
| 56 | chr10:3143971-3173641 | qA1 | 5 | -0.833959 | NA | Cnksr3 |
| 57 | chr10:21998927-22072759 | qA3 | 9 | 1.198978 | NA | Raet1a, Raet1b, Raet1c |
| 58 | chr11:64967894-64995611 | qB3 | 3 | -1.389859 | NA | AU040829, Myocd |
| 59 | chr11:70989862-71107103 | qB4 | 11 | -4.621536 | NA | Nlrp1b, Nlrp1c |
| 60 | chr11:74184906-74199333 | qB5 | 3 | 1.143538 | NA | Garnl4 |
| 61 | chr11:116606695-116626075 | qE2 | 3 | 2.337632 | NA | BC018473 |
| 62 | chr12:18231390-18294097 | qA1.2 | 3 | 0.776790 | NA | |
| 63 | chr12:28429347-28483451 | qA2 | 3 | -1.971665 | NA | |
| 64 | chr12:89456049-89496285 | qD3 | 3 | 0.718055 | NA | |
| 65 | chr12:104954787-105048427 | qE | 1 | -1.345111 | NA | Serpina1b, Serpina1d |
| 66 | chr12:105048428-105184867 | qE | 4 | -3.309873 | NA | Serpina1a, Serpina1c |
| 67 | chr12:105184868-105263855 | qE | 9 | -1.345111 | NA | Serpina1e, Serpina11, Serpina9 |
| 68 | chr12:115580565-115880290 | qF2 | 30 | 0.670437 | NA | |
| 69 | chr12:116213616-116261498 | qF2 | 6 | -1.108799 | NA | |
| 70 | chr12:116271371-116291089 | qF2 | 3 | -6.528841 | NA | |
| 71 | chr12:116515691-116624944 | qF2 | 3 | -1.613819 | NA | |
| 72 | chr12:116840011-117047397 | qF2 | 7 | 1.336554 | NA | |
| 73 | chr12:117198164-117274793 | qF2 | 5 | -1.853296 | NA | |
| 74 | chr13:12690823-12723384 | qA1 | 3 | -2.121752 | NA | Ero1lb, Gpr137b-ps |
| 75 | chr13:61743712-62048006 | qB3 | 7 | -2.311447 | NA | |
| 76 | chr13:101053361-101110842 | qD1 | 4 | 1.383961 | NA | Birc1f, Naip7 |
| 77 | chr13:118460047-118476029 | qD2.3 | 3 | -1.415279 | NA | Hcn1 |
| 78 | chr14:23364372-23383481 | qA3 | 3 | -1.368270 | NA | 1700112E06Rik |
| 79 | chr14:23726437-23748305 | qA3 | 4 | -1.239921 | NA | 1700112E06Rik |
| 80 | chr14:40986423-52376882 | qB - qC2 | 595 | -0.457839 | NA | Sh2d4b, Tspan14, 5730469M10Rik... |
| 81 | chr14:52376883-52409033 | qC2 | 4 | -1.403499 | NA | Ang4 |
| 82 | chr14:52409034-73574009 | qC2 - qD3 | 1691 | -0.457839 | NA | AY358078, Ear6, Ear7... |
| 83 | chr14:86407031-86441138 | qE1 | 3 | -1.399797 | NA | |
| 84 | chr14:118170191-118234161 | qE4 | 9 | -0.841844 | NA | Gpc6 |
| 85 | chr15:30446329-30461588 | qB2 | 3 | -2.678653 | NA | Ctnnd2 |
| 86 | chr15:32304625-32359969 | qB3.1 | 6 | -0.989429 | NA | Sema5a |
| 87 | chr15:36619040-36678505 | qB3.1 | 3 | -1.323933 | NA | |
| 88 | chr15:78554311-78592285 | qE1 | 5 | -0.755283 | NA | Mfng |
| 89 | chr16:35483725-35550180 | qB3 | 7 | 0.992612 | NA | Pdia5, Sema5b |
| 90 | chr16:36245661-36319122 | qB3 | 6 | -2.793851 | NA | 2010005H15Rik, Stfa1 |
| 91 | chr16:74549019-74655334 | qC3.1 | 3 | -1.891430 | NA | |
| 92 | chr17:21393034-21406886 | qA3.2 | 3 | 1.009189 | NA | V1rf2 |
| 93 | chr17:22675168-22785051 | qA3.3 | 11 | 0.873075 | NA | Vmn2r111, Vmn2r112 |

| Event No | Chr | Cytoband | #Probes | Amp/Del | P-value | Annotations |
|---|---|---|---|---|---|---|
| 94 | chr17:27390315-27558396 | qA3.3 | 5 | 0.830453 | NA | |
| 95 | chr17:30586287-31049473 | qA3.3 | 50 | 1.021757 | NA | Btbd9, Glo1, Dnahc8... |
| 96 | chr17:36199712-36245963 | qB1 | 4 | 0.943597 | NA | H2-Bl |
| 97 | chr17:36860348-36909276 | qB1 | 3 | -0.954666 | NA | |
| 98 | chr17:40098323-40241053 | qB1 - qB2 | 4 | -3.530537 | NA | |
| 99 | chr17:52145888-52211123 | qC | 6 | -0.804702 | NA | |
| 100 | chr18:9795777-10034659 | qA1 | 20 | 0.869283 | NA | Colec12, Thoc1, Usp14 |
| 101 | chrX:100008370-100023414 | qD | 3 | 1.038788 | NA | |

Amp=Amplification
Del=Deletion
